# Inorganic salts and compatible solutes help mesophilic bacteria inhabit the high temperature waters of a Trans-Himalayan sulfur-borax spring

**DOI:** 10.1101/678680

**Authors:** Nibendu Mondal, Chayan Roy, Aditya Peketi, Masrure Alam, Tarunendu Mapder, Subhrangshu Mandal, Svetlana Fernandes, Sabyasachi Bhattacharya, Moidu Jameela Rameez, Prabir Kumar Haldar, Samida Prabhakar Volvoikar, Nilanjana Nandi, Tannisha Bhattacharya, Aninda Mazumdar, Ranadhir Chakraborty, Wriddhiman Ghosh

**Author notes:** Department of Biological Sciences, Aliah University, IIA/27 New Town, Kolkata - 700160, India. Equal contributions (Nibendu Mondal, Chayan Roy). **Correspondence: Wriddhiman Ghosh**, **Email:**.

## Abstract

While geographically-/geologically-distinct hot springs harbor different levels of microbial diversity, some of them encompass several such taxa which have no strain reported for laboratory growth at >45°C. We, therefore, hypothesized that native geomicrobial factors could be potent determinants of the microbial habitability of hot spring environments. To test this hypothesis, aquatic microbial communities were revealed metataxonomically, and considered in the context of spring-water chemistry, along the 85-14°C hydrothermal gradient of a sulfur-boron spring named *Lotus Pond* located at 4,436 m, within the Puga geothermal area of the Indian Trans-Himalayan region of Ladakh. Water samples were studied from four distinct sites along *Lotus Pond*’s spring-water transit from the vent to an adjacent river called *Rulang*. Insinuations obtained from geomicrobiological data were tested via pure-culture growth experiments in habitat-inspired media. Microbial diversities were found to be high at all the sample-sites; majority of the genera identified at the 70-85°C sites were found to have no report of laboratory growth at >45°C; concurrently, these sample-sites had high concentrations of the kosmotropic solutes boron, lithium, sodium, sulfide, thiosulfate and sulfate, which are known to biophysically stabilize macromolecules. Based on the universal thermodynamic status of these solutes, we conjectured that they may be instrumental in helping mesophiles withstand high *in situ* temperatures. Corroboratively, growth experiments with a mesophilic, 80°C-isolate, *Paracoccus* SMMA_5 showed that at 50°C and 70°C, depending on the incubation-time, lithium/boron/sulfate/sodium/glycine-betaine either increases the number of colony-forming units present in the culture or arrests decline of the same. Incubations at 70°C, followed by fluorescein diacetate staining and flow cytometry, showed that these solutes keep more cells under viable condition than in ready-to-divide state. We concluded that kosmotropes and compatible solutes help mesophiles overcome the chaotropic effects of heat by augmenting such indigenous, entropy-minimizing biophysical mechanisms that apparently trade-off cell division for cell viability.

## 1. Introduction

The association of typically thermophilic/hyperthermophilic microorganisms with hydrothermal habitats is axiomatic (Pace, 1997; Schwartzman and Lineweaver, 2004; Martin et al., 2008). Consequently, our knowledge on thermal adaptation is based on either the hyperthermophiles (often referred to as true thermophiles) that grow in the laboratory exclusively at ≥ 80°C (Kristjansson and Stetter, 1992; Stetter, 1999; Vieille and Zeikus, 2001; Berezovsky and Shakhnovich, 2005) or the facultative/moderate thermophiles that grow *in vitro* between 30°C and 80°C (Moreira et al., 2000; Alves et al., 2003; Rainey et al., 2003; Endo et al., 2006; Shih and Pan, 2011; Goh et al., 2014). However, mesophilic microbial taxa, though unexpected in high-temperature environments, are not alien to hot spring habitats (Baker et al., 2001). Several high-throughput DNA-sequencing-based explorations, in the recent times, have revealed that many high-temperature geothermal microbial communities encompass rich diversities of such taxa which have no members reported for laboratory growth at > 45°C (Jiménez et al., 2012; Wemheuer et al., 2013; Chan et al., 2015; Ghosh et al., 2015; Menzel et al., 2015; Roy et al., 2016). Furthermore, when the findings of a number of such culture-independent studies were collated, it was evident that temperature-wise-similar waters of hot spring systems located in discrete geographical/geological areas harbor different levels of microbial diversity (see Fig. 1; Supplementary Table S1 and references therein). We, therefore, hypothesized that some unknown chemical constituents of the spring-waters, and/or microbial factors present *in situ*, could be instrumental in enhancing the habitability of certain hydrothermal ecosystems, thereby facilitating the survival of mesophilic microorganisms in those environments. Notably, the plausibility of environmental/geochemical factors (other than only temperature, pH and dissolved O_2_) determining the structures and functions of hydrothermal microbial communities has also been envisaged in a few previous publications (Xie et al., 2011; Alsop et al., 2014; Cowan et al., 2015). Furthermore, environment-aided survival of mesophiles in high temperature habitats seems feasible in the light of the fact that researchers have found no special molecular/structural biological feature in the biomacromolecules of thermophiles/hyperthermophiles that can be exclusively attributed to their high temperature adaptation. In other words, all the biophysical features which have thus far been attributed to the high-temperature adaptation of thermophiles/hyperthermophiles (via *in vitro* studies) - for instance specialized topologies and functions of proteins, nucleic acids and membrane lipids (Kumar et al., 2000; Dekker et al., 2003, Zeldovich et al., 2007; Koga, 2012; Wang et al., 2014), protein-stabilizing adaptive mutations (Fields, 2001), and presence of compatible solutes (Welsh, 2000; Fields, 2001) - are present in some mesophile or the other (Fields, 2001; Vieille and Zeikus, 2001; Hallsworth et al., 2003; Rudolph et al., 2010; Dibrova et al., 2014; Ezemaduka et al., 2014; de Lima Alves et al., 2015; Hamerly et al., 2015; Jing et al., 2017; Pucci and Rooman, 2017).

**Fig. 1.**
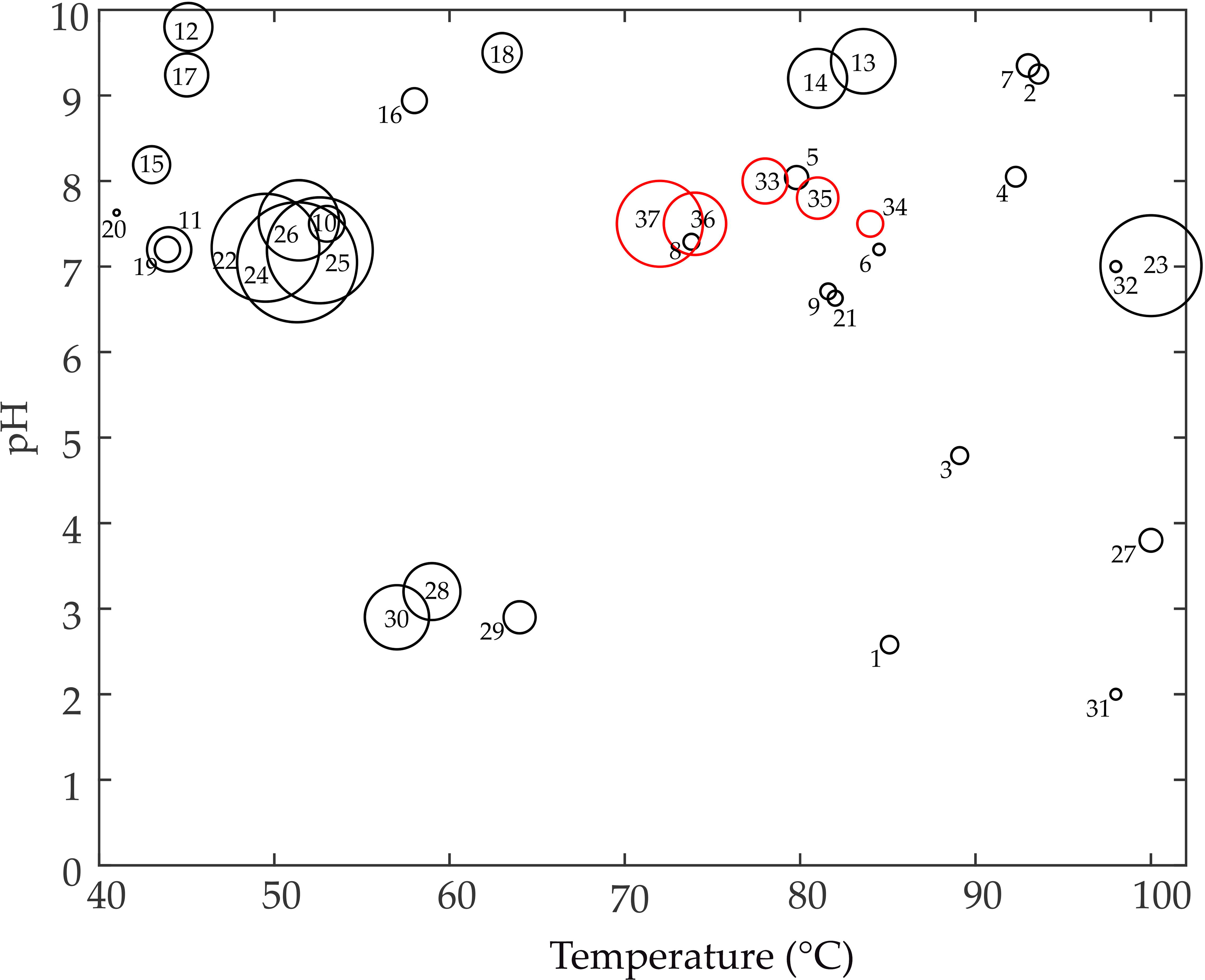
Bubble plot comparing the *in situ* temperature, pH, and OTU-count of the various *Lotus Pond* samples with the similar parameters reported for other geographically-discrete terrestrial hot springs. Temperature is plotted along the X-axis, pH along the Y-axis, while individual springs are represented by circles numbered 1 through 37. Size of the circles is proportional to the OTU-count of the springs: the smallest circle, numbered 20, represents the Eiland hot spring (Limpopo Province of South Africa) from where 6 OTUs were reported; the largest circle, numbered 24, represents the Garbanabra hot spring (Eritrea) from where 2354 OTUs were reported. Circles representing the *Lotus Pond* sample-sites are in red. Number code for springs: (1) Diretiyanqu, China; (2) Jiemeiquan - Sisters Spring, China; (3) Zhenzhuquan – Pearl Spring, China; (4) Huitaijing – Pregnancy Spring, China; (5) Shuirebaozha - Hydrothermal Explosion, China; (6) Dagunguo – Great Boiling Pot, China; (7) Gumingquan– Drum Beating Spring, China; (8) Gongxiaoshe – Co-op Hotel (side), China; (9) Jinze - Golden Pond Motel, China; (10) Jermuk, Southeast Armenia; (11) Arzakan, Southeast Armenia; (12) Lake Magadi, Hot spring 1, Kenya; (13) Lake Magadi, Hot spring 2, Kenya; (14) Lake Magadi, Hot spring 3, Kenya; (15) Mqhephu hot spring, Limpopo Province of South Africa; (16) Tshipise hot spring, Limpopo Province of South Africa; (17) Sagole hot spring, Limpopo Province of South Africa; (18) Siloam hot spring, Limpopo Province of South Africa; (19) Soutinig hot spring, Limpopo Province of South Africa; (20) Eiland hot spring, Limpopo Province of South Africa; (21) Great Boiling Spring 1002W, Great Basin, USA; (22) Akwar, Eritrea; (23) Elegedi, Eritrea; (24) Garbanabra, Eritrea; (25) Gelti, Eritrea; (26) Maiwooi, Eritrea; (27) Icelandic hot springs P0, Iceland; (28) Icelandic hot springs P1, Iceland; (29) Icelandic hot springs P2, Iceland; (30) Icelandic hot springs P3, Iceland; (31) Leirgerdur, Iceland; (32) Hrifla spring, Iceland; (33) Lotus Pond HTP, as explored at 17:30 h of 23.07.2013; (34) Lotus Pond HTP, as explored at 8:30 h of 20.10.2014; (35) Lotus Pond HTP, as explored at 14:30 h of 20.10.2014; Lotus Pond MTP, as explored at 8:30 h of 20.10.2014; (37) Lotus Pond MTP, as explored at 14:30 h of 20.10.2014. Refer to Additional file 1: Tables S1 for the references from where all these information were sourced.

In order to search for potential *in situ* geochemical/microbiological factors that may be correlated with the high diversity of a hot spring system, metataxonomic composition of the aquatic microbial communities and chemical properties of the spring-water were explored along the 85-14°C hydrothermal gradient of a sulfur-boron spring named *Lotus Pond* (Ghosh et al., 2012), the vent-water of which was previously indicated to be microbial-diversity-rich, based on a single sample survey (Roy et al., 2016). *Lotus Pond* is located in the Puga geothermal area of Eastern Ladakh, within the Trans-Himalayan region of India, at an altitude of 4,436 m, where water boils at 85°C (Garrett, 1998). The pH-neutral discharge of *Lotus Pond*, like all other hot springs of the Puga valley area, is relatively poor in salts of silicon but rich in those of boron and sulfur (Garrett 1998; Rai 2001; Ghosh et al., 2012). This unique geochemical feature apparently distinguishes the Puga hot springs from other microbiologically well-studied terrestrial hot spring systems of the world (Jones et al., 2000; Skirnisdottir et al., 2000; Johnson et al., 2001; Johnson et al., 2003; Hetzer et al., 2007; Roeselers et al., 2007; Owen et al., 2008; Huang et al., 2013; Ghosh et al., 2015). Since pH-neutral hot springs rich in various boron and sulfur species, but poor in terms of silicon compounds, are extremely infrequent on Earth, their biogeochemistry is completely unexplored. We therefore had a notional expectation that revelation of the geomicrobiological dynamics of *Lotus Pond* would afford some novel insight into the potential environmental drivers of microbial survival in hydrothermal habitats. Delineation of *Lotus Pond*’s aquatic microbiome, followed by elucidation of the system’s physicochemical constraints and opportunities in the light of the biophysical controls of ecosystem maintenance in other high-entropy habitats, did implicate that certain *in situ* inorganic salts and compatible solutes could be instrumental in supporting high diversity of mesophiles in *Lotus Pond*’s waters. This hypothesis was subsequently validated by conducting laboratory growth experiments in habitat-inspired media and culture-conditions, with a mesophilic strain of *Paracoccus*, isolated from an 80°C water-sample of *Lotus Pond*.

## 2. Materials and methods

### 2.1. Geography and geology of the study area

The *Lotus Pond* hot spring is located (at GPS coordinates: 33° 13’ 46.3” N and 78° 21’ 19.7” E) in the eastern flank of the Puga valley of Eastern Ladakh (within the state of Jammu and Kashmir, India), which is the most high energy geothermal field in India (Shanker 1988; Shanker et al., 2012). The Puga geothermal area, which is a part of the greater Ladakh-Tibet borax-spring zone (Harinarayana et al., 2006), lies just south of the tectonically-active collision junction between the Indian and Asian continental crusts involved in Himalayan orogeny (Gansser, 1964). Geological evidences suggest that the subsurface of Puga is underlain by two geothermal reservoirs at two sequential depths (Chowdhury et al., 1974; Chowdhury et al., 1984; Saxena and D’Amore, 1984; Harinarayana et al., 2006) – a shallow heat-reservoir (~160°C) at a depth of ~450 m within the fractured basement rocks (breccia) saturated with hot water; and the deeper main geothermal reservoir (~260°C) commencing at a depth of ~2 km and extending across the upper Himalayan crust up to a depth of ~8 km. Beneath these levels lie the magma chambers or partial melts, which resulted from the high pressures and temperatures generated from the collision and subduction of the Indian continental plate with the Asian plate during Himalayan orogeny (Harinarayana et al., 2006). These molten rocks are the ultimate source of the high heat-flow and geothermal activity in the region (Harinarayana et al., 2006).

### 2.2 *Topography of the* Lotus Pond *hot spring and location of the sample-sites*

Situated on the bank of a mountain brook called *Rulang* (also known as the Puga rivulet), the *Lotus Pond* spring vents profuse hot water (Fig. 2A) with diurnal fluctuations in temperature, flow rate and pH (Fig. 3A; Table 1; Supplementary Table S2). *Lotus Pond*’s vent is seated within an old crater that had apparently formed from the same hydrothermal eruption which had given rise to the spring. On the northern side of the crater there is copious deposit of boratic sinters on the collapsed wall of the crater. These sinters contain old accretions of hydrothermal minerals as well as recent condensates of *Lotus Pond*’s fumarolic vapor (Fig. 2B) (Ghosh et al., 2012). On the southern flank, *Lotus Pond*’s spring water flows into the river *Rulang*, over few-centimeters-high terraces made up of fresh, soft, white sinter deposits. The trajectory of this spring-water transit in the vent-to-river direction represents an 85-14°C thermal gradient, within which four distinct sample-sites (Fig. 2C) were explored on 20 October 2014 for the spring-water’s chemistry and microbial diversity. These sample-sites - designated as (i) HTP (the 85-78°C, <u>h</u>ighest <u>t</u>emperature <u>p</u>oint, situated at the center of the vent); (ii) MTP (the 78-70°C, <u>m</u>oderately-<u>h</u>igh <u>t</u>emperature <u>p</u>oint, situated on the cms-high terraces formed by the sub-aquatic sinter deposits of *Lotus Pond*); (iii) LTP (the lowest temperature point, situated at the interface between the spring-water’s outflow and the *Rulang* river’s water current; here the water temperature fluctuates now and then between 25–45°C); and (iv) RVW (the 14-24°C, <u>r</u>i<u>v</u>er-<u>w</u>ater sampling site, situated within *Rulang*’s own water current, well beyond the LTP site) - were located at *Lotus Pond*’s vent-center, and 1, 1.5 and 1.8 m away from the vent-center, respectively (Table 1). For each sample-site, *in situ* metataxonomic diversity of microorganisms was revealed at 6:30, 8:30, 14:30 and 20:30 h of the sampling-day (see Fig. 4A-D, and refer to Supplementary Table S3 for the summary statistics of the amplified 16S rRNA gene sequence-based OTU analyses for all 16 water-samples explored in this way). Pure culture isolation was carried out from an 80°C, HTP water-sample collected at 15:00 h. Water-chemistry of HTP, MTP, LTP and RVW was explored at 10:30 and 17:30 h, in addition to the four sampling-hours mentioned above, by collecting the water-samples thrice on each occasion (all the values for water chemistry parameters given in Table 1 and Supplementary Table S2 are, therefore, averages of three individual tests).

**Fig. 2.**
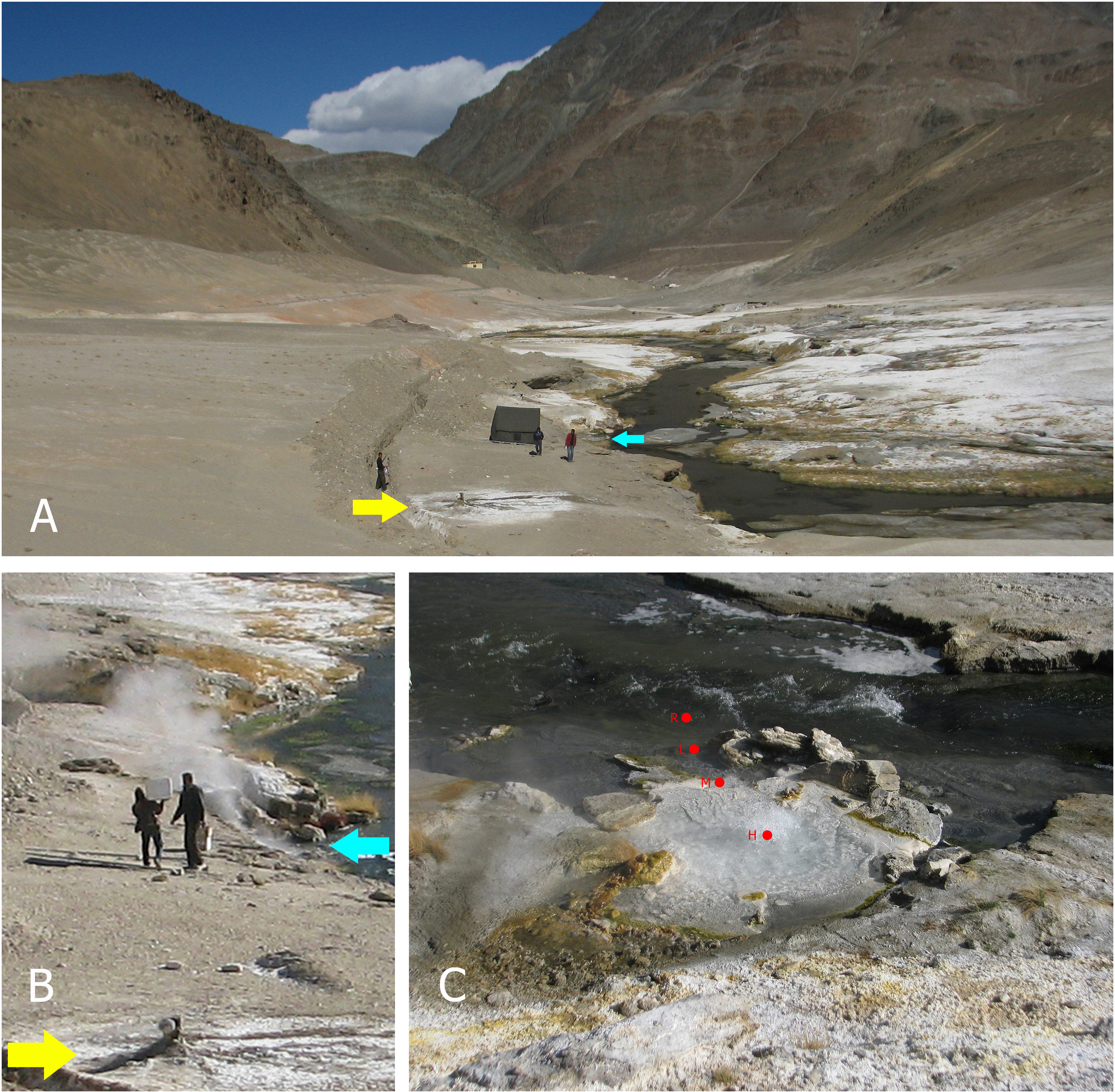
*Lotus Pond* hot spring, *Rulang* river, and the microbialite hot spring *Shivlinga* (microbiota of the latter spring has been studied by Roy et al., 2016; Roy et al., Unpublished), as seen on the sampling-day of this study (20 October 2014): (**A**) environmental context of the entire Puga valley; (**B**) *Lotus Pond* at 8:30 h, showing copious fumarolic activity; (**C**) *Lotus Pond* at 14:30 h, showing almost no fumarolic activity; positions of the four sample-sites HTP (marked as H), MTP (marked as M), LTP (marked as L) and RVW (marked as R) are indicated by red circles. For (**A**) and (**B**), the cyan and yellow arrows demarcate the locations of *Lotus Pond* and *Shivlinga* respectively.

**Fig. 3.**
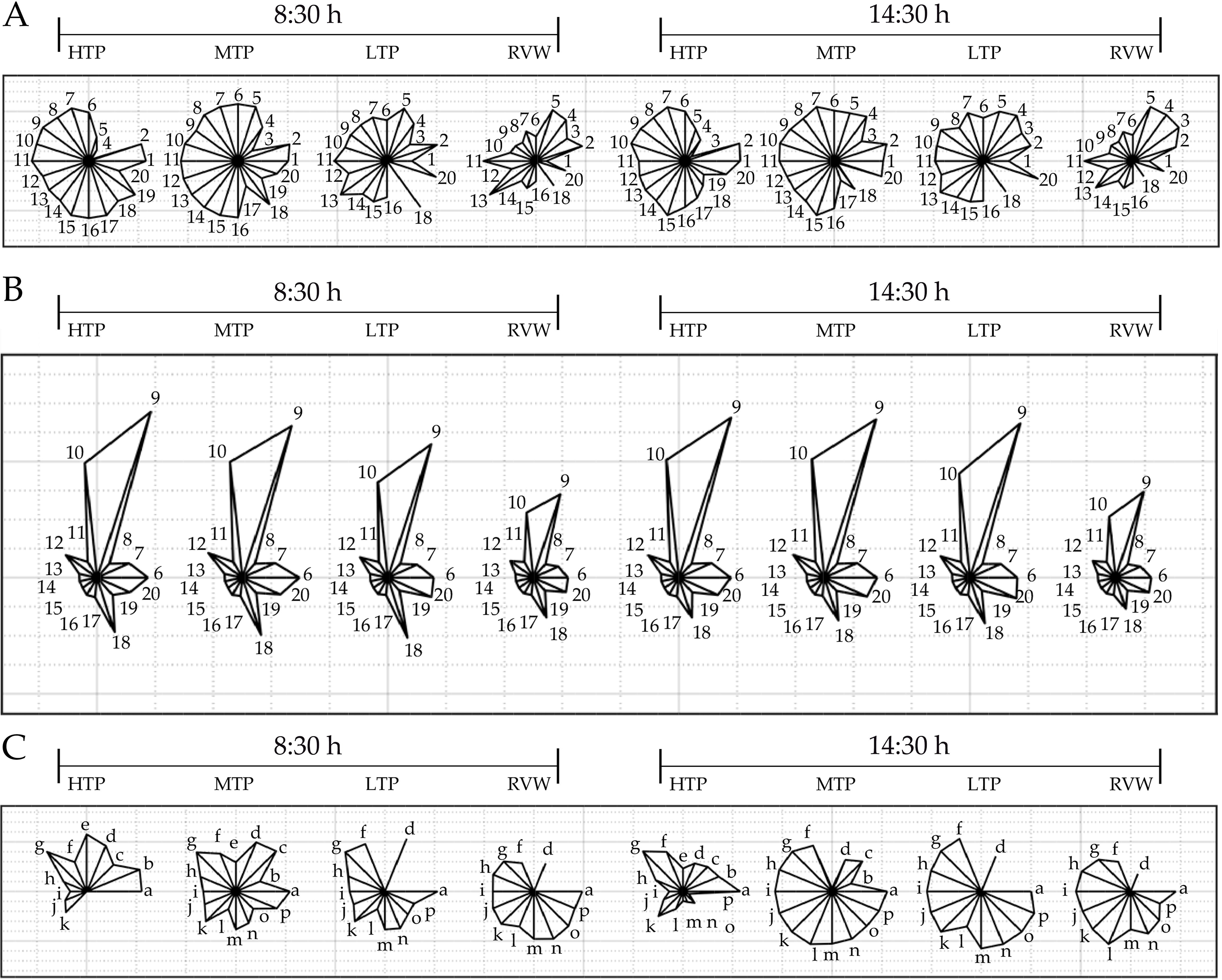
Glyph star plots mapping the data for the major parameters of bacterial diversity and chemistry determined for the water-samples HTP, MTP, LTP and RVW at 8:30 and 14:30 h of 20 October 2014. Glyph plots mapping the normalized values for temperature, pH, number of OTUs detected, number of genera detected, number of phyla/classes detected, concentrations of boron, silicon, lithium, sodium, chloride, calcium, potassium, magnesium, rubidium, strontium, cesium, sulfide, thiosulfate, sulfite, and sulfate (indicated by parameter numbers 1 through 20 respectively), across the sample-sites and sampling-hours; each value under a given parameter was normalized by taking its ratio with the highest value available under that parameter (refer to Supplementary Table S2 for the actual values of all the 20 parameters across the eight samples). (**B**) Glyph plots mapping the matrix-standardized values of the same concentrations for boron, silicon, lithium, sodium, chloride, calcium, potassium, magnesium, rubidium, strontium, cesium, sulfide, thiosulfate, sulfite, and sulfate, that are given in Supplementary Table S2 [parameter numbers are same as those used in (**A**)]. (**C**) Glyph plots mapping the normalized values for (a) pH; (b) concentration of sulfide; (c) δ^34^S of sulfide; concentrations of (d) thiosulfate, (e) sulfite, (f) sulfate, (g) δ^34^S of sulfate, (h) number of S^2−^, S^0^, S_2_ O_3_^2−^ and/or SO_3_^2−^ oxidizing genera present; (i) number of S^2−^ oxidizing genera present; (j) number of S^0^ oxidizing genera present; (k) number of S_2_O_3_^2−^ oxidizing genera present; (l) number of SO_3_^2−^ oxidizing genera present; (m) number of S_2_O_4_^2−^, SO_3_^2−^ and/or SO ^2−^ reducing genera present; (n) number of S_2_O_3_^2−^ reducing genera present; (o) number of SO_3_^2−^ reducing genera present; (p) number of SO_4_^2−^ reducing genera present, across the sample-sites and sampling-hours (refer to Supplementary Table S7 for the actual values of all the 16 parameters across the eight samples).

**Fig. 4.**
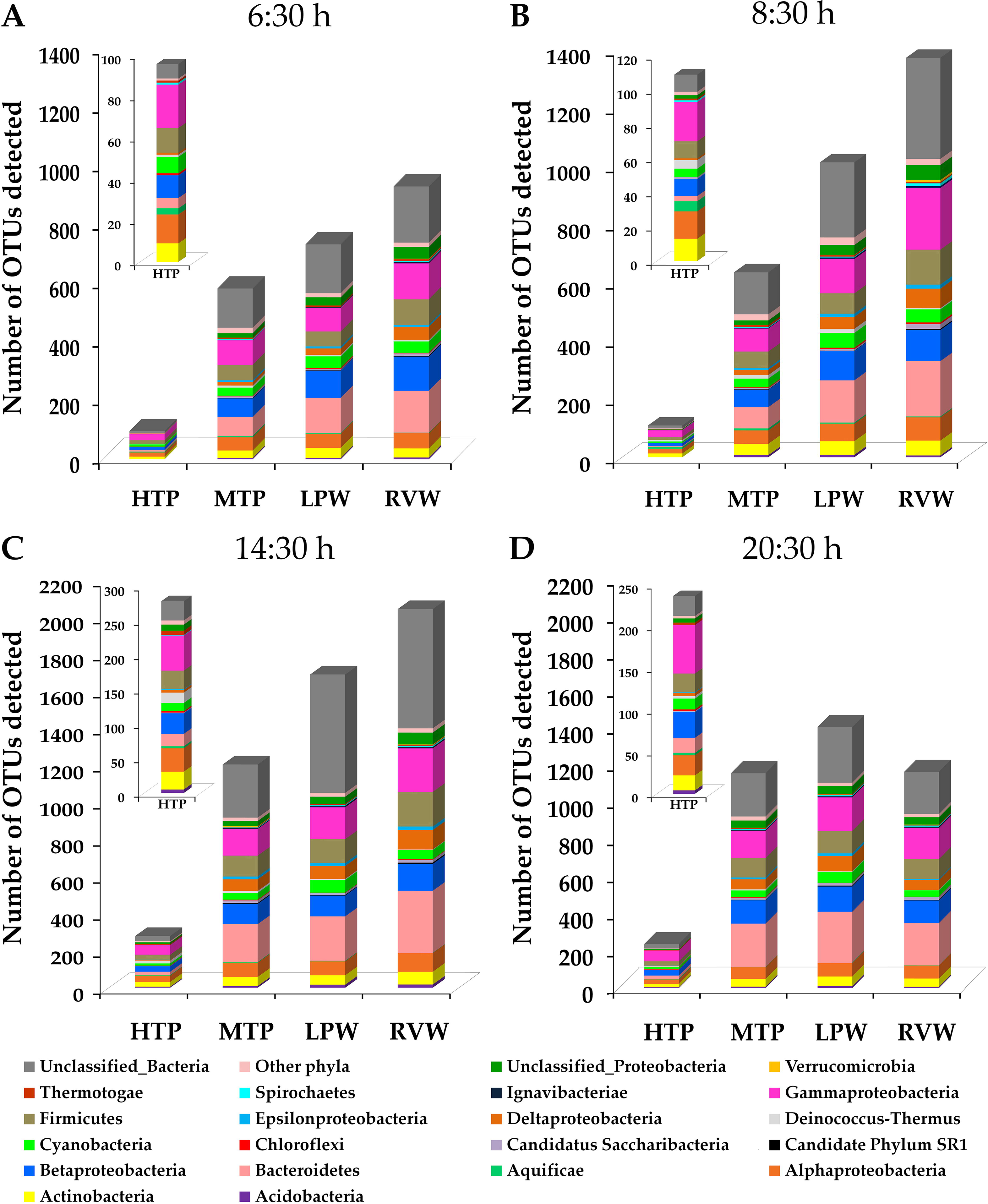
Microbial community architecture revealed along *Lotus Pond*’s hydrothermal gradient: phylum-/class-level distribution of the OTUs identified in HTP, MTP, LTP and RVW at (**A**) 6:30, (**B**) 8:30, (**C**) 14:30 and (**D**) 20:30 h; color-code for the identification of taxa is common for all the four panels.

**Table 1.**
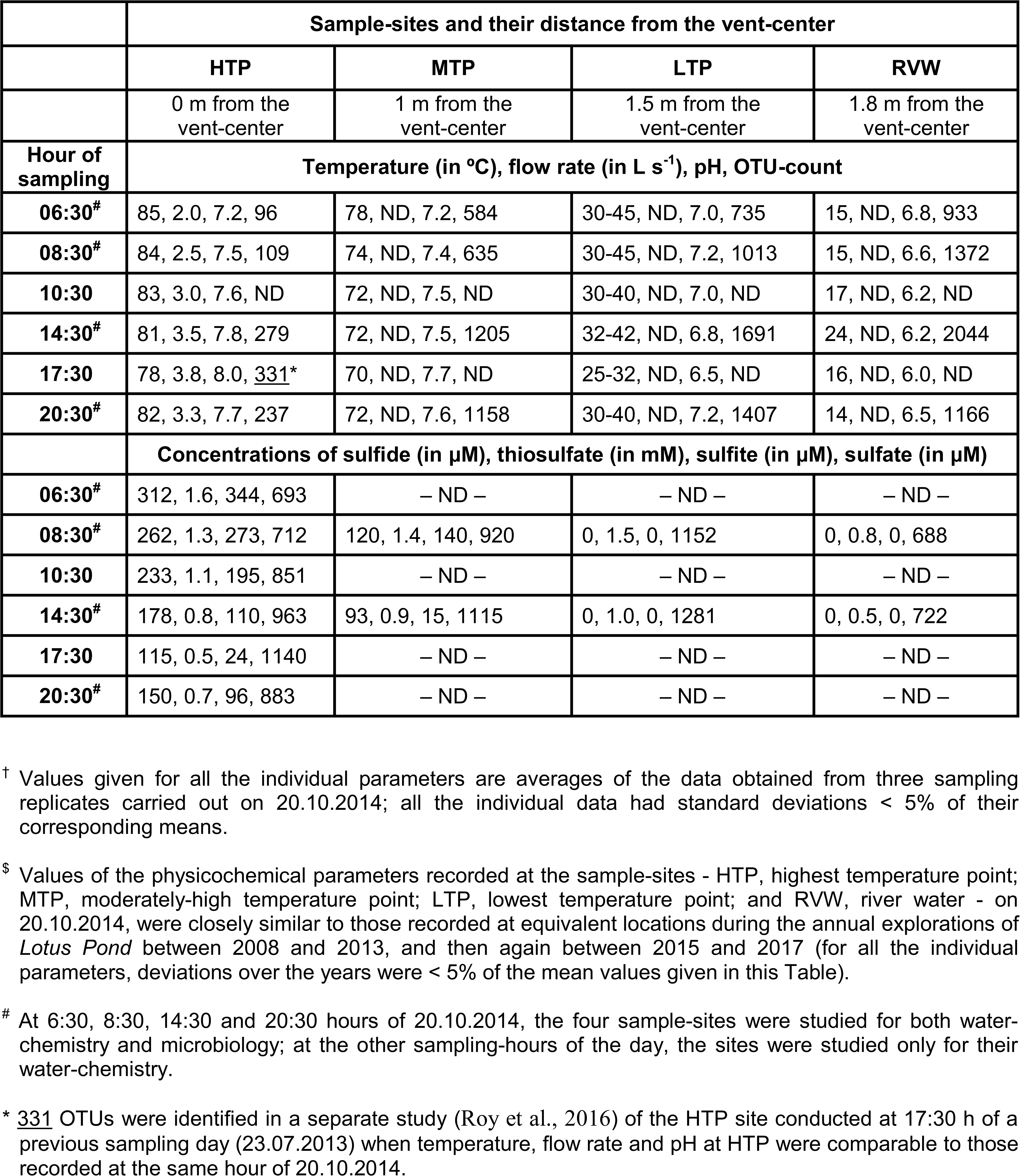
Physicochemical properties^†^ of the sample-sites explored along *Lotus Pond*’s spring-water transit representing a hydrothermal gradient in the vent-to-river trajectory (sampling date: 20.10.2014^$^).

### 2.3. Sample collection for water chemistry

At 6:30, 8:30, 10:30, 14:30, 17:30 and 20:30 h of 20 October 2014, water-samples were collected for chemical analyses from the individual sample-sites using separate 25 mL sterile glass-pipettes on each occasion. Every 100 mL batch of water-sample meant for the quantification of metallic elements was acidified *in situ* (to pH ≤ 2) via addition of 69% (w/v) HNO_3_ (400 μL). 40 mL of each water-sample meant for the determination of sulfate concentration by gravimetric method was acidified *in situ* to pH 2.0 by adding few drops of 1.2 N HCl, and then precipitated for BaSO_4_ by adding 1 mL preheated 1 M barium chloride (BaCl_2_) solution and subsequent vigorous mixing. Dissolved sulfides (ΣHS^−^, which includes H_2_S, HS^−^ and/or S_x_^2–^) were precipitated *in situ* from 25 mL batches of water-samples in the form of cadmium sulfide (CdS), by adding 1 mL 2 M cadmium nitrate [Cd(NO_3_)_2_], inside butyl septum bottles. The bottles were crimp-sealed immediately leaving no head space, and stored at 4°C until further analysis in the laboratory. CdS precipitates meant for sulfur isotope analysis were converted to silver sulfide (Ag_2_S) by the adding silver nitrate (AgNO_3_) solution. Before the above treatments, all water-samples were filtered through 0.22 μm cellulose acetate filters (Sartorius Stedim Biotech, Goettingen, Germany). For all the sample-sites, *in situ* temperatures were measured with a mercury-column glass thermometer; pH values were measured using Neutralit indicator strips (Merck, Darmstadt, Germany). Flow/discharge rate of hot water at the vent (HTP) was determined using Soluble-tracer Dilution technique described previously (Rantz, 1982).

### 2.4. Analytical techniques

Concentrations of boron, calcium, lithium, magnesium, and potassium were determined by inductively coupled plasma mass spectrometry (ICPMS) on a Thermo iCAP QICPMS (Thermo Fisher Scientific, Waltham, Massachusetts, USA), following manufacturer’s protocol; standard calibration curves were prepared using ICPMS standards supplied by Sigma Aldrich (St. Louis, Missouri, USA) and VHG Labs Inc. (Manchester, New Hampshire, USA). Based on replicate analyses of the standards, deviations from actual concentrations were < 2%, < 1%, < 2.1%, < 1% and < 2.5% for boron, calcium, lithium, magnesium and potassium, respectively. Concentration of sodium was determined using an Agilent 240 AA atomic absorption spectrometer (Agilent Technologies, Santa Clara, California, USA), following standard protocol provided by the instrument’s manufacturer; standard curves were prepared from Sigma Aldrich AAS standards. Based on multiple analyses of the standard, deviations from actual sodium concentrations were < 3%. Silicon concentration was determined using a UV-visible spectrophotometer (CARY 100, Varian Deutschland GmbH, Darmstadt, Germany), as described previously (Grasshoff et al., 1999). Double distilled nitric acid, prepared using a teflon acid purification system (Savillex, EdenPrarie, Minnesota, USA), and ultra pure water, prepared using a Chorus laboratory water purification system (ELGA LabWater, High Wycombe, UK), were used for sample preparation during the analysis. Chloride was quantified by precipitation titration with silver nitrate (0.1 N) (Mazumdar et al., 2009) using a Titrino 799GPT auto titrator (Metrohm AG, Herisau, Switzerland).

Thiosulfate and sulfate concentrations were determined by iodometric titration, and gravimetric precipitation with barium chloride, respectively (Kelly and Wood, 1994; Alam et al., 2013). Sulfite was analyzed spectrophotometrically with pararosaniline hydrochloride (Sigma-Aldrich) as the indicator (West and Gaeke, 1956). Dissolved sulfides that were precipitated as CdS from the water-samples *in situ* were subjected to colorimetric measurement based on the principle that N, N-dimethyl-p-phenylenediamine dihydrochloride and H_2_S react stoichiometrically in the presence of FeCl_3_ and HCl to form a blue-colored complex (Cline, 1969). To corroborate the results obtained by the above methods, sulfate, sulfite and thiosulfate were additionally quantified in the CdS-precipitated ΣHS^−^-free water-samples using a Metrohm ion chromatograph (Basic IC plus 883) equipped with a suppressed conductivity detector (Metrohm, IC detector 1.850.9010) and a MetrosepASupp 5 (150/4.0) anion exchange column (Metrohm AG, Herisau, Switzerland).

To determine their stable sulfur isotope ratios, dried and homogenized BaSO_4_ or Ag_2_S precipitates were mixed with V_2_O_5_, flash combusted at 1,150°C in an EA1112 elemental analyzer (Thermo Fisher Scientific), and then analyzed using a Thermo Delta V plus continuous flow isotope ratio mass spectrometer (Thermo Fisher Scientific) (Fernandes et al., 2018). All results were reported in standard delta notation (δ^34^S) as per mil (‰) deviations from the VCDT (Vienna Canyon Diablo Troilite), with reproducibility of ± 0.3 ‰. IAEA standards S-1, S-2, SO-5 and SO-6 were used for instrument calibration.

### 2.5. Sample collection for analyzing aquatic communities

At 6:30, 8:30, 14:30 and 20:30 h of 20 October 2014, water was collected from each of the four sample-sites in multiple 1000 mL batches for total environmental DNA isolation and analysis; separate 25 mL sterile glass-pipettes were used for each sampling occasion. Additionally, at 15:00 h, multiple 1000 mL batches of water were sampled from the HTP site (*in situ* temperature 80°C) for pure culture isolation. Each 1000 mL batch of water-sample was passed through a sterile 0.22 μm cellulose acetate filter 4.7 cm in radius (Sartorius Stedim Biotech, Goettingen, Germany), and the filter was inserted into a cryovial containing 5 mL of 50 mM Tris:EDTA (TE, pH 7.8) or 15% glycerol plus 0.9% NaCl solution, according as the sample was meant for environmental DNA-based analysis or pure culture isolation. After filter-insertion each cryovial was sealed with Parafilm (Bemis Company Inc., Neenah, WI, USA), packed in polyethylene bags, and immediately put into dry ice. In the laboratory, samples destined for culture-independent and culture-dependent analyses were stored at −20°C and 4°C respectively.

### 2.6. Preparation of total environmental DNA from water-samples

Total environmental DNA was extracted from the microbial cells arrested in the 0.22 μm cellulose acetate filters, thereby ensuring that no cell-free DNA from the spring/river water-samples infiltrated the preparation (Roy et al., 2016). Each filter was cut into small pieces with sterile scissors, within the TE-containing 8 mL cryovial in which it was brought from Puga. The cryovial was vortexed for 30 min, following which the filter shreds were discarded, and the residual TE was distributed equally into five 1.5 mL microfuge tubes. Each 1.5 mL microfuge tube was centrifuged (10,800 *g* for 30 min, at 4°C) and then its upper 900 μL supernatant was discarded; the residual 100 μL TE was vortexed for 15 min. Subsequently, the contents of all the five 1.5 mL microfuge tube were pooled up in a fresh 1.5 mL microfuge tube. The pooled TE (500 μL) was again centrifuged (10,800 *g* for 30 min, at 4°C), following which the upper 400 μL supernatant was discarded. The remaining 100 μL contained all of the microbial cells that were present in the original 1000 mL water-sample. Total environmental DNA was isolated from this 100 μL cell suspension by the QIAamp DNA Mini Kit (Qiagen, Hilden, Germany), following the manufacturer’s protocol.

### 2.7. Assessment of metataxonomic diversity within the Lotus Pond-Rulang ecosystem

V3 regions of all bacterial/archaeal 16S rRNA genes present in an environmental DNA preparation were PCR amplified using Bacteria-/Archaea-specific universal oligonucleotides, following the fusion primer protocol described previously (Ghosh et al., 2015; Roy et al., 2016; Fernandes et al., 2018). Amplification was carried out using a 16S forward primer prefixed with an Ion Torrent adapter and a unique sample-specific barcode or multiplex identifier in the following 5′ to 3′ organization: (a) a 26-mer A-linker followed by a 4-mer A-linker key common for all sample-specific primers (see bases in bold fonts in the sequences given below), (b) a 10-mer barcode unique to each sample-specific primer, which is followed by a common 3-mer barcode adaptor (all denoted as stars in the sequences below), and then (c) the relevant domain-specific universal forward primer in its 5′ to 3′ direction (see underlined bases in the sequences given below). A reverse primer, in contrast, had (a) a common trP1 adapter (see bases represented in italics), followed by (ii) the relevant domain-specific universal reverse primer in its 5′ to 3′ direction (see underlined bases in the sequences given below). In this way, V3 portions of all bacterial 16S rRNA genes present in a sample DNA were amplified using the forward primer 5′ – **CCA TCT CAT CCC TGC GTG TCT CCG ACT CAG *** *** *** \*\*\*\***CC TAC GGG AGG CAG CAG – 3′ and the reverse primer 5’-*CCT CTC TAT GGG CAG TCG GTG AT*A TTA CCG CGG CTG CTG G - 3′ (where the underlined segments represent the universal primers 341f and 515r respectively). For the amplification of archaeal 16S rRNA genes, we used the forward primer 5’ - **CCA TCT CAT CCC TGC GTG TCT CCG ACT CAG *** *** *** *** \***AA TTG GAN TCA ACG CCG G –3’ (where the underlined bases represent the universal primer 344f) and the reverse primer 5’ - *CCT CTC TAT GGG CAG TCG GTG A*TC GRC GGC CAT GCA CCW C – 3’ (where the underlined bases represent the universal primer 522r).

PCR products from all the samples were pooled up at equal concentrations for sequencing on Ion PGM (Thermo Fisher Scientific). Prior to sequencing, size distribution and DNA concentration within the amplicon pool was checked using a Bioanalyzer 2100 (Agilent Technologies) and adjusted to 26 pM. Amplicons were then attached to the surface of Ion Sphere Particles (ISPs) using an Ion Onetouch 200 Template kit (Thermo Fisher Scientific). Manually enriched, templated-ISPs were then sequenced by PGM on an Ion 316 Chip for 500 flows. Before retrieval from the sequencing machine, all reads were filtered by the inbuilt PGM software to remove low-quality and polyclonal-sequences; sequences matching the PGM 3′ adaptor were also trimmed. Quality-filtered readsets were exported as fastq files for downstream analyses. The individual sequence files were deposited to Sequence Read Archive (SRA) of the National Center for Biotechnology Information (NCBI, Bethesda, MD, USA), under the BioProject accession number PRJNA296849, with the distinct run accession numbers listed in Supplementary Tables S3.

All V3 sequence reads were filtered a second time for high quality value (QV 20) and length threshold of 100 bp; selected reads were then converted to fasta from fastq using Fastx_toolkit 0.0.13.2 (http://hannonlab.cshl.edu/fastx_toolkit/download.html). Reads obtained for the individual water-samples were clustered into OTUs unified at the 97% sequence similarity level using the various modules of UPARSE (Edgar, 2013), and singletons were discarded. A Perl programming-script, available within UPARSE, was used to determine the Abundance-based Coverage Estimator, and Shannon and Simpson Indices. Rarefaction analysis was carried out using the Vegan Package in R (Team, 2014). The consensus sequence of every OTU was taxonomically classified by the RDP Classifier tool located at http://rdp.cme.msu.edu/classifier/classifier.jsp.

### 2.8. Bubble and Glyph plots

For visual comprehension of the complex information contained in our multidimensional data Glyph and Bubble charts/plots were constructed (Few, 2009; Kelleher and Wagener, 2011; Ware, 2012; Kosara and Mackinlay, 2013). Bubble chart is one category of scatter plot, where one can depict three-dimensional data in a two-dimensional space by using the X and Y axes for two independent variable and size of bubbles for the dependent variable. Multi series bubble chart can also be plotted with different bubble colors/shades to include more than one dependent variable (Few, 2009). The single series bubble plot shown in Fig. 1 was constructed using the *Scatter* tool in ‘MATLAB 2017b’ (Martinez et al., 2017); the syntax used was: scatter(datafile.Temperature(1:32), datafile.pH(1:32), datafile.OTUs(1:32),‘k’); hold on scatter(datafile.Temperature(33:end), datafile.pH(33:end), datafile.OTUs(33:end),‘r’);

Glyph plots were constructed for visual comparison of the multidimensional geomicrobiological data. A glyph conveys every aspect of a multidimensional data mapped in terms of features such as position, length/size, shape, color, etc. (Ribarsky et al., 1994). To construct the various Glyph stars of Fig. 3, the *Glyphplot* package in ‘MATLAB 2017b’ (Martinez et al., 2017) was used with the syntax: glyphplot(data, ‘glyph’, ‘star’, ‘Color’, ‘black’, ‘standardize’, ‘column’, ‘centers’, [<co-ordinates of the glyph centres>], ‘radius’, <radius of the glyphs>).

### 2.9. Isolation of mesophilic bacterial strains from HTP water-samples

Sterile 0.22 μm cellulose acetate filters through which batches of 1000 mL HTP water-samples had been passed during sampling, and those which were rehydrated in 5 mL 15% glycerol plus 0.9% NaCl solution, were cut into pieces with sterile scissors, within the same 8 mL cryovials in which the filters were brought from the field. The cryovials were vortexed for 30 min, following which the filter shreds were discarded, and the residual (~5 mL) glycerol-NaCl solutions were added to 25 mL R2A or MST broth medium. 1 L R2A (pH 7.0) contained 0.5 g proteose peptone, 0.5 g casamino acids, 0.5 g yeast extract, 0.5 g dextrose, 0.5 g soluble starch, 0.3 g K_2_HPO_4_, 0.05 g MgSO_4_.7H_2_O, 0.3 g sodium pyruvate, while 1 L MST (pH 7.0) contained 1 g, NH_4_Cl; 4 g, K_2_HPO_4_; 1.5 g, KH_2_PO_4_; 0.5 g, MgSO_4_.7H_2_O; 5 g (20 mM), Na_2_S_2_O_3_.5H_2_O; 50 mg, yeast extract (as growth factor supplement); and 5 mL, trace metals solution (Ghosh and Roy, 2006). The mixtures were incubated at 37°C on a rotary shaker (180 rpm) until OD_600_ of the R2A cultures reached 0.8, or the pH of the spent MST medium became < 6 due to conversion of thiosulfate to sulfuric acid. Serial dilutions from the mixed cultures were then plated on R2A- or MST-agar, and incubated at 37°C. Single colonies of pure strains, apparently unique in terms of colony morphology and/or rate and extent of acid production in chemolithoautotrophic MST medium, were isolated as pure cultures and maintained in Luria-Bertani (LB) agar slants or MST-agar plates, at 37°C.

### 2.10. Growth of SMMA_5 at different temperatures in the absence/presence of kosmotropic or compatible solutes

Cellular growth of *Paracoccus* SMMA_5 was recorded at regular intervals over 12 h of incubation in chemolithoautotrophic MST broth at 37°C, 40°C, 45°C, 50°C, 60°C and 70°C. Growth was also recorded over 50 min and 12 h of incubation, at 37°C, 50°C and 70°C, in different variants of the MST medium, such as MST-Li (MST supplemented with 1.0 mM LiOH.H_2_O), MST-B_4_O_7_ (MST supplemented with 4.0 mM Na_2_B_4_O_7_.10H_2_O), MST-SO_4_ (MST supplemented with 1.0 mM Na_2_SO_4_), MST-Na (MST supplemented with 17.0 mM NaCl) and MST-GB (MST supplemented with 10.0 mM C_5_H_11_NO_2_). In all these experiments, inocula from early log phase cultures of SMMA_5 grown in LB broth at 37°C (OD_600_ ~0.3) were transferred (1% v/v) to relevant MST-based experimental broths and incubated at the temperature specified for the experiment. To determine the number of CFUs or ready-to-divide cells present mL^−1^ of an experimental culture at any time-point of incubation (including the 0 h), its various dilution grades were plated in triplicates on to LB-agar and single colonies counted in each of them after 36 h incubation at 37°C. Colony-counts in the different dilution-plates were multiplied by their respective dilution factors, then summed-up across the plates, and finally averaged to get the number of CFUs that were present mL^−1^ of the experimental culture. Cellular growth yield after a given time-period of incubation was expressed as what percentage of the 0 h CFU-count still remained ready-to-divide after that period of incubation.

### 2.11. Determination of glycine-betaine concentration in spent medium

The *Paracoccus* isolate SMMA_5 was incubated in MST broth for 12 h at 50°C. Cells were precipitated out from 100 mL spent MST medium by centrifugation at 6000 *g* for 20 min. The supernatant was lyophilized down to 1 mL final volume, and then filtered by passing through 0.22 μm cellulose acetate filter (Sartorius Stedim Biotech). 25 μL of the filtrate was analyzed by High Performance Liquid Chromatography (HPLC) using standard UV detection as described previously (Zamarreño et al., 1997). HPLC was carried out on a Waters platform encompassing a Waters FlexInject sample-injector, Waters 1525 Binary HPLC pump, a Waters 2998 photodiode array detector, and the software Breeze 2.0 (Waters Corporation, Milford, Massachusetts, USA). Isocratic elution was carried out in a Hypersil SCX column having 5 μm particle size, 250 mm length and 4.6 mm inner diameter (Phenomenex, Torrance, California, USA), using a mixture of disodium phosphate buffer (0.05 M, pH 4.7) and methanol (95:5%, v/v) as the mobile phase (flow rate: 1 mL min^−1^, at room temperature; detection wavelength: 195 nm; and injection volume: 25 μL). Standard glycine-betaine (Sigma Chemical Company, St. Louis, Missouri, USA) was dissolved in distilled water at final concentrations of 0, 2, 4, 8 and 10 mM and analyzed as above to generate the calibration curve. Glycine-betaine in a sample was identified by comparing its retention time with that of the standard; purity of chromatographic peaks was checked using Breeze 2.0. Glycine-betaine in a sample was quantified based on the area of its chromatographic peak using the calibration curve Y = mX + C, where Y is the peak-area; X is the concentration of glycine-betaine; m is the slope of the calibration curve; C is the curve intercept for the sample.

### 2.12. Determining the percentage of viable cells in an experimental culture by flow cytometry

After incubation in MST, MST-Li, MST-B_4_O_7_, MST-SO_4_, MST-Na or MST-GB broths for 50 min at 70°C, SMMA_5 cultures were tested for the proportions of viable and non-viable cells present, using flow cytometry (Battin, 1997; Samaddar et al., 2016). In these experiments, inocula from early log phase cultures of SMMA_5 grown in LB broth at 37°C (OD_600_ ~0.3) were transferred (3% v/v) to relevant MST-based experimental broths and incubated at 70°C. After incubation, cells were precipitated from 100 mL spent media by centrifugation at 6000 *g* for 20 min. The supernatants were discarded and the cell pellets resuspended in 2 mL 0.9% NaCl solution. 4 μl FDA (Sigma Chemical Company) (5 mg mL^−1^ in dimethyl sulfoxide) was added to the resuspended cells solutions following which cells were incubated for 15-20 min at 37°C. Incubated cells were washed and resuspended again in 0.9% NaCl solution (final resuspension was done in 500 μl 0.9% NaCl solution), and analyzed using a FACSVerse flow cytometer (Becton Dickinson, New Jersey, USA). Fluorescence was measured through the excitation wavelengths 475-495 nm and the emission wavelengths 520-530 nm. 10000 randomly-taken cells were analyzed for each sample and dot plots were generated by plotting the fluorescence of each cell against its forward scattering of 488 nm wave detected by a photodiode array detector. The data were analyzed using the software BD FACSuite (Becton Dickinson) by specific quadrant gating for each experiment that in turn were predetermined based on unstained samples.

## 3. Results and discussion

### 3.1. High microbial diversity along Lotus Pond’s spring-water transit

Total environmental DNA yield from all the individual water-samples was < 30 ng 1000 mL^−1^, which was quantitatively insufficient for direct shotgun sequencing; so, their microbial diversities were revealed by PCR-amplification and sequencing of the V3 regions of all 16S rRNA genes present in the environmental DNA preparations. For all the samples, Bacteria-specific V3 primers generated PCR products of desired size (~200 bp) that were subsequently sequenced using an Ion Torrent Personal Genome Machine (Ion PGM). Archaea-specific V3 primers, however, did not yield any PCR product, suggesting that very low numbers of archaeal cells are present in these water-samples. Reads of the individual Bacteria-specific V3 sequence datasets were clustered into operational taxonomic units (OTUs) that represented putative species-level entities unified at the 97% sequence similarity level. Rarefaction analysis of all the 16 datasets confirmed that their extents of read-sampling (sequence data throughput) were sufficient to reveal most of the OTU-level diversity present (Supplementary Fig. S1).

The undetectable status of archaea in *Lotus Pond*’s diversity analyses was intriguing because metagenomic data from almost all the well-studied hot springs of the world reveal the presence of archaea (Kvist et al., 2007; Huang et al., 2011, 2013; Wemheuer et al., 2013; Hedlund et al., 2015). Whilst the Archaea-specific primers used in the present study may have failed to amplify the particular taxa present *in situ*, it is, at the same time, worth exploring whether the overwhelming presence of bacteria in *Lotus Pond* have had any antagonistic effect on the native archaeal populations. This is because cases of tandem high and low abundances of bacteria and archaea respectively, has been reported from a number of hot springs across the world (López-López et al., 2015; Hussain et al., 2017). Some of these instances have been attributed to the alkaline pH of the spring-waters concerned, whereas high relative abundance of archaea, elsewhere, has apparently been linked with low pH levels (Menzel et al., 2015). It, therefore, is not improbable that the neutral to slightly alkaline pH of the *Lotus Pond* spring-water, in conjunction with the bacterial influx caused by mixing of cold meteoritic waters from low temperature aquatic systems (see below), has some role in the bacterial predominance of this ecosystem.

High bacterial diversities were revealed along *Lotus pond*’s hydrothermal gradient across the different sampling-hours of 20 October 2014 (Fig. 4A-D). This was consistent with the results of another exploration of the *Lotus Pond* vent-center (the same sample-site as the HTP of this study) at 17:30 h of a previous sampling day (23 July 2013) (Roy et al., 2016; see also Fig. 1 and Table 1). Collectively, all these results pointed out a higher habitability of this ecosystem in comparison to other hot springs having equivalent temperature and pH (Fig. 1; also see Supplementary Table S1 and references therein). For instance, among the global hydrothermal sites having temperatures > 70°C, none other than *Lotus Pond* MTP and the *Elegedi* spring in Eritrea harbors > 700 OTUs. Whilst the *Elegedi* spring has a temperature of 100°C, for *Lotus Pond*, situated at 4,436 m above the mean sea level, the heat-contents of its 70-85°C water-samples are also either equal to, or quite close to, that of boiling water (notably, both *Lotus Pond* MTP and *Elegedi* both have neutral to near-neutral pH). Furthermore, it is remarkable that all > 70°C hydrothermal sites, whether situated at high altitudes or in the plain lands, harbor < 100 OTUs; the only few exceptions to this are *Lotus Pond* MTP (pH 7.2-7.7), *Elegedi* (pH 7), and *Hot Springs 2* and *3* in *Lake Magadi*, Kenya (pH 9.2-9.4). At the same time it is noteworthy that all < 70°C hydrothermal sites - except the *Eiland* hot spring of South Africa - harbor > 100 OTUs. Whereas spring-waters having near-neutral to moderately-alkaline pH, irrespective of their temperatures, are clearly associated with high microbial diversity, no spring having pH < 7, except the Icelandic hot springs P1 through P3, have > 100 OTUs reported. Moreover, *Lotus Pond* and *Elegedi* are apparently the only boiling/near-boiling and pH neutral/near-neutral hydrothermal sites that have > 100 OTUs (in fact *Elegedi* and some *Lotus Pond* sites have > 1000 OTUs).

Albeit the number of OTUs identified in a habitat varies depending on the primer sets, sequencing platforms and analysis pipelines used, all the differences reported in the OTU-counts of geographically and geologically diverse hot springs cannot be solely due to technological inconsistencies. This is because the relevant studies had all followed standard, reproducible procedures of 16S rRNA gene fragment amplification, sequencing, and OTU-clustering, including rarefaction analyses. So, even after factoring in some degree of discrepancy due to methodological variations, the body of literature available regarding the biodiversity of hot springs across the world collectively indicates that all high temperature habitats do not harbor the same number of OTUs. In the context of the current investigation, the main objective was to evaluate whether the OTU-counts obtained for the various *Lotus Pond* samples were normal, high or low, as compared to the OTU-counts reported from other hot springs across the world. Once we came to know (via this comparison) that very few hot springs of equivalent temperatures harbor diversities as high as *Lotus Pond*, the imperative was to check whether the OTU-counts obtained for the *Lotus Pond* samples involved overestimation of diversity. Towards the latter objective we applied different OTU-clustering methods on all the *Lotus Pond* sequence datasets, and found UPARSE-based analyses (data from which were eventually used in this paper) to give the lowest number of OTUs, as compared to the other well-known clustering algorithms such as ESPRIT-Tree, MOTHUR, dada2 and deblur within QIIME2. This conclusion for *Lotus Pond*, irrespective of whether the instances of high OTU-count in comparator hot springs were attributable to inconsistent methodologies, was considered as a premise strong enough to start asking and addressing new questions on (i) the plausible mechanisms / physicochemical bases of microbial transportation across hydrothermal ecosystems, (ii) the fate of “accidentally-introduced” mesophilic microorganisms in the face thermal stress, and (iii) the potential secondary/environment-guided adaptations that microbes may undergo when thrust into unfamiliar environmental challenge (such as high heat) by physical forces of nature.

On the other hand, with regards to the diversity comparison between *Lotus Pond* and other global hot springs, it is noteworthy that low diversity-level is the expected phenomenon for hot springs in general, and the same was found to hold true in the literature available for most of the comparator hot springs. In this way, all doubts that can potentially be there regarding the data used in this comparison, converges towards the hot springs having high diversity. Whereas within the scope of the present study there was no room for deciphering what percentages of the high diversities reported for the few comparator hot springs were actually attributable of contaminations (whether intrinsic to the physiography of the ecosystem itself, or attributable to field techniques, laboratory reagents or researchers’ bodies), the no sample” controls included in all our experiments did not yield any measurable DNA either at the metagenome isolation stage or the PCR amplification stage. This confirmed that the diversities revealed indeed came from *Lotus Pond* samples.

### 3.2. Spatiotemporal fluxes in microbial diversity along Lotus Pond’s spring-water transit

Bacterial OTUs identified from the four samples-sites of the *Lotus Pond*-*Rulang* system at two time points each in the forenoon (6:30 and 8:30 h) and afternoon (14:30 and 20:30 h) of the sampling-day were first classified and analyzed at the phylum-level (Fig. 4A-D; Supplementary Table S4). Results showed the individual communities, especially the one at HTP, to remain largely consistent over the 2 h time-span surveyed in the forenoon (i.e., between 6:30 and 8:30 h; compare Fig. 4A and B), and then again over the 6 h time-span surveyed in the afternoon (i.e., between 14:30 and 20:30 h; compare Fig. 4C and D), even though OTU-counts increased and decreased to some extents in-between the 2 h and 6 h time-frames respectively. Rise in OTU-count at individual sample-sites, however, was much steeper across the forenoon-afternoon divide (i.e., between 8:30 and 14:30 h; compare Fig. 4B and C), in view of which OTUs identified in these 8:30 and 14:30 h datasets were further analyzed at the genus-level. Consequently, detailed geochemical data from all the four sample-sites were also compared for these two time-points only.

For all the individual sample-sites, total number of genera over which the OTUs were distributed, were also found to increase significantly at 14:30 h, as compared to 8:30 h (more of the genus-level analyses can be seen in the subsequent sections). For all the four sample-sites explored, temporal changes in OTU-count notwithstanding, the proportions of OTU-affiliation across different phyla remained largely unchanged across the sampling-hours. For instance, 60-75%, of the OTUs present at any sampling-hour, in any sampling site, were affiliated to *Actinobacteria*, *Alphaproteobacteria*, *Betaproteobacteria*, *Gammaproteobacteria*, *Firmicutes* and Unclassified *Bacteria* (Supplementary Table S4). This not only reflected the contiguous nature of the aquatic communities explored along *Lotus Pond*’s spring-water transit, but also indicated a qualitative consistency of the microbiome in both time and space.

On the spatial scale, total number of OTUs present at individual sample-sites increased steadily along *Lotus Pond*’s hydrothermal gradient in the vent-to-river trajectory, at all the four sampling-hours, except for a small decrease from LTP to RVW at 20:30 h (Fig. 4D). Site to site increases in diversity, remarkably, were maximal between HTP and MTP, where 6.1, 5.8, 4.3 and 4.9 fold increases in OTU-count were recorded at 6:30, 8:30, 14:30 and 20:30 h respectively. OTU-count for almost every individual phylum detected in the system also increased sharply from HTP to MTP, at all the four sampling-hours, except for *Aquificae* at 14:30 and 20:30 h, and *Deinococcus*-*Thermus* and *Thermotogae* at 14:30 h (Supplementary Table S4). These steep rises in diversity, at all the four sampling-hours, coincided with mere 7-10°C drops in temperature between HTP and MTP, which could be reflective of a strong habitability-barrier, somewhere between 85°C and 72°C, along the hydrothermal gradient. Below 72°C, however, thermal constraint on diversity was clearly less acute, as OTU-count for individual sample-sites, or the number of phyla over which OTUs were distributed, increased gradually or even decreased in some cases, at all the sampling-hours of the day. In this context it is noteworthy that a recent study conducted across 925 geothermal springs of New Zealand has also pointed out that temperature has a significant effect on measured *in situ* diversity at values > 70°C, whereas diversity is primarily influenced by pH at sites having temperatures < 70°C (Power et al., 2018). This said, possibilities remain that the sharp rises in diversity at MTP (as compared to HTP) is augmented by the slowing down of the spring-water in transit over the centimeters-tall terraces of *Lotus Pond*’s apron at MTP, which in turn causes the microorganisms, coming into the system along with the underground geothermal water, accumulate in the MTP territory (see next Results section for more on microbial influx from the subsurface). Furthermore, interaction of the flowing spring-water with the phototrophic mat communities growing along the banks of the outflow channel (these are visible in Fig. 2C) may also contribute to the increased diversity at MTP.

### 3.3. Spatiotemporal fluxes in the physicochemical conditions can account for the biodiversity fluxes

From the early morning (6:30 h) till the evening (17:30 h) of the sampling day, temperature of the vent-water (HTP) decreased gradually from 85°C to 78°C; correspondingly, vent-water discharge increased from 2 to 3.9 L s^−1^. With the onset of darkness, however, there was a quick rise in the temperature to 82°C (as recorded at 20:30 h), accompanied by a drop in the discharge rate to 3.4 L s^−1^ (Table 1). It is known from previous geological studies that the hydrothermal discharges of Puga springs come from a reservoir of formation waters trapped in geological sediments; this reservoir which is distinct from the aquifer, is said to be recharged by melting snow in the higher mountain reaches above 5,600 m (Navada and Rao, 1991). Furthermore, stable isotope ratios for hydrogen and oxygen in these hot water discharges are indicative of their brief underground-retention time and moderate interaction with the underlying rocks (Navada and Rao, 1991). When these facts are considered in conjunction with the negative correlation observed between the diurnal variations in the temperature and discharge-rate of *Lotus Pond* vent-water (Pearson correlation coefficient *R* = −0.9, with corresponding probability value *P* = 0.007) it is evident that with progressive rise in air temperature over the day, snow melts in the higher mountain reaches and increasing volumes of cold oxygenated water is pumped into the hypothermal reservoir, leading to higher discharge volumes and lower temperatures of the vent-water.

The above mentioned geophysical dynamics could be one of the potent causes of diurnal biodiversity flux in *Lotus Pond*’s vent-water as well as in the downstream of it. For instance, in sync with the progressive increase in vent-water discharge from the early morning (6:30 h) till the evening (17:30 h), OTU-count increased at all the four sample-sites of the *Lotus Pond*-*Rulang* system. These hikes, remarkably, were maximal across the forenoon-afternoon divide, i.e. between 8:30 and 14:30 h - maximum increase (2.6 times), at this temporal juncture, was recorded at HTP, whereas minimum increase (1.5 times) was recorded at RVW (compare Fig. 4B and C). We, therefore, hypothesized that as more melting-snow waters enter the tectonic faults which crisscross the region and lead to the underlying hydrothermal reservoir (Harinarayana et al., 2006), denudation/wearing away of the sediments and breccia potentially introduces soil/sub-surface microorganisms into it. These microbes, together with those present in the snow-melts, are potentially transported through the hot-water conduit and eventually ejected through *Lotus Pond* and several other vents spread across the valley. A similar phenomenon has recently been reported from weakly-acidic (pH 4.0-6.0) hot springs across Yellowstone National Park, USA, where the mixing of meteoric and geothermal waters gives rise to wide-ranging disequilibria in redox reactions that, in turn, support idiosyncratically higher chemosynthetic microbial diversity, as compared to those encountered in the springs having minimal or no mixing of the water-types (Colman et al., 2019). That the recharge of the Puga hydrothermal reservoir by melting snow over the day-time indeed causes influx of microorganisms in the vent system was further corroborated by the simultaneous reversing of the incremental trends of biodiversity flux as well as vent-water discharge, with the onset of darkness. For instance, at 20:30 h, in sync with the decrease in vent-water discharge there were subtle decreases in the OTU-counts at HTP, MTP and LTP, and a conspicuous (~40%) decrease in the OTU-count at RVW (Table 1). In fact, the decrease in RVW OTU-count at 20:30 h was so sharp that for the first occasion in the entire sampling-day it became even lower than the OTU-count of LTP (notably, at 20:30 h, OTU-count of RVW was almost same as that of MTP). This last observation, together with the elements of consistency detected in the microbial diversity of the individual sample-sites across the day, reflected that the microbial communities along *Lotus Pond*’s spring-water transit were not necessarily being washed away, all the while, into the river, and the higher OTU-count of RVW (as compared to HTP, MTP and LTP) observed throughout the day-time was not due to dumping of microorganisms by *Lotus Pond*’s outflow. Instead, the latter now seemed attributable to the river’s own denudation/transportation dynamics rooted in its visibly higher water flow between 6:30 and 17:30 h, as compared to 20:30 h. However, in the context of community stability in the face of flowing spring-water, unresolved questions remain as regards how the native microorganisms counter the high flow rate and hold on to particular territories along the thermal gradient, what is the nature and extent of their mobility up and/or down the trajectory of the spring-water transit, and what is the net effect of influx, efflux and residence of microorganisms on the community structure.

Distributions of unique and shared genera across the sample-sites at 8:30 h (Fig. 5A; Supplementary Table S5) and 14:30 h (Fig. 5B; Supplementary Table S6) also illuminate potential mechanisms of bacterial diversity flux in this hydrothermal system. At both the sampling-hours, remarkably high number of genera was found to be present simultaneously in MTP, LTP and RVW: at 8:30 h, 72 out of the total 267 genera detected in the entire *Lotus Pond*-*Rulang* system; and at 14:30 h, 111 out of the total 316 genera detected in the system, were present at all these three sites. On the other hand, at 8:30 and 14:30 h, 36 and 41 genera were found to be simultaneously present in all four sample-sites respectively. These distributions, together with the presence of unique genera at both ends of the gradient (i.e. at HTP and RVW), and genera that were common and restricted to HTP and MTP, or LTP and RVW, insinuated that movement of bacteria, in this ecosystem, plausibly happens along the vent-to-river as well as river-to-vent trajectories throughout the day. While microbial transport in the vent-to-river trajectory is apparently driven by the hot-water flow, the reverse traffic potentially involves inching of microbes along the wobbly sub-aquatic sediments/sinters, if not swimming against the current of spring-water transit. Another mechanism of microbial flux in the river-to-vent trajectory could be washing of microbes into to the LTP from cooler zones near the banks of the outflow channel, which are connected to the river-water and not always agitated by the hot-water outflow. Supplementary Note 1 discusses additional features of spatiotemporal distribution of genera that suggest that influx of new bacteria from the vent as well as the river (into the hydrothermal territory) increase with the progress of the day.

**Fig. 5.**
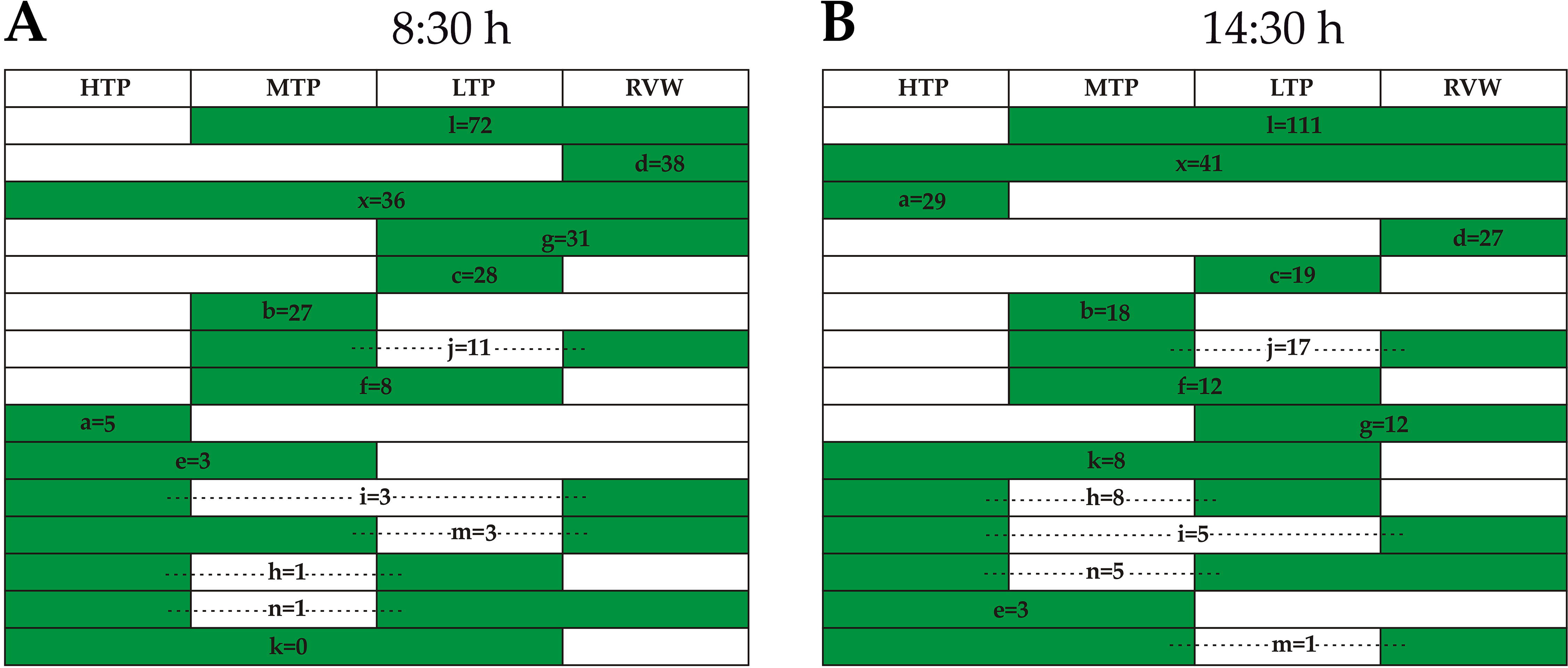
The numbers of genera that are unique to the different sites or common between the sites, across HTP, MTP, LTP and RVW, at (**A**) 8:30 and (**B**) 14:30 h. For both the sampling-hours, a = number of genera that were present only in HTP; b = number of genera that were present only in MTP; c = number of genera that were present only in LTP; d = number of genera that were present only in RVW; e = number of genera that were present in both HTP and MTP, but not elsewhere; f = number of genera that were present in both MTP and LTP but not elsewhere; g = number of genera that were present in both LTP and RVW, but not elsewhere; h = number of genera that were present in both HTP and LTP, but not elsewhere; i = number of genera that were present in both HTP and RVW, but not elsewhere; j = number of genera that were present in both MTP and RVW, but not elsewhere; k = number of genera that were present in HTP, MTP as well as LTP, but not in RVW; l = number of genera that were present in MTP, LTP as well as RVW, but not in HTP; m = number of genera that were present in HTP, MTP as well as RVW, but not in LTP; n = number of genera that were present in HTP, LTP as well as RVW, but not in MTP; x = number of genera that were present in all the four sample-sites.

Whatever may be the mechanism of microbial flux across the *Lotus Pond*-*Rulang* system, consistent presence of 28 bacterial genera at both HTP and MTP (irrespective of whether they were also present in LTP and/or RVW) across the forenoon-afternoon divide indicated that a considerable fraction of the *in situ* microbial diversity resided at high temperature sites for a considerable period of time. Likewise, another 103 genera were found to be present in at least one of the two ≥ 72°C sites (i.e. HTP or MTP) at 8:30 as well as 14:30 h, irrespective of whether they were also present in LTP and/or RVW (Supplementary Tables S5 and S6 show the sample-site-wise distributions of the total 267 and 316 genera identified in the *Lotus Pond-Rulang system* at 8:30 and 14:30 h respectively).

### 3.4. Sulfur speciation and diversity of chemolithotrophs along Lotus Pond’s spring-water transit

Incremental recharge of the hydrothermal reservoir with cold oxygenated water during the day-time could also be instrumental in the progressive increase of sulfate and concomitant decrease of dissolved sulfide, thiosulfate and sulfite in the vent-water between 6:30 and 17:30 h (Table 1). Consistent with this hypothesis, after sun-down (i.e. between 17:30 and 20:30 h), discharge-rate and sulfate concentration decreased, whereas temperature of the vent-water and concentrations of the reduced sulfur species increased (Table 1). Throughout the day, thiosulfate concentration in the vent-water was 4-5 times higher than that of the *in situ* sulfide. This indicated that thiosulfate originates in the deeper parts of the hydrothermal system, even as oxidation of dissolved sulfide to thiosulfate may also happen when the upward-moving deep hydrothermal water mixes with the more aerated snow-melted water in the shallower parts of the system. Furthermore, abiotic hydrolysis (Xu et al., 1998) and/or microbial disproportionation of elemental sulfur, which is abundant in the Puga geothermal system (Harinarayana et al., 2006), may also contribute to the formation of thiosulfate, sulfite and/or sulfate, in the shallower parts of the system. In this context it is remarkable that between 6:30 and 17:30 h, increase in sulfate and concomitant decrease in reduced sulfur species did not lower the pH of the vent-water; instead pH increased from 7.2 to 8.0. Diurnal build-up of acidity seems to be neutralized by the potential alkaline buffering conferred by the bicarbonate and borate salts abundant in all hydrothermal discharges of the Puga valley (Garrett, 1998).

Fluxes in pH and sulfur speciation along the spring-water’s transit are influenced primarily by aerial oxidation of the reduced sulfur compounds discharged from the vent; so, these fluctuations are different from the sub-surface-processes-driven fluxes recorded for the same parameters in the vent-water over the day (Table 1; Fig. 3C; Supplementary Table S2). In the vent-to-river trajectory, at both 8:30 and 14:30 h, there were significant increases in the concentrations of thiosulfate and sulfate up to LTP (both the sulfur-species, however, declined sharply in RVW); this was accompanied by steady decreases in dissolved sulfide, sulfite, and pH (sulfide and sulfite were no more there in LTP and RVW, at 8:30 h as well as 14:30 h). These trends are apparently attributable to the rapid oxidation of dissolved sulfide and sulfite to thiosulfate/sulfate and sulfate respectively, during transit from HTP to LTP. Notably, the dissolved sulfates precipitated from the HTP water-samples at 8:30 and 14:30 h had similar sulfur isotope ratios (δ^34^S values 16.3‰ and 16.5‰ VCDT respectively; see Supplementary Table S7). This, on one hand, reflected the steadiness of the biogeochemical processes related to sulfur cycling in the sub-surface, while on the other, suggested that sulfate supply is largely in excess of what is plausibly reduced microbially and/or precipitated epithermally from the upward-moving geothermal water. However, despite increases in sulfate concentrations from HTP to LTP, at both 8:30 and 14:30 h, δ^34^S values remained between +16.1‰ and +16.6‰ VCDT for the sulfates precipitated from MTP and LTP at either sampling-hour (Supplementary Table S7). Albeit we do not know the relative contributions of sulfide and sulfite in the increased sulfate concentrations along *Lotus Pond*’s spring-water transit, it, anyway, is evident from the above data that the cumulative sulfur isotope fractionations rendered by the two oxidative processes do not significantly alter the native sulfur isotope ratio of the vent-water sulfate. As observed for dissolved sulfate, sulfur isotope ratio of dissolved sulfide in the HTP water also remained consistent between 8:30 and 14:30 h (δ^34^S values 4.6‰ and 4.3‰ VCDT respectively), thereby reiterating the steadiness of the biogeochemical sulfur cycle in the vent’s sub-surface. In contrast, however, drop in the sulfide concentration of the spring-water in transit from HTP to MTP (followed by complete extinction in LTP), at both the sampling-hours, and concomitant enrichment of ^34^S in the dissolved sulfide (ΔS^2−^_MTP-HTP_ was 2.4‰ and 1‰ at 8:30 and 14:30 h respectively; see Supplementary Table S7), were consistent with the potential oxidation of sulfide during this passage. Furthermore, in this context, it is noteworthy that the sulfates precipitated from RVW had lower δ^34^S values (12.4‰ and 13.1‰ VCDT at 8:30 and 14:30 h respectively; see Supplementary Table S7) than their HTP, MTP and LTP counterparts. This indicated that the sulfate-source and/or sulfur biogeochemistry of the river are different from those of the hot spring system.

Variations in the concentrations of sulfur species along *Lotus Pond*’s spring-water transit indicated that opportunities for chemolithotrophic life were aligned anti-parallel to the hydrothermal gradient offering resistance to microbial colonization. Concurrently, 15-33% of the genera detected at the individual sample-sites, at any sampling-hour, have known sulfur-chemolithotrophic members (Supplementary Table S7; also refer to Supplementary Tables S5 and S6 for all the known sulfur-metabolizing attributes of the genera identified across *Lotus Pond*’s sample-sites at 8:30 and 14:30 h respectively). Sulfur-oxidizing genus count at a sample-site, when considered as a percentage of the total number genera present, was highest for HTP, at both 8:30 h (33% of total genus-count) and 14:30 h (20% of total genus-count) (Fig. 3C; Supplementary Table S7). This parameter was remarkably comparable for MTP, LTP and RVW across the two sampling-hours (15-18% of the genera detected individually in these samples have known sulfur-oxidizing members).

In contrast to the rich diversity of sulfur-oxidizing bacteria present along *Lotus Pond*’s hydrothermal gradient, very few genera detected at individual sample-sites, at either sampling-hour of the day, encompass species/strains capable of reducing S_2_O_3_^2−^, SO_3_^2−^ and/or SO_4_^2−^, according to published literature. Whereas the numbers of sulfur-reducing genera present were only 0 and 2 for HTP at 8:30 and 14:30 h respectively, the same count ranged between 8 and 12 (i.e. 6-8% of the total number genera detected *in situ*) for MTP, LTP and RVW, at either sampling-hour of the day (Fig. 3C; Supplementary Table S7). It is difficult to elucidate why specific types of microorganisms often congregate in particular hot springs. Even as hydrothermal ecosystems, under influence of several extraneous factors, are always in a dynamic state, niche selection may drive localized assembly of specific microbial types (Power et al., 2018).

### 3.5. Predominance of mesophiles in Lotus Pond’s high-temperature sites

Compositions of the aquatic microbial communities revealed from all the four sample-sites, at both 8:30 and 14:30 h, were remarkably dominated by such bacterial taxa, members of which are incapable of laboratory growth at > 45°C, according to published pure culture studies. Out of the 57 genera that were detected exclusively at HTP and/or MTP at 8:30 and/or 14:30 h (Supplementary Tables S5 and S6), 41 have no report of laboratory growth at > 45°C, 11 are known to grow *in vitro* at maximum temperatures ranging between 48-60°C, whereas 5 do not have any information on their upper temperature-limit for laboratory growth (see Supplementary Table S8 and references therein). So, every genus confined to HTP and/or MTP was actually present at *in situ* temperature(s) surpassing the limits established for their growth/survival *in vitro*.

The overall mesophile-count in the 70-85°C temperature-zone of *Lotus Pond* becomes much higher if we consider the genera that were present at LTP and/or RVW in addition to being present at HTP and/or MTP. For instance, out of the total 52 genera that were there in HTP at 8:30 h (47 of these were present in MTP, LTP and/or RVW as well) (Fig. 5A; Supplementary Table S5), only 3 (namely *Caldicellulosiruptor*, *Fervidobacterium* and *Hydrogenobacter*) have reports of laboratory growth at > 80°C; 13 are known to grow in the laboratory at > 45°C, but not at > 80°C; 31 have no report of laboratory growth at > 45°C (refer to Supplementary Table S9 and references therein); while 5 out of the 52 genera are not there in the list of validly-published taxa (http://www.bacterio.net/) (Parte, 2013) and so are not included in Supplementary Table S9. Again, out of the total 100 genera that were there in HTP at 14:30 h [71 of these were present in MTP, LTP and/or RVW as well (Fig. 5B; Supplementary Table S6)], only 3 (*Caldicellulosiruptor*, *Fervidobacterium* and *Hydrogenobacter*) have reports of laboratory growth at > 80°C; 22 are known to grow at > 45°C, but not at > 80°C, *in vitro*; 63 have no report of laboratory growth at > 45°C; and there is no information in the literature regarding the upper temperature-limit of laboratory growth for the remaining 12 (refer to Supplementary Table S10 and references therein).

The above data were consistent with the findings of the 2013 exploration of the HTP site where 84 out of the 107 genera identified in metataxonomic analysis were found to have no report of laboratory growth above 45°C (Roy et al., 2016). Consistent prevalence of mesophiles in *Lotus Pond*’s HTP sites was also consistent with the findings of an environmental 16S rRNA gene cloning and sequencing-based analysis of the *in situ* bacterial diversity undertaken following an expedition conducted in August 2009. The 50 environmental 16S rRNA gene fragment clones constructed and sequenced in that study using Bacteria-specific universal primers (Gerhardt et al., 1994), had yielded 17 species-level clusters unified at the level 97% 16S rRNA gene sequence similarity. Of the 19 type strains of established bacterial species with which the representative clones of the 17 species-level clusters had maximum sequence similarities, only one have laboratory growth reported at 50°C; all the others have their upper limit of temperature for *in vitro* growth at ≤ 45°C (refer to Supplementary Table S11 for the summary details of the 16S rRNA gene cloning and sequencing-based analysis conducted in 2009).

### 3.6. Potential geochemical determinants of high habitability

The habitability of *Lotus Pond*, like any other hot spring, is influenced by the macromolecule-disordering (chaotropic) activities of high temperature that constraints the functionality of cell systems in the same entropic way as chemical chaotropic agents do, i.e., by reducing intracellular water activity, decreasing hydrogen bonding and electrostatic interactions of hydrated biomacromolecules, and in the process bringing about bioenergetically-costly cellular stress responses that eventually forces metabolism to collapse (Koynova et al., 1997; Hallsworth et al., 2003; Bhaganna et al., 2010; Chin et al., 2010; Ball and Hallsworth, 2015; Cray et al., 2015). While trying to find the potential biophysical basis of high bacterial diversity and mesophile predominance in the high-temperature sites of *Lotus Pond* we observed that the HTP and MTP sample-sites have high *in situ* concentrations of boron, lithium, sodium, sulfide, thiosulfate and sulfate (Fig. 3A-B; Supplementary Table S2), all of which are kosmotropic, and can therefore stabilize, structure, and impart rigidity to biomacromolecules (Brown, 1990; Koynova et al., 1997; Wiggins, 2001; Hribar et al., 2002; Mancinelli et al., 2007), and in doing so mitigate against the disruption of cellular systems by chaotropic agents (Hallsworth et al., 2007; Chin et al., 2010; Cray et al., 2013, 2015; de Lima Alves et al., 2015; Yakimov et al., 2015). Furthermore, this assumption was buttressed by the fact from HTP to MTP, at both 8:30 and 14:30 h, the steep rise in diversity concomitant with the little drops in temperature coincided with small but definite increases in the concentrations of boron, thiosulfate and sulfate (apparently due to a combination of factors such as aerial oxidation of the reduced states, lower flow rate of the spring-water over the sintered terraces of the apron, and desorption from the subaquatic mineral sinters); lithium also increased from HTP to MTP at 8:30 h but remained almost unchanged at 14:30; sodium concentration remained almost unchanged between HTP and MTP at both the sampling-hours. In this context it is further noteworthy that the geomicrobiological scenario of high microbial diversity at a high-entropy environment, as revealed here for *Lotus Pond*, is analogous to that reported for chaotropic, 2.50-3.03 M MgCl_2_-containing deep sea brines, where habitability higher than that of 2.2-2.4 M MgCl_2_ brines is facilitated by the co-occurrence of kosmotropic sodium and sulfate ions that act to stabilize biomacromolecules and lower the thermodynamic cost of maintaining microbial cell systems (Hallsworth et al., 2007; Yakimov et al., 2015). All these considerations collectively prompted us to hypothesize that *in situ* abundance of kosmotropic solutes could confer high habitability to the high-temperature waters of *Lotus Pond* in the same biophysical way as they do in other comparator chaotropic environments such as hypersaline brines. Consequently, we tested this hypothesis via high temperature incubations of the pure culture of a mesophilic *Lotus Pond* isolate, in the presence/absence of the kosmotropes identified above.

### 3.7. *Isolation and characterization of* Paracoccus *SMMA_5, the model organism used to test the effects of solutes on cell survival at elevated temperatures*

When two individual sets of residues obtained from the filtration of two 1000-mL batches of HTP-water were enriched separately in MST (modified basal and mineral salts solution supplemented with thiosulfate as the sole source of energy and electrons) and Reasoner’s 2A (R2A) broth media at 37°C, 10^6^ and 10^9^ CFUs mL^−1^ broth culture were obtained after 14 h of incubations respectively. Lag phase of both the mixed-cultures was recorded to be approximately 2 h. These cellular yields for the chemolithoautotrophic MST- and chemoorganoheterotrophic R2A-based enrichment cultures corresponded to an oxidation of 10 mM thiosulfate to an equivalent amount of sulfate (both equal to 20 mM sulfur) and OD_600_ of 0.8, respectively. The generation times (of the two different mixed-cell populations) 65 and 60 min, determined from the above data for the MST- and R2A-broth-enrichments respectively, were used to calculate the potential numbers of live chemolithotrophic and chemoorganotrophic mesophiles in 1000-mL HTP-water as 1.5 x 10^5^ and 7.5 x 10^7^ cells respectively. These two numbers, in conjunction with the small time-span within which high CFU numbers were reached mL^−1^ of the enrichment cultures, indicated that the mesophile populations of the HTP water-sample not only had high cell-counts, but were also metabolically so healthy as to readily divide upon withdrawal of thermal stress.

From the two mixed-culture consortia, several mesophilic bacterial strains were isolated (at 37°C) via dilution plating and iterative steps of dilution streaking on corresponding agar plates. Of the several tens of pure bacterial strains isolated in this way, a cluster of 12 facultatively sulfur-chemolithoautotrophic strains obtained from the MST-based isolation series were found to be taxonomically closest to members of the genus *Paracoccus* of the family *Rhodobacteraceae* (order *Rhodobacterales*) of class *Alphaproteobacteria*, and the strain SMMA_5 was the representative of this cluster. The 16S rRNA gene sequence of SMMA_5 (GenBank accession number LN869532) exhibited 97% similarity with strains belonging to *Paracoccus versutus*, *Paracoccus huijuniae*, *Paracoccus aminovorans*, *Paracoccus bengalensis*, *Paracoccus aestuariivivens* and several other *Paracoccus* species. In this context it is noteworthy that OTUs affiliated to the genus *Paracoccus* were also detected in all the four sample-sites at all the sampling-hours, except for LTP at 8:30 h (Supplementary Tables S5 and S6). SMMA_5 was deposited to the Microbial Type Culture Collection and Gene Bank (MTCC), Chandigarh, India with the public accession number MTCC 12601.

SMMA_5 exhibited similar cellular growth yields after 12 h incubation at 37°C, 40°C and 45°C in the chemolithoautotrophic MST medium. At all the three temperatures, after 12 h incubation, > 100-fold increases were recorded in the number of colony-forming units (CFUs) present, with respect to the initial CFU-counts. In contrast, 12 h incubation in MST at 50°C left only ~2% of the initial CFU-counts as ready to divide and capable of forming single colonies; 12 h at 60 or 70°C left no CFU in ready-to-divide state (Supplementary Fig. S2).

Generation time (g) in MST at 37°C was determined by putting the values of CFU-count mL^−1^ culture obtained at the start-point (30 min) and end point (12 h) of the first exponential phase (i.e. 1.6 x 10^6^ and 3.6 x 10^8^ respectively; see Supplementary Fig. S3) to Equation 1 (Schlegel and Zaborosch, 1993), where t denotes the time duration of the exponential phase, N stands for cell number after time t, and N_0_ denotes cell number at the beginning of the exponential phase. Notably, the lag phage of SMMA_5 growth in MST medium at 37°C (Supplementary Fig. S3) lasts for only the first 30 min of incubation. Subsequent to this, exponential growth occurs for the next 11 h 30 min, i.e. upto the 12^th^ hour of incubation. Albeit growth continues further, till the 18^th^ hour of incubation, the second 6 h phase has a lower rate, consequent to which the growth curve (still a straight line) assumes a different slope. Generation time (88 min) was caculated based on the first exponential phase (i.e. 30 min to 12 h) because this datum was to be discussed only in the context of what happens to SMMA_5 cultures within 50 min of exposure to heat.

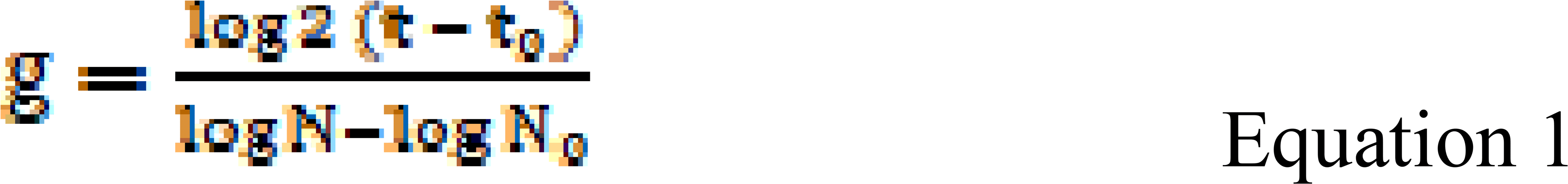

### 3.8. Kosmotropic solutes increase the proportion of ready-to-divide cells in cultures at elevated temperatures

In order to check whether the kosmotropic solutes present in *Lotus Pond* can mitigate against heat *in vitro*, the effects of Li^+^ (1.0 mM), B_4_O_7_^2−^ (4.0 mM; equivalent to 16.0 mM B), SO_4_^2−^ (1.0 mM) and Na^+^ (17.0 mM) were tested on the high-temperature growth/survival of the mesophilic HTP-isolate *Paracoccus* strain SMMA_5. Kosmotrope concentrations used in these experiments were comparable to the concentration-ranges recorded for lithium, boron, sulfate and sodium in the HTP-MTP territory (Supplementary Table S2). Whereas the cellular growth yield of SMMA_5 in MST at 37°C corresponded to a generation time of approximately 88 min, incubation for only 50 min in MST at 70°C reduced CFU-count to ~2% of the initial level (i.e. the CFU-count recorded at 0 h of incubation) (Fig. 6A-B). However, after 50 min at 70°C in MST-Li, MST-B_4_O_7_, MST-SO_4_ or MST-Na approximately 21%, 1%, 3% and 8% CFUs of SMMA_5 remained ready to divide and form single colonies respectively (Fig. 6A-B). On the other hand, at 50°C, while CFU-count remained unchanged after 50 min incubation in MST, the same more than doubled from the 0 h levels in MST-Li, MST-B_4_O_7_, MST-SO_4_ and MST-Na cultures (Fig. 6C-D). In this context it is noteworthy that when incubation at 50°C was extended to 12 h, the MST-Li, MST-B_4_O_7_, MST-SO_4_ and MST-Na cultures had approximately 30%, 14%, 11% or 2% of the initial CFU-count in ready-to-divide state respectively, as compared to ~2% in MST (Supplementary Fig. S4C-D). In contrast, after 12 h at 70°C, no CFUs were left as the MST as well as the kosmotrope-supplemented-MST cultures (Supplementary Fig. S4A-B). The above results reflected that the effects of kosmotropic ions on the proportion of microbial cells remaining ready-to-divide depend on the time-span of exposure to high temperatures. Incubations at 37°C, for both 50 min (Fig. 6E-F) and 12 h (Supplementary Fig. S4E-F), showed none of the kosmotropic solutes to be inhibitory to SMMA_5; rather they were found to be stimulatory to growth.

**Fig. 6.**
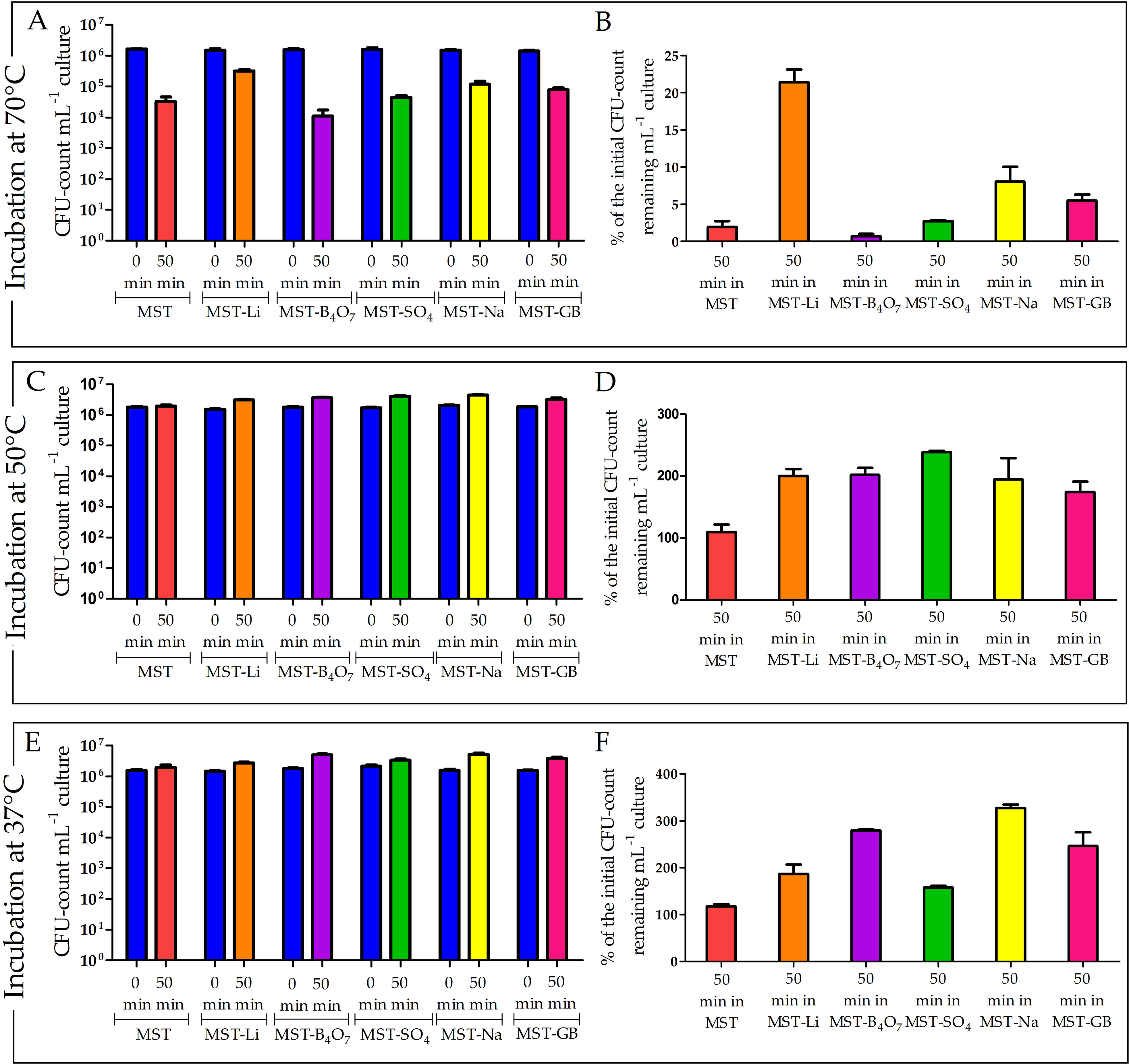
CFU-counts recorded mL^−1^ culture at 0 and 50 min, and percentages of the initial CFU-counts that remained ready-to-divide after incubating SMMA_5 for 50 min, in different variants of MST at 70°C, 50°C and 37°C: (**A**, **C**and **E**) CFU-counts recorded mL^−1^ culture in MST, or MST supplemented with salts of Li^+^ (MST-Li), B_4_O_7_^2−^ (MST-B_4_O_7_), SO_4_^2−^ (MST-SO_4_), Na^+^ (MST-Na) or glycine-betaine (MST-GB), at 0 and 50 min of incubation at 70°C, 50°C and 37°C respectively; (**B**, **D**and **F**) percentages of the initial CFU-counts that remained ready-to-divide in MST, MST-Li, B_4_O_7_^2−^, MST-SO_4_, MST-Na or MST-GB, after 50 min incubation at 70°C, 50°C and 37°C respectively. In (**A**, **C**and **E**), all 0 h CFU-counts, irrespective of the test medium and incubation temperature, are represented by blue bars; CFU-count recorded after 50 min incubations in MST, MST-Li, MST-B_4_O_7_, MST-SO_4_, MST-Na or MST-GB are represented by red, orange, violate, green, yellow or pink bars respectively. In (**B**, **D**and **F**), percentages of the initial CFU-counts that remained ready-to-divide in MST, MST-Li, MST-B_4_O_7_^2−^, MST-SO_4_, MST-Na or MST-GB, after 50 min incubation, irrespective of the incubation temperature, are represented by red, orange, violate, green, yellow or pink bars respectively. All the data shown in this figure are means for three different experiments; error bars indicate the standard deviations.

Among the four kosmotropic solutes tested, the alkali metal lithium was found to be the most thermoprotective at both 50°C and 70°C. This finding is consistent with previous studies where *Listeria monocytogenes* heated in raw milk at 62.8°C for 10, 15, and 20 min was recovered after 48, 96 and 144 h respectively, by adding 7 g L^−1^ LiCl to the revival culture incubated at 30°C in rich heterotrophic medium (Mendonca and Knabel, 1994). Thermoprotective effect of lithium could be linked, but not necessarily restricted, to its activities at the periphery of cells. Lithium readily ionizes to Li^+^ in aqueous milieu, and efficiently forms hydrogen bonds with nearby water molecules, as well as charged surfaces such as cell membranes, thereby modifying their intrinsic van der Waals, and electrostatic, forces (Roark et al., 2012). In doing so, lithium can not only stabilize the biomacromolecules it interacts with but also enhance the permeability of those such as cellular membranes, which in turn can enhance the entry of other thermoprotective kosmotropes such as sodium (Hesketh et al., 1978). This potential mechanism - which may also facilitate the import of potential compatible solutes present in the aqueous milieu - is consistent with previous reports showing that *Escherichia coli*, under the action of 2.5% LiCl, produces spheroplasts that can multiply by binary fission, have high osmotic stability, and can revert to normal bacilli forms when transferred to lithium-free environment (Pitzurra and Szybalski, 1959).

### 3.9. Glycine-betaine as a native thermoprotectant for the Lotus Pond microflora

Large number of OTUs identified across the *Lotus Pond* sample-sites were affiliated to the genera *Bacillus*, *Ectothiorhodospira*, *Escherichia*, *Halomonas*, *Pseudomonas*, *Staphylococcus*, etc., members of which are proficient producers of the osmoprotective compatible solute glycine-betaine or N,N,N trimethylglycine (C_5_H_11_NO_2_) (Galinski and Troper, 1982; Lamark et al., 1992; Boch et al., 1996; Rosenstein et al., 1999; Canovas et al., 2000; Sleator and Hill, 2002). SMMA_5, after 12 h incubation in MST at 50°C, also released glycine-betaine in the spent medium at a concentration of 4.4 mM.

Glycine-betaine is taken-up/synthesized in large amounts by halophilic/halotolerant microbes as their secondary response to external salt concentrations > 0.5 M NaCl (Sleator and Hill, 2002). Compatible solutes in general are highly soluble organic molecules that have no net charge at physiological pH (Galinski, 1995), so they can be accumulated at even > 1 M intracellular concentration without interfering with the functionalities of biomacromolecules such as DNA or protein (Brown, 1976; Strøm and Kaasen, 1993; Galinski and Tròper, 1994; Record Jr et al., 1998). Besides acting as osmotic balancers, compatible solutes impart a general stabilizing effect on biomacromolecules (Brown, 1976; Arakawa and Timasheff, 1985; Colaco et al., 1992; Qu et al., 1998). Preferential exclusion of compatible solutes from the immediate hydration sphere of proteins leads to a non-homogeneous distribution of the solute within the cell-water, thereby causing a thermodynamic disequilibrium (Welsh, 2000). This disequilibrium is minimized via reduction in the volume of cell-water from which the solute is excluded; this, in turn, is achieved by reduction in the surface-area/volume of the proteins through increases in sub-unit assembly and stabilization of secondarily-folded tertiary structures (Welsh, 2000). All such biophysical maneuvers at the molecular-level potentially add-up at the level of the organism, to confer tolerance against a variety of chaotropic stressors such as high salt, desiccation and heat (Lippert and Galinski, 1992; Welsh, 2000).

From the results of the growth experiments conducted in MST medium it was evident that the 4.4 mM glycine-betaine produced indigenously by SMMA_5 was not sufficient to arrest declines in CFU-count at temperatures ≥ 50°C. However, this trimethylammonium compound, when supplied extraneously at a concentration of 10 mM, did so at both 50°C and 70°C. Whereas incubation for only 50 min in MST at 70°C reduced CFU-count to ~2% of the initial level (Fig. 6A-B), after 50 min at 70°C in MST-GB approximately 6% CFUs of SMMA_5 remained ready to divide and form single colonies respectively (Fig. 6A-B). On the other hand, at 50°C, while CFU-count remained unchanged after 50 min incubation in MST, the same almost doubled from the 0 h levels in the MST-GB culture (Fig. 6C-D). When incubation-time was extended to 12 h, in MST-GB, at 50°C, growth equivalent to an approximately 5.3-fold increase from the initial CFU-count was recorded; in contrast, only ~2% of the initial CFU-count had remained ready to divide in MST (Supplementary Fig. S4C-D). In contrast, after 12 h at 70°C, no CFUs were left as the MST or MST-GB cultures (Supplementary Fig. S4A-B). This indicated that the effect of glycine-betaine on the divisibility of cells, much like that of kosmotropic solutes, depends on the time-span of exposure to high temperatures. Incubations at 37°C, for both 50 min (Fig. 6E-F) and 12 h (Supplementary Fig. S4E-F), showed that glycine-betaine was not inhibitory to SMMA_5; rather it was found to be stimulatory to growth.

### 3.10. At high temperature, effect of a kosmotropic/compatible solute is more on cell viability than divisibility

Effects of Li^+^ (1.0 mM), B_4_O_7_^2−^ (4.0 mM; equivalent to 16.0 mM B), SO_4_^2−^ (1.0 mM), Na^+^ (17.0 mM), or C_5_H_11_NO_2_ (10.0 mM), on the viability (or the metabolically-active state) of cells incubated at 70°C for 50 min was tested by assessing their ability to accumulate the non-toxic and non-fluorescent molecule fluorescein diacetate (FDA), and hydrolyze it to fluorescein, which in turn was detected via flow cytometry (Wieder, 1999). After incubation in MST for 50 min at 70°C, approximately 15% of the 10,000 SMMA_5 cells taken randomly from the culture were found to remain viable (i.e., capable of taking FDA stain) (Fig. 7A); this, remarkably, was 7.5 times higher than what percentage of initial CFU-count had remained ready to divide under the same culture condition (Fig. 6B). On the other hand, after incubation in MST-Li, MST-B_4_O_7_, MST-SO_4_, MST-Na or MST-GB for 50 min at 70°C, approximately 71%, 43%, 55%, 64% and 55% of the 10,000-cell samples taken randomly from the corresponding SMMA_5 cultures were found to remain viable respectively (Fig. 7B-F). These numbers were not only 3-5 times higher than the 15% cell-viability observed in MST (Fig. 7A), the viable-cell percentages recorded in MST-Li, MST-B_4_O_7_, MST-SO_4_, MST-Na or MST-GB after 50 min at 70°C (Fig. 7B-F) were approximately 3, 43, 18, 8 and 9 times higher respectively than the corresponding percentages of initial CFU-counts that had remained ready to divide in the growth experiments under the same five culture conditions (Fig. 6B).

**Fig. 7.**
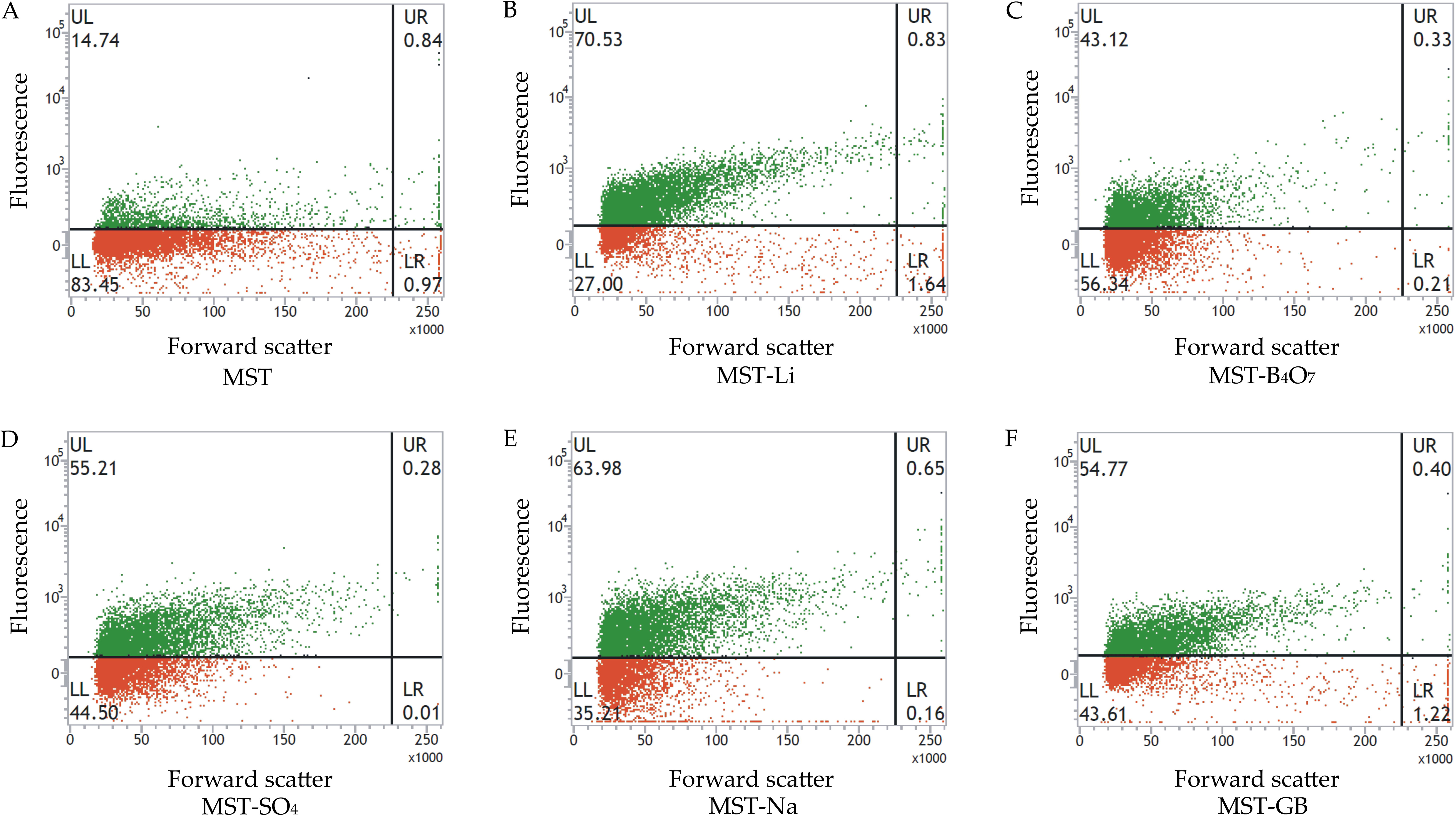
Flow-cytometry-based dot plots showing the proportions of FDA-stained and FDA-unstained cells in different SMMA_5 cultures incubated at 70°C for 50 min in (**A**) MST, (**B**) MST-Li, (**C**) MST-B_4_O_7_, (**D**) MST-SO_4_, (**E**) MST-Na, or (**F**) MST-GB medium. In each figure, fluorescence versus forward scatter data for 10000 randomly-taken cells have been potted; green and red dots represent cells that were stained and not stained by FDA respectively.

Comparison between the results of flow-cytometry-based cell-viability assays and CFU-count-based growth data pointed out that at 70°C, the effect of a kosmotrope/compatible solute on cell viability is more than its effect on the ability of cells to divide readily and form single-colonies. Consequently, questions arose with regard to the physiological basis of the wide difference observed between the percentage of cells remaining ready to divide and that remaining viable in all the experimental cultures. Furthermore, it was imperative to check whether all individuals within the large populations of viable cells detected in the individual experimental cultures were capable of dividing at some later stage. Remarkably, when all the CFU-count-based experiments were repeated by prolonging the 37°C-incubation of the LB-agar plates to 10 days, many more colonies appeared after 36 h in a staggered manner, thereby revising the final CFU-counts of the individual experimental cultures up to levels comparable with the corresponding viable-cell percentages determined by flow cytometry.

## 4. Conclusions

The geomicrobial characteristics of *Lotus Pond* suggested, and the results of our pure-culture-based growth experiments corroborated, that under variable levels of heat and exposure-time, kosmotropes and compatible solutes can enhance the divisibility, as well as the intrinsic viability, of mesophilic microbial cells up to different extents. This finding ushers a new biophysical baseline that can determine the habitability of hydrothermal ecosystems. Albeit the experimental evidences provided here are based on one pure culture isolate, it is noteworthy that in almost all occasions when unknown biological phenomena are revealed, a model organism is studied extensively to understand the process, with the expectation that discoveries made in the model organism will provide insight into the workings of other organisms (Fields and Johnston, 2005). This said, there are other reasons as well to believe that the present evidences obtained from SMMA_5 are not one-off phenomena, and that *Lotus Pond*’s microbial communities do adapt collectively to the spatiotemporal changes in their ambient temperature. Recent studies have shown that corals can adapt to their thermal environment and inherit heat tolerance across generations, apparently with the help of their symbiotic microflora that have the capacity for more rapid adaptation (Ziegler et al., 2017). Furthermore, the geobiological linkages drawn between the ecosystem constraints and the unique geochemistry of *Lotus Pond* on one hand, and its mesophiles-dominated microbiome on the other, stand up (in the light of the thermodynamic logic used) to propound, by themselves, the kosmotropes-mediated survival of mesophiles in high-temperature habitats. This is because the biophysical (thermodynamic) basis envisaged for *Lotus Pond*’s peculiar microbiome architecture emanates from comparisons with entropically-similar microbial habitats, and various cited studies which have already demonstrated that this metabolic response to the thermodynamic status of the solutes is universal (cited papers that demonstrate the mitigation of chaotropic-stress-induced cell-system failure via kosmotropic-solutes-mediated entropy-minimization and stabilization of biomacromolecules include: Hallsworth et al., 2007; Bhaganna et al., 2010; Chin et al., 2010; Cray et al., 2013, 2015; de Lima Alves et al., 2015; Yakimov et al., 2015).

Hydrothermal spring waters are likely to get contaminated with mesophilic bacteria from various surface/sub-surface sources (Power et al., 2018; Colman et al., 2019), and while such contaminants are likely to be detected in environmental DNA sequencing-based investigations, many of them may also withstand the thermal stress which they have been thrust into (Xie et al., 2011; Alsop et al., 2014; Cowan et al., 2015). In this context, simultaneous detection of the same mesophilic entities in the culture-independent and culture-dependent investigations of a hydrothermal sample - as has happened in the present study, as well as in a previous investigation of Indonesian hot springs (Baker et al., 2001) - vouches for not only the existence of the mesophiles, but also their survival, *in-situ*. However, unlike for the present study, no geochemical factors was pointed out as instrumental behind the high temperature *in vitro* growth of the isolates from Indonesian hot springs (Baker et al., 2001); yet, our close scrutiny of the media in which the Indonesian isolates belonging to the *Xanthomonas*-*Frateuria* cluster of *Gammaproteobacteria* were grown at 70-85°C, suggested that the high sodium, sulfate, or some unknown ingredient(s) of the trace metals solution used in the media, may had aided the isolates belonging to taxa not normally associated with thermophilicity, grow at high temperatures *in vitro*.

Growth experiments carried out at high temperatures with the present mesophilic isolate SMMA_5 afforded two remarkable findings: (i) in the presence as well as absence of kosmotropes/compatible solutes, greater proportions of cells remain viable than what remain ready-to-divide immediately upon withdrawal of heat, and (ii) both types of thermoprotective substances cause more pronounced increases in viable cell-count than in the number of cells that are ready to divide and form colonies upon withdrawal of heat. These results indicate that to mitigate the entropy-increasing effects of heat, microbial cells indigenously deploy such entropy-minimizing and macromolecule-stabilizing biophysical resources and mechanisms that lower the energy-cost of cell-system maintenance, and in the process trade-off cell-division for cell-viability [these biophysical mechanisms, according to literature, may involve facilitation of ion pairs, hydrogen bonds, hydrophobic interactions, disulfide bridges, packing and inter-subunit interactions, decrease of the entropy of unfolding/disintegration, etc (Vieille and Zeikus, 2001)]. Extraneously available kosmotropes and compatible solutes augment these mechanisms in such a way as to increase the proportion of viable cells more than that of ready-to-divide cells, within the population. As maintenance of appropriate balance between the structural stability (integrity needed to avoid denaturation) and flexibility (necessary for proper functioning) of biomacromolecules is the key to survival/growth at high temperatures (Fields, 2001), all entropy-minimizing biophysical mechanisms do have their limitations with respect to the temperature-levels and exposure-times over which they can enhance survival/growth. Furthermore, all individuals within a cell-population are not equivalently benefitted from the macromolecule-stabilizing effects of the kosmotropes or compatible solutes plausibly due to their differential metabolic states at the time of introduction to heat. Precise delineation of how microbial populations in open-ended hydrothermal ecosystems as *Lotus Pond* subsist and/or proliferate through high temperature over time and space require comprehensive revelation of the entire gamut of enabling factors (such as kosmotropes and compatible solutes) that are present *in situ*. This, in turn, require further holistic knowledge on the inorganic and organic geochemistry of the system, followed-up by culture-dependent studies involving combinatorial use of the enabling factors alongside the time-variable. The prospective knowledge thus obtained would have immense potential for getting translated into a broad-based, habitat-inspired, laboratory protocol for culturing mesophilic microorganisms at high temperatures. Much like other ongoing efforts to improve thermotolerance using selection pressure (Zhou et al., 2019), potential formulation of such culture conditions, which impart thermotolerance to industrially/economically useful microorganisms without compromising on their metabolic activities, can revolutionize the field of process biotechnology and fermentation.

## Author’s contribution

WG initiated and developed the work, designed the experiments, interpreted all the results, and wrote the paper in conjunction with RC who made critical intellectual contributions to the paper. NM, CR, MA, SM, SB, MJR, PKH, NN and TB carried out the studies of microbiology. AP, SF, SPV and AM carried out the studies of geochemistry. CR, TM, AP, NM and AM participated in data analysis. WG, CR, PKH and MJR carried out the fieldworks. All authors read and vetted the manuscript.

## Acknowledgements

This research was financed by Bose Institute as well as Science and Engineering Research Board (SERB), GoI (SERB grant numbers were SR/FT/LS-204/2009 and EMR/2016/002703). Sri Pankaj Kumar Ghosh (Chinsurah, West Bengal, India) provided additional travel grants philanthropically. NM received fellowship from Science and Engineering Research Board, Government of India (GoI). CR and MJR received fellowships from University Grants Commission, GoI. MA and SB received a post-doctoral and doctoral fellowship from Bose Institute respectively. SM received fellowship from Department of Science and Technology, GoI. On field technical assistance provided by Baishali Ghosh, Asgar Ali, Bikash Jana, Rimjhim Bhattacherjee, Srabana Bhattacherjee, Lotus Sonam and Amrit Pal Singh, over the decade long exploration of Puga geothermal area, is gratefully acknowledged. Useful discussions were provided by John Edward Hallsworth (Queen’s University Belfast, Northern Ireland), Subhra Kanti Mukhopadhyay and Ambarish Mukherjee (University of Burdwan, India), Johann Peter Gogarten, R. Thane Papke and Daniel Gage (University of Connecticut, CT, USA), Ashish Kumar Nanda (University of North Bengal, India), Asoke Chandra Ghose (Bose Institute, India) and Ranjan Kumar Nandy (National Institute of Cholera and Enteric Diseases, India). We are also thankful to our friends Ashish Kumar George, Pravin Nilawe, Atima Agarwal, Neeraj Kumar Chauhan and Disha Banerjee (Invitrogen BioServices India Pvt Ltd) for regularly tutoring us on next generation DNA sequencing and data analysis.

## Appendix A. Supplementary materials

A single MS Word file named Mondal et al_Supplementary_Materials, which contains 10 Supplementary Tables and their associated References, plus one Supplementary Note and four Supplementary Figures.

## Appendix B. Supplementary materials

A single MS Excel file named Mondal et al_Dataset 1, which contains one Supplementary Table.

## References

Alam, M., Pyne, P., Mazumdar, A., Peketi, A., Ghosh, W., 2013. Kinetic enrichment of 34S during proteobacterial thiosulfate oxidation and the conserved role of SoxB in S-S bond breaking. Appl. Environ. Microbiol. 79, 4455–4464. https://doi.org/10.1128/AEM.00956-13.

Alsop, E.B., Boyd, E.S., Raymond, J., 2014. Merging metagenomics and geochemistry reveals environmental controls on biological diversity and evolution. BMC. Ecol. 14, 16. https://doi.org/10.1186/1472-6785-14-16.

Alves, M.P., Rainey, F.A., Nobre, M.F., da Costa, M.S., 2003. *Thermomonas hydrothermalis* sp. nov., a new slightly thermophilic gammaproteobacterium isolated from a hot spring in central portugal. Syst. Appl. Microbiol. 26, 70–75. https://doi.org/10.1078/072320203322337335.

Arakawa, T., Timasheff, S.N., 1985. The stabilization of proteins by osmolytes. Biophys. J. 47, 411–414. https://doi.org/10.1016/S0006-3495(85)83932-1.

Baker, G.C., Gaffar, S., Cowan, D.A., Suharto, A.R., 2001. Bacterial community analysis of Indonesian hot springs. FEMS. Microbiol. Lett. 200, 103–109. https://doi.org/10.1111/j.1574-6968.2001.tb10700.x.

Ball, P., Hallsworth, J.E., 2015. Water structure and chaotropicity: their uses, abuses and biological implications. Phys. Chem. Chem. Phys. 17, 8297–8305. https://doi.org/10.1039/C4CP04564E.

Battin, T.J., 1997. Assessment of fluorescein diacetate hydrolysis as a measure of total esterase activity in natural stream sediment biofilms. Sci. Total. Environ. 198, 51–60. https://doi.org/10.1016/S0048-9697(97)05441-7.

Berezovsky, I.N., Shakhnovich, E.I., 2005. Physics and evolution of thermophilic adaptation. Proc. Natl. Acad. Sci. USA. 102, 12742–12747. https://doi.org/10.1073/pnas.0503890102.

Bhaganna, P., Volkers, R.J., Bell, A.N., Kluge, K., Timson, D.J., McGrath, J.W., et al., 2010. Hydrophobic substances induce water stress in microbial cells. Microb. Biotechnol. 3, 701–716. https://doi.org/10.1111/j.1751-7915.2010.00203.x.

Boch, J., Kempf, B., Schmid, R., Bremer, E., 1996. Synthesis of the osmoprotectant glycine betaine in *Bacillus subtilis*: characterization of the *gbsAB* genes. J. Bacteriol. 178, 5121–5129. https://doi.org/10.1128/jb.178.17.5121-5129.1996.

Brown, A.D., 1976. Microbial water stress. Bacteriol. Rev. 40, 803–846.

Brown, A.D., 1990. Microbial water stress physiology. Chichester: John Wiley and Sons.

Canovas, D., Vargas, C., Kneip, S., Moron, M.J., Ventosa, A., Bremer. E., et al., 2000. Genes for the synthesis of the osmoprotectant glycine betaine from choline in the moderately halophilic bacterium *Halomonas elongata* DSM 3043. Microbiology. 146, 455–463. https://doi.org/10.1099/00221287-146-2-455.

Chan, C.S., Chan, K.G., Tay, Y.L., Chua, Y. H., Goh, K.M., 2015. Diversity of thermophiles in a Malaysian hot spring determined using 16S rRNA and shotgun metagenome sequencing. Front. Microbiol. 6, 177. https://doi.org/10.3389/fmicb.2015.00177.

Chin, J.P., Megaw, J., Magill, C.L., Nowotarski, K., Williams, J.P., Bhaganna, P., et al., 2010. Solutes determine the temperature windows for microbial survival and growth. Proc. Natl. Acad. Sci. U.S.A. 107, 7835–7840. https://doi.org/10.1073/pnas.1000557107.

Chowdhury, A.N., Bose, B.B., Pal, J., Yudhisthir., Sengupta, N.R., 1984. Studies of some minor and rare elements in hot spring deposit from Puga, Ladakh. Geol. Surv. India. Spec. Publ. 12, 585–591.

Chowdhury, A.N., Handa, B.K., Das, A.K., 1974. High lithium, rubidium and cesium contents of thermal spring water, spring sediments and borax deposits in Puga valley, Kashmir, India. Geochem. J. 8, 61–65. https://doi.org/10.2343/geochemj.8.61.

Cline, J.D., 1969. Spectrophotometric determination of hydrogen sulfide in natural water. Anal. Chem. 14, 454–458. https://doi.org/10.4319/lo.1969.14.3.0454.

Colaco, C., Sen, S., Thangavelu, M., Pinder, S., Roser, B., 1992. Extraordinary stability of enzymes dried in trehalose: simplified molecular biology. Nature. Biotechnol. 10, 1007. https://doi.org/10.1038/nbt0992-1007.

Colman, D.R., Lindsay, M.R., Boyd, E.S., 2019. Mixing of meteoric and geothermal fluids support hyperdiverse chemosynthetic hydrothermal communities. Nat. Comm. 10, 681. https://doi.org/10.1038/s41467-019-08499-1.

Cray, J.A., Russell, J.T., Timson, D.J., Singhal, R.S., Hallsworth, J.E., 2013. A universal measure of chaotropicity and kosmotropicity. Environ. Microbiol. 15, 287–296. https://doi.org/10.1111/1462-2920.12018.

Cray, J.A., Stevenson, A., Ball, P., Bankar, S.B., Eleutherio, E.C., Ezeji, T.C., et al., 2015. Chaotropicity: a key factor in product tolerance of biofuel-producing microorganisms. Curr. Opin. Biotechnol. 33, 228–259. https://doi.org/10.1016/j.copbio.2015.02.010.

Cowan, D.A., Ramond, J.B., Makhalanyane, T.P., De Maayer, P., 2015. Metagenomics of extreme environments. Curr. Opin. Microbiol. 25, 97–102.

de Lima Alves, F., Stevenson, A., Baxter, E., Gillion, J.L., Hejazi, F., Hayes, S., et al., 2015. Concomitant osmotic and chaotropicity-induced stresses in *Aspergillus wentii*: compatible solutes determine the biotic window. Curr. Genet. 61, 457–477. https://doi.org/10.1007/s00294-015-0496-8.

Dekker, N.H., Viard, T., de La Tour, C.B., Duguet, M., Bensimon, D., Croquette, V., 2003. Thermophilic topoisomerase I on a single DNA molecule. J. Mol. Biol. 329, 271–282. https://doi.org/10.1016/S0022-2836(03)00320-6.

Dibrova, D.V., Galperin, M.Y., Mulkidjanian, A.Y., 2014. Phylogenomic reconstruction of archaeal fatty acid metabolism. Environ. Microbiol. 16, 907–918. https://doi.org/10.1111/1462-2920.12359.

Edgar, R.C., 2013. UPARSE: highly accurate OTU sequences from microbial amplicon reads. Nat. Methods. 10, 996–998. https://doi.org/10.1038/nmeth.2604.

Endo, A., Sasaki, M., Maruyama, A., Kurusu, Y., 2006. Temperature adaptation of *Bacillus subtilis* by chromosomal groEL replacement. Biosci. Biotechnol. Biochem. 70, 2357–2362. https://doi.org/10.1271/bbb.50689.

Ezemaduka, A.N., Yu, J., Shi, X., Zhang, K., Yin, C.C., Fu, X., et al., 2014. A small heat shock protein enables *Escherichia coli* to grow at a lethal temperature of 50°C conceivably by maintaining cell envelope integrity. J. Bacteriol. 196, 2004–2011. https://doi.org/10.1128/JB.01473-14.

Fernandes, S., Mazumdar, A., Bhattacharya, S., Peketi, A., Mapder, T., Roy, R., et al., 2018. Enhanced carbon-sulfur cycling in the sediments of Arabian Sea oxygen minimum zone center. Sci. Rep. 8, 8665. https://doi.org/10.1038/s41598-018-27002-2.

Few, S., 2009. Now you see it: simple visualization techniques for quantitative analysis. USA: Analytics Press.

Fields, P. A., 2001. Protein function at thermal extremes: balancing stability and flexibility. Comp. Biochem. Physiol. A. Mol. Lntegr. Physiol. 129, 417–431. https://doi.org/10.1016/S1095-6433(00)00359-7.

Fields, S., Johnston, M., 2005. Whither model organism research? Science. 307, 1885–1886. https://doi.org/10.1126/science.1108872.

Galinski, E.A., 1995. Osmoadaptation in bacteria. Adv. Microb. Physiol. 37, 273–328. https://doi.org/10.1016/S0065-2911(08)60148-4.

Galinski, E.A., Troper, H.G., 1982. Betaine, a compatible solute in the extremely halophilic phototrophic bacterium *Ectothiorhodospira halocloris*. FEMS. Microbiol. Lett. 13, 357–360. https://doi.org/10.1111/j.1574-6968.1982.tb08287.x.

Galinski, E.A., Tròper, H.G., 1994. Microbial behaviour in salt stressed ecosystems. FEMS. Microbiol. Rev. 15, 95–108. https://doi.org/10.1111/j.1574-6976.1994.tb00128.x.

Gansser, A., 1964. Geology of the Himalayas. London: Interscience Publishers.

Garrett, D.E., 1998. Borates: handbook of deposits, processing, properties, and use. New York: Academic Press.

Gerhardt, P., Murray, R.G.E., Wood, W.A., Krieg, N.R., 1994. Methods for General and Molecular Bacteriology. Washington: American Society for Microbiology.

Ghosh, W., Roy, P., 2006. *Mesorhizobium thiogangeticum* sp. nov., a novel sulfur-oxidizing chemolithoautotroph from rhizosphere soil of an Indian tropical leguminous plant. Int. J. Syst. Evol. Microbiol. 56, 91–97. https://doi.org/10.1099/ijs.0.63967-0.

Ghosh, W., Mallick, S., Haldar, P.K., Pal, B., Maikap, S.C., Gupta, S.K.D., 2012. Molecular and Cellular Fossils of a Mat-Like Microbial Community in Geothermal Boratic Sinters. Geomicrobiol. J. 29, 879–885. https://doi.org/10.1080/01490451.2011.635761.

Ghosh, W., Roy, C., Roy, R., Nilawe, P., Mukherjee, A., Haldar, P.K., et al., 2015. Resilience and receptivity worked in tandem to sustain a geothermal mat community amidst erratic environmental conditions. Sci. Rep. 5, 12179. https://doi.org/10.1038/srep12179(2015).

Goh, K.M., Gan, H.M., Chan, K.G., Chan, G.F., Shahar, S., Chong, C.S., et al., 2014. Analysis of *Anoxybacillus* genomes from the aspects of lifestyle adaptations, prophage diversity, and carbohydrate metabolism. PLoS. One. 9, e90549. https://doi.org/10.1371/journal.pone.0090549.

Grasshoff, K., Kremling, K., Ehrhardt, M., 1999. Methods of seawater analysis. Weinheim: Wiley-VCH.

Hallsworth, J.E., Prior, B.A., Nomura, Y., Iwahara, M., Timmis, K.N., 2003. Compatible solutes protect against chaotrope (ethanol)-induced, nonosmotic water stress. Appl. Environ. Microbiol. 69, 7032–7034. https://doi.org/10.1128/AEM.69.12.7032-7034.2003.

Hallsworth, J.E., Yakimov, M.M., Golyshin, P.N., Gillion, J.L., D’Auria, G., de Lima Alves, F., et al., 2007. Limits of life in MgCl_2_-containing environments: chaotropicity defines the window. Environ. Microbiol. 9, 801–813. https://doi.org/10.1111/j.1462-2920.2006.01212.x.

Hamerly, T., Tripet, B., Wurch, L., Hettich, R.L., Podar, M., Bothner, B., et al., 2015. Characterization of Fatty Acids in Crenarchaeota by GC-MS and NMR. Archaea. 2015. https://doi.org/10.1155/2015/472726.

Harinarayana, T., Azeez, K.K.A., Murthy, D.N., Veeraswamy, K., Rao, S.P.E., Manoj, C., et al., 2006. Exploration of geothermal structure in Puga geothermal field, Ladakh Himalayas, India by magnetotelluric studies. J. Appl. Geophys. 58, 280–295. https://doi.org/10.1016/j.jappgeo.2005.05.005.

Hedlund, B.P., Murugapiran, S.K., Alba, T.W., Levy, A., Dodsworth, J.A., Goertz, G.B., et al., 2015. Uncultivated thermophiles: current status and spotlight on ‘Aigarchaeota’. Curr. Opin. Microbiol. 25, 136–145. https://doi.org/10.1016/j.mib.2015.06.008.

Hesketh, J.E., Loudon, J.B., Reading, H.W., Glen, A.I., 1978. The effect of lithium treatment on erythrocyte membrane ATPase activities and erythrocyte ion content. Br. J. Clin. Pharmacol. 5, 323–329. https://doi.org/10.1111/j.1365-2125.1978.tb01715.x.

Hetzer, A., Morgan, H.W., McDonald, I.R., Daughney, C.J., 2007. Microbial life in Champagne Pool, a geothermal spring in Waiotapu, New Zealand. Extremophiles. 11, 605–614. https://doi.org/10.1007/s00792-007-0073-2.

Hribar, B., Southall, N.T., Vlachy, V., Dill, K.A., 2002. How ions affect the structure of water. J. Am. Chem. Soc. 124, 12302–12311. https://doi.org/10.1021/ja026014h.

Huang, Q., Dong, C.Z., Dong, R.M., Jiang, H., Wang, S., Wang, G., et al., 2011. Archaeal and bacterial diversity in hot springs on the Tibetan Plateau, China. Extremophiles. 15, 549. https://doi.org/10.1007/s00792-011-0386-z.

Huang, Q., Jiang, H., Briggs, B.R., Wang, S., Hou, W., Li, G., et al., 2013. Archaeal and bacterial diversity in acidic to circumneutral hot springs in the Philippines. FEMS. Mcrobiol. Ecol. 85, 452–464. https://doi.org/10.1111/1574-6941.12134.

Hussein, E.I., Jacob, J.H., Shakhatreh, M.A.K., Abd Alrazaq, M.A., Juhmani, A.S.F., Cornelison, C.T., 2017. Exploring the microbial diversity in Jordanian hot springs by comparative metagenomic analysis. Microbiology Open. 6, e00521. https://doi.org/10.1002/mbo3.521.

Jiménez, D.J., Andreote, F.D., Chaves, D., Montaña, J.S., Osorio-Forero, C., Junca, H., et al., 2012. Structural and functional insights from the metagenome of an acidic hot spring microbial planktonic community in the Colombian Andes. PloS. One. 7, e52069. https://doi.org/10.1371/journal.pone.0052069.

Jing, X., Evangelista Falcon, W., Baudry, J., Serpersu, E.H., 2017. Thermophilic Enzyme or Mesophilic Enzyme with Enhanced Thermostability: Can We Draw a Line?. J. Phys. Chem. B. 121, 7086–7094. https://doi.org/10.1021/acs.jpcb.7b04519.

Johnson, D.B., Body, D.A., Bridge, T.A.M., Bruhn, D.F., Roberto, F.F., 2001. Biodiversity of Acidophilic Moderate Thermophiles Isolated from Two Sites in Yellowstone National Park and Their Roles in the Dissimilatory Oxido-Reduction of Iron, ed. A.L. Reysenbach, M. Voytek, and R. Mancinelli (Boston: Springer US) 23–39.

Johnson, D.B., Okibe, N., Roberto, F.F., 2003. Novel thermo-acidophilic bacteria isolated from geothermal sites in Yellowstone National Park: physiological and phylogenetic characteristics. Arch. Microbiol. 180, 60–68. https://doi.org/10.1007/s00203-003-0562-3.

Jones, B., Renaut, R.W., Rosen, M.R., 2000. Stromatolites Forming in Acidic Hot-Spring Waters, North Island, New Zealand. Palaios. 15, 450–475. https://doi.org/10.1669/0883-1351(2000)015<0450:SFIAHS>2.0.CO;2.

Kelleher, C., Wagener, T., 2011. Ten guidelines for effective data visualization in scientific publications. Environ. Modell. Softw. 26, 822–827. https://doi.org/10.1016/j.envsoft.2010.12.006.

Kelly, D.P., Wood, A.P., 1994. Synthesis and determination of thiosulfate and polythionates. Methods. Enzymol. 243, 475–501. https://doi.org/10.1016/0076-6879(94)43037-3.

Koga, Y., 2012. Thermal adaptation of the archaeal and bacterial lipid membranes. Archaea. 2012. https://doi.org/10.1155/2012/789652.

Koynova, R., Brankov, J., Tenchov, B., 1997. Modulation of lipid phase behavior by kosmotropic and chaotropic solutes: Experiment and thermodynamic theory. Eur. Biophys. J. 25, 261–274. https://doi.org/10.1007/s002490050038.

Kristjansson, J.K., Stetter, K.O., 1992. Thermophilic bacteria, ed. J.K. Kristjansson (London: CRC Press Inc), 1–18.

Kumar, S., Tsai, C.J., Nussinov, R., 2000. Factors enhancing protein thermostability. Protein. Eng. 13, 179–191. https://doi.org/10.1093/protein/13.3.179.

Kosara, R., Mackinlay, J., 2013. Storytelling: The next step for visualization. Computer. 46, 44–50. https://doi.org/10.1109/MC.2013.36.

Kvist, T., Ahring, B.K., Westermann, P., 2007. Archaeal diversity in Icelandic hot springs. FEMS. Microbiol. Ecol. 59, 71–80. https://doi.org/10.1111/j.1574-6941.2006.00209.x.

Lamark, T., Styrvold, O.B., Strom, A.R., 1992. Efflux of choline and glycine betaine from osmoregulating cells of *Escherichia coli*. FEMS. Microbiol. Lett. 96, 149–154. https://doi.org/10.1111/j.1574-6968.1992.tb05408.x.

Lippert, K., Galinski, E.A., 1992. Enzyme stabilization by ectoine-type compatible solutes: protection against heating, freezing and drying. Appl. Microbiol. Biotechnol. 37, 61–65. https://doi.org/10.1007/BF00174204.

López-López, O., Knapik, K., Cerdán, M.E., González-Siso, M.I. 2015. Metagenomics of an alkaline hot spring in Galicia (Spain): microbial diversity analysis and screening for novel lipolytic enzymes. Front. Microbiol. 6, 1291. doi:10.3389/fmicb.2015.01291.

Mancinelli, R., Botti, A., Bruni, F., Ricci, M.A., Soper, A.K., 2007. Hydration of sodium, potassium, and chloride ions in solution and the concept of structure maker/breaker. J. Phys. Chem. B. 111, 13570–13577. https://doi.org/10.1021/jp075913v.

Martin, W., Baross, J., Kelley, D., Russell, M.J., 2008. Hydrothermal vents and the origin of life. Nat. Rev. Microbiol. 6, 805–814. https://doi.org/10.1038/nrmicro1991.

Martinez, W.L., Martinez, A.R., Solka, J., 2017. Exploratory data analysis with MATLAB. Abingdon: CRC Press.

Mazumdar, A., Dewangan, P., Joäo, H.M., Peketi, A., Khosla, V.R., Kocherla, M., et al., 2009. Evidence of paleo&#8211;cold seep activity from the Bay of Bengal, offshore India. Geochem. Geophy. Geosy. 10, 1–15. https://doi.org/10.1029/2008GC002337.

Mendonca, A.F., Knabel, S.J., 1994. A novel strictly anaerobic recovery and enrichment system incorporating lithium for detection of heat-injured *Listeria monocytogenes* in pasteurized milk containing background microflora. Appl. Environ. Microbiol. 60, 4001–4008.

Menzel, P., Gudbergsdóttir, S.R., Rike, A.G., Lin, L., Zhang, Q., Contursi, P., et al., 2015. Comparative Metagenomics of Eight Geographically Remote Terrestrial Hot Springs. Microb. Ecol. 70, 411–424. https://doi.org/10.1007/s00248-015-0576-9.

Moreira, C., Rainey, F.A., Nobre, M.F., da Silva, M.T., da Costa, M.S., 2000. *Tepidimonas ignava* gen. nov., sp. nov., a new chemolithoheterotrophic and slightly thermophilic member of the beta-Proteobacteria. Int. J. Syst. Evol. Microbiol. 50, 735–742. https://doi.org/10.1099/00207713-50-2-735.

Navada, S.V., Rao, S.M., 1991. Isotope Studies of some Geothermal Waters in India. Isot. Environ. Healt. S. 27, 153–163. https://doi.org/10.1080/10256019108622498.

Owen, R.B., Renaut, R.W., Jones, B., 2008. Geothermal diatoms: a comparative study of floras in hot spring systems of Iceland, New Zealand, and Kenya. Hydrobiologia. 610, 175–192. https://doi.org/10.1007/s10750-008-9432-y.

Pace, N.R., 1997. A molecular view of microbial diversity and the biosphere. Science. 276, 734–740. https://doi.org/10.1126/science.276.5313.734.

Parte, A.C., 2013. LPSN-list of prokaryotic names with standing in nomenclature. Nucleic Acids. Res. 42, 613–616. https://doi.org/10.1093/nar/gkt1111.

Pitzurra, M., Szybalski, W., 1959. Formation and multiplication of spheroplasts of *Escherichia coli* in the presence of lithium chloride. J. Bacteriol. 77, 614.

Power, J.F., Carere, C.R., Lee, C.K., Wakerley, G.L., Evans, D.W., Button, M., et al., 2018. Microbial biogeography of 925 geothermal springs in New Zealand. Nat. Commun. 9, 2876. https://doi.org/10.1038/s41467-018-05020-y.

Pucci, F., Rooman, M., 2017. Physical and molecular bases of protein thermal stability and cold adaptation. Curr. Opin. Struct. Biol. 42, 117–128. https://doi.org/10.1016/j.sbi.2016.12.007.

Qu, Y., Bolen, C.L., Bolen, D.W., 1998. Osmolyte-driven contraction of a random coil protein. Proc. Natl. Acad. Sci. USA. 23, 143–148. https://doi.org/10.1073/pnas.95.16.9268.

Rainey, F.A., Silva, J., Nobre, M.F., Silva, M.T., da Costa, M.S., 2003. *Porphyrobacter cryptus* sp. nov., a novel slightly thermophilic, aerobic, bacteriochlorophyll a-containing species. Int. J. Syst. Evol. Microbiol. 53, 35–41. https://doi.org/10.1099/ijs.0.02308-0.

Rantz, S. E., 1982. Measurement of discharge by tracer dilution. In: Measurement and computation of streamflow: volume 1. Measurement of stage and discharge, (Washington DC, USA: Geological Survey and U.S. Government Printing Office), 211–259.

Record, Jr M.T., Courtenay, E.S., Cayley, D.S., Guttman, H.J., 1998. Responses of *Escherichia coli* to osmotic stress: large changes in amounts of cytoplasmic solutes and water. Trends. Biochem. Sci. 23, 143–148. https://doi.org/10.1016/S0968-0004(98)01196-7.

Ribarsky, W., Ayers, E., Eble, J., Mukherjea, S., 1994. Glyphmaker: creating customized visualizations of complex data. Computer. 27, 57–64. https://doi.org/10.1109/2.299412.

Roark, T.C., Palacio, L.A., Gurnev, P.A., Ray, B.D., Petrache, H.I., 2012. Interactions of Lithium Ions with Lipid Membranes. Biophys. J. 102, 96. https://doi.org/10.1016/j.bpj.2011.11.545.

Roeselers, G., Norris, T.B., Castenholz, R.W., Rysgaard, S., Glud, R.N., Kuhl, M., et al., 2007. Diversity of phototrophic bacteria in microbial mats from Arctic hot springs (Greenland). Environ. Microbiol. 9, 26–38. https://doi.org/10.1111/j.1462-2920.2006.01103.x.

Rosenstein, R., Futter-Bryniok, D., Gotz, F., 1999. The Choline-Converting Pathway in *Staphylococcus xylosus* C2A: Genetic and Physiological Characterization. J. Bacteriol. 181, 2273–2278.

Roy, C., Alam, M., Mandal, S., Haldar, P.K., Bhattacharya, S., Mukherjee, T., et al., 2016. Global Association between Thermophilicity and Vancomycin Susceptibility in Bacteria. Front. Microbiol. 7, 412. https://doi.org/10.3389/fmicb.2016.00412.

Rudolph, B., Gebendorfer, K.M., Buchner, J., Winter, J., 2010. Evolution of *Escherichia coli* for growth at high temperatures. J. Biol. Chem. 285, 19029–19034. https://doi.org/10.1074/jbc.M110.103374.

Samaddar, S., Grewal, R.K., Sinha, S., Ghosh, S., Roy, S., Gupta, S.K., 2016. Dynamics of Mycobacteriophage-Mycobacterial host interaction&#8211;evidence for secondary mechanisms for host lethality. Appl. Environ. Microbiol. 82, 124–133. https://doi.org/10.1128/AEM.02700-15.

Saxena, V.K., D’Amore, F., 1984. Aquifer chemistry of the Puga and Chumathang high temperature geothermal systems in India. J. Volcanol. Geotherm. 21, 333–346. https://doi.org/10.1016/0377-0273(84)90029-5.

Schlegel, H.G., Zaborosch, C., 1993. General microbiology, (New York: Cambridge University Press) 212.

Schwartzman, D.W., Lineweaver, C.H., 2004. The hyperthermophilic origin of life revisited. Biochem. Soc. Trans. 32, 168–171. https://doi.org/10.1042/bst0320168.

Shanker, R., 1988. Heat flow map of India and its geological and economic significance. Indian Minerals. 42, 89–110.

Shanker, R., Absar, A., Bajpai, P., 2012. Heat flow map of India update. 21st New Zealand Geothermal Workshop. 157–162.

Shih, T.W., Pan, T.M., 2011. Stress responses of thermophilic *Geobacillus* sp. NTU 03 caused by heat and heat-induced stress. Microbiol. Res. 166, 346–359. https://doi.org/10.1016/j.micres.2010.08.001.

Skirnisdottir, S., Hreggvidsson, G.O., Hjorleifsdottir, S., Marteinsson, V.T., Petursdottir, S.K., Holst, O., et al., 2000. Influence of sulfide and temperature on species composition and community structure of hot spring microbial mats. Appl. Environ. Microbiol. 66, 2835–2841. https://doi.org/10.1128/AEM.66.7.2835-2841.2000.

Sleator, R.D., Hill, C., 2002. Bacterial osmoadaptation: the role of osmolytes in bacterial stress and virulence. FEMS. Microbiol. Rev. 26, 49–71. https://doi.org/10.1111/j.1574-6976.2002.tb00598.x.

Stetter, K.O., 1999. Extremophiles and their adaptation to hot environments. FEBS. Lett. 452, 22–25. https://doi.org/10.1016/S0014-5793(99)00663-8.

Strøm, A. R., Kaasen, I., 1993. Trehalose metabolism in *Escherichia coli*: stress protection and stress regulation of gene expression. Mol. Microbiol. 8, 205–210. https://doi.org/10.1111/j.1365-2958.1993.tb01564.x.

Team, R.C., 2014. R: A Language and Environment for Statistical Computing. Vienna: R Core Team. R. Found. Stat. Comput. 1, 409. https://doi.org/http://www.R-project.org/.

Vieille, C., Zeikus, G.J., 2001. Hyperthermophilic enzymes: sources, uses, and molecular mechanisms for thermostability. Microbiol. Mol. Biol. Rev. 65, 1–43. https://doi.org/10.1128/MMBR.65.1.1-43.2001.

Wang, W., Ma, T., Zhang, B., Yao, N., Li, M., Cui, L., et al., 2014. A novel mechanism of protein thermostability: a unique N-terminal domain confers heat resistance to Fe/Mn-SODs. Sci. Rep. 4, 7284. https://doi.org/10.1038/srep07284.

Ware, C., 2012. Information visualization: perception for design. Waltham: Morgan Kaufman.

Welsh, D.T., 2000. Ecological significance of compatible solute accumulation by microorganisms: from single cells to global climate. FEMS. Microbiol. Rev. 24, 263–290. https://doi.org/10.1111/j.1574-6976.2000.tb00542.x.

Wemheuer, B., Taube, R., Akyol, P., Wemheuer, F., Daniel, R., 2013. Microbial diversity and biochemical potential encoded by thermal spring metagenomes derived from the Kamchatka Peninsula. Archaea. 2013. https://doi.org/10.1155/2013/136714.

West, P.W, Gaeke, G.C., 1956. Fixation of sulfur dioxide as disulfitomercurate (II) and subsequent colorimetric estimation. Anal. Chem. 28, 1816–1819. https://doi.org/10.1021/ac60120a005.

Wieder, R., 1999. Cell Growth, Differentiation and Senescence: A Practical Approach, ed. G. Studzinski (Oxford: Oxford University Press), 52.

Wiggins, P.M., 2001. High and low density intracellular water. Cell. Mol. Biol. 47, 735–744.

Xie, W., Wang, F., Guo, L., Chen, Z., Sievert, S.M., Meng, J., et al., 2011. Comparative metagenomics of microbial communities inhabiting deep-sea hydrothermal vent chimneys with contrasting chemistries. ISME. J. 5, 414–426. https://doi.org/10.1038/ismej.2010.144.

Xu, Y., Schoonen, M.A.A., Nordstrom, D.K., Cunningham, K.M., Ball, J.W., 1998. Sulfur geochemistry of hydrothermal waters in Yellowstone National Park: I The origin of thiosulfate in hot spring waters. Geochim. Cosmochim. Acta. 62, 3729–3743. https://doi.org/10.1016/S0016-7037(98)00269-5.

Yakimov, M.M., La Cono, V., Spada, G.L., Bortoluzzi, G., Messina, E., Smedile, F., et al., 2015. Microbial community of the deep-sea brine Lake Kryos seawater-brine interface is active below the chaotropicity limit of life as revealed by recovery of mRNA. Environ. Microbiol. 17, 364–382. https://doi.org/10.1111/1462-2920.12587.

Zamarreño, A., Cantera, R.G., Garcia-Mina, J.M., 1997. Extraction and determination of glycine betaine in liquid fertilizers. J. Agric. Food. Chem. 45, 774–776. https://doi.org/10.1021/jf960342h.

Zeldovich, K.B., Berezovsky, I.N., Shakhnovich, E.I., 2007. Protein and DNA sequence determinants of thermophilic adaptation. PLoS. Comput. Biol. 3, e5. https://doi.org/10.1371/journal.pcbi.0030005.

Ziegler, M., Seneca, F.O., Yum, L.K., Palumbi, S.R., Voolstra, C.R., 2017. Bacterial community dynamics are linked to patterns of coral heat tolerance. Nat. Commun. 8, 14213. https://doi.org/10.1038/ncomms14213.

Zhou, N., Ishchuk, O.P., Knecht, W., Compagno, C., Piskur, J., 2019. Improvement of thermotolerance in *Lachancea thermotolerans* using a bacterial selection pressure. J. Ind. Microbiol. Biotechnol. 46, 133–145. https://doi.org/10.1007/s10295-018-2107-4.

