## Supplementary Tables, Supplementary Notes and Supplementary figures for "Inorganic salts and compatible solutes help mesophilic bacteria inhabit the high temperature waters of a Trans-Himalayan sulfur-borax spring"

**Present Address:**

^#^	**Equal contributions**

***	Correspondence:	Wriddhiman Ghosh**

**Table S1.** Number of species-level microbial entities (OTUs)^1^ reported from geographically-discrete terrestrial hot springs.

| **Sl. No.** | **Name and Geographical Location of the vent** | **Temperature of the sample-site (in °C)** | **pH of the sample-site** | **No. of OTUs^1^ identified** | **Reference** |
| --- | --- | --- | --- | --- | --- |
| 1 | Diretiyanqu, China^2^ | 85.1 | 2.58 | 49 | (Hou *et al*., 2013) |
| 2 | Jiemeiquan -Sisters Spring, China^2^ | 93.6 | 9.25 | 59 | (Hou *et al*., 2013) |
| 3 | Zhenzhuquan – Pearl Spring, China^2^ | 89.1 | 4.79 | 46 | (Hou *et al*., 2013) |
| 4 | Huitaijing – Pregnancy Spring, China^2^ | 92.3 | 8.05 | 63 | (Hou *et al*., 2013) |
| 5 | Shuirebaozha - Hydrothermal Explosion, China^2^ | 79.8 | 8.04 | 87 | (Hou *et al*., 2013) |
| 6 | Dagunguo – Great Boiling Pot, China^2^ | 84.5 | 7.20 | 21 | (Hou *et al*., 2013) |
| 7 | Gumingquan– Drum Beating Spring, China^2^ | 93.0 | 9.35 | 82 | (Hou *et al*., 2013) |
| 8 | Gongxiaoshe – Co-op Hotel (side), China^2^ | 73.8 | 7.29 | 41 | (Hou *et al*., 2013) |
| 9 | Jinze - Golden Pond Motel, China^2^ | 81.6 | 6.71 | 40 | (Hou *et al*., 2013) |
| 10 | Jermuk, Southeast Armenia^2,4^ | 53.0 | 7.50 | 209 | (Hedlund *et al*., 2013) |
| 11 | Arzakan, Southeast Armenia^2,4^ | 44.0 | 7.20 | 321 | (Hedlund *et al*., 2013) |
| 12 | Lake Magadi, Hot spring 1, Kenya^3,4^ | 45.1 | 9.80 | 377 | (Kambura *et al*., 2016) |
| 13 | Lake Magadi, Hot spring 2, Kenya^3,4^ | 83.6 | 9.40 | 680 | (Kambura *et al*., 2016) |
| 14 | Lake Magadi, Hot spring 3, Kenya^3,4^ | 81.0 | 9.20 | 568 | (Kambura *et al*., 2016) |
| 15 | Mqhephu hot spring, Limpopo Province of South Africa^3^ | 43.0 | 8.19 | 224 | (Tekere *et al*., 2015) |
| 16 | Tshipise hot spring, Limpopo Province of South Africa^3^ | 58.0 | 8.94 | 101 | (Tekere *et al*., 2015) |
| 17 | Sagole hot spring, Limpopo Province of South Africa^3^ | 45.0 | 9.24 | 303 | (Tekere *et al*., 2015) |
| 18 | Siloam hot spring, Limpopo Province of South Africa^3^ | 63.0 | 9.50 | 251 | (Tekere *et al*., 2015) |
| 19 | Soutinig hot spring, Limpopo Province of South Africa^3^ | 43.9 | 7.20 | 105 | (Tekere *et al*., 2015) |
| 20 | Eiland hot spring, Limpopo Province of South Africa^3^ | 41.0 | 7.63 | 6 | (Tekere *et al*., 2015) |
| 21 | Great Boiling Spring 1002W, Great Basin, USA^3^ | 82.0 | 6.63 | 36 | (Cole *et al*., 2012) |
| 22 | Akwar, Eritrea^3,4^ | 49.5 | 7.22 | 1898 | (Ghilamicael *et al*., 2017) |
| 23 | Elegedi, Eritrea^3,4^ | 100.0 | 7.01 | 1662 | (Ghilamicael *et al*., 2017) |
| 24 | Garbanabra, Eritrea^3,4^ | 51.3 | 7.05 | 2354 | (Ghilamicael *et al*., 2017) |
| 25 | Gelti, Eritrea^3,4^ | 52.6 | 7.19 | 1834 | (Ghilamicael *et al*., 2017) |
| 26 | Maiwooi, Eritrea^3,4^ | 51.4 | 7.54 | 1054 | (Ghilamicael *et al*., 2017) |
| 27 | Icelandic hot springs P0, Iceland^3^ | 100.0 | 3.80 | 84 | (Krebs *et al*., 2014) |
| 28 | Icelandic hot springs P1, Iceland^3^ | 59.0 | 3.20 | 529 | (Krebs *et al*., 2014) |
| 29 | Icelandic hot springs P2, Iceland^3^ | 64.0 | 2.90 | 168 | (Krebs *et al*., 2014) |
| 30 | Icelandic hot springs P3, Iceland^3^ | 57.0 | 2.90 | 674 | (Krebs *et al*., 2014) |
| 31 | Leirgerdur, Iceland^3^ | 98.0 | 2.00 | 19 | (Krebs *et al*., 2014) |
| 32 | Hrifla spring, Iceland^3^ | 98.0 | 7.00 | 19 | (Krebs *et al*., 2014) |
| 33 | Lotus Pond HTP^3^, as explored at 17:30 h of 23.07.2013 | 78.0 | 8 | 331 | (Roy *et al*., 2016) |
| 34 | Lotus Pond HTP^3^, as explored at 8:30 h of 20.10.2014 | 84.0 | 7.5 | 109 | (This study) |
| 35 | Lotus Pond HTP^3^, as explored at 14:30 h of 20.10.2014 | 81.0 | 7.8 | 279 | (This study) |
| 36 | Lotus Pond MTP^3^, as explored at 8:30 h of 20.10.2014 | 74.0 | 7.5 | 635 | (This study) |
| 37 | Lotus Pond MTP^3^, as explored at 14:30 h of 20.10.2014 | 72.0 | 7.5 | 1205 | (This study) |

^1^	In all the reports considered, microbial diversity of aquatic or semi-aquatic communities was estimated in terms of the total number of species-level entities or operational taxonomic units (OTUs) present. OTUs were determined based on high throughput 16S rRNA genes sequence data.

^2^	OTU counts for these hydrothermal communities involved both bacteria and archaea.

^3^	OTU counts for these hydrothermal communities involved only bacteria.

^4^	These studies involved semi-aquatic communities; all the other studies involved aquatic communities.

**References used in Table S1**

Cole JK, Peacock JP, Dodsworth JA, Williams AJ, Thompson DB, Dong H, et al. Sediment microbial communities in Great Boiling Spring are controlled by temperature and distinct from water communities. ISME J. 2013;7:718.

Ghilamicael AM, Budambula NL, Anami SE, Mehari T, Boga HI. Evaluation of prokaryotic diversity of five hot springs in Eritrea. BMC Microbiol. 2017;17:203.

Hedlund BP, Dodsworth JA, Cole JK, Panosyan HH. An integrated study reveals diverse methanogens, Thaumarchaeota, and yet-uncultivated archaeal lineages in Armenian hot springs. Antonie van Leeuwenhoek. 2013;104:71-82.

Hou W, Wang S, Dong H, Jiang H, Briggs BR, Peacock JP, et al. A comprehensive census of microbial diversity in hot springs of Tengchong, Yunnan Province China using 16S rRNA gene pyrosequencing. PloS One. 2013;8:53350.

Kambura AK, Mwirichia RK, Kasili RW, Karanja EN, Makonde HM, Boga HI. Bacteria and Archaea diversity within the hot springs of Lake Magadi and Little Magadi in Kenya. BMC Microbiol. 2016;16:136.

Krebs JE, Vaishampayan P, Probst AJ, Tom LM, Marteinsson VT, Andersen GL, et al. Microbial community structures of novel Icelandic hot spring systems revealed by PhyloChip G3 analysis. Astrobiology. 2014;14:229-240.

Roy C, Alam M, Mandal S, Haldar PK, Bhattacharya S, Mukherjee T, et al. Global association between thermophilicity and vancomycin susceptibility in Bacteria. Front Microbiol. 2016;7:412.

Tekere M, Lötter A, Olivier J, Venter S. Bacterial diversity in some South African thermal springs:a metagenomic analysis. Proc World Geotherm Congr. 2015;19-25.

**Table S2.** Major parameters of bacterial diversity and water chemistry measured for the HTP, MTP, LTP and RVW water-samples at 8:30 and 14:30 h of 20 October 2014.

|  |  | **8:30 h** | | | | **14:30 h** | | | |
| --- | --- | --- | --- | --- | --- | --- | --- | --- | --- |
| **Serial**  **no.** | **Physicochemical /**  **microbial parameters** | **HTP** | **MTP** | **LTP** | **RVW** | **HTP** | **MTP** | **LTP** | **RVW** |
| **1** | **Temperature (in °C)** | 84 | 74 | 30-45 | 15 | 81 | 72 | 32-42 | 24 |
| **2** | **pH** | 7.5 | 7.4 | 7.2 | 6.6 | 7.8 | 7.5 | 6.8 | 6.2 |
| **3** | **No. of OTUs detected** | 109 | 635 | 1013 | 1372 | 279 | 1205 | 1691 | 2044 |
| **4** | **No. of genera detected** | 52 | 160 | 177 | 195 | 100 | 211 | 216 | 219 |
| **5** | **No. of phyla/classes detected** | 15 | 34 | 33 | 32 | 20 | 30 | 31 | 34 |
| **6** | **B conc. in water (in ppm)** | 123.9 | 145.5 | 104.4 | 62.6 | 126.7 | 128.8 | 110.3 | 66.6 |
| **7** | **Si conc. in water (in ppm)** | 65.2 | 67 | 53.9 | 36.7 | 66.7 | 67 | 59.3 | 36 |
| **8** | **Li conc. in water (in ppm)** | 4.7 | 5 | 3.9 | 2 | 4.7 | 4.6 | 3.6 | 2.1 |
| **9** | **Na conc. in water (in ppm)** | 576 | 521.6 | 451.3 | 258 | 553.5 | 546.8 | 531.3 | 266.8 |
| **10** | **Cl conc. in water (in ppm)** | 358.8 | 363.5 | 287.7 | 176 | 370.6 | 373 | 320 | 161 |
| **11** | **Ca conc. in water (in ppm)** | 2.6 | 2.6 | 2.4 | 2.4 | 2.1 | 2.5 | 2.2 | 2.2 |
| **12** | **K conc. in water (in ppm)** | 76.4 | 89.7 | 62.8 | 39.8 | 75 | 77.9 | 69.5 | 40 |
| **13** | **Mg conc. in water (in ppm)** | 2.5 | 2.9 | 2.9 | 2.9 | 2.5 | 2.5 | 2.7 | 2.3 |
| **14** | **Rb conc. in water (in ppm)** | 1.2 | 1.4 | 1 | 0.6 | 1.2 | 1.2 | 1.1 | 0.6 |
| **15** | **Sr conc. in water (in ppm)** | 0.4 | 0.4 | 0.3 | 0.2 | 0.4 | 0.4 | 0.3 | 0.2 |
| **16** | **Cs conc. in water (in ppm)** | 9.3 | 9.2 | 5.9 | 3.3 | 7.5 | 7.7 | 6.5 | 3.5 |
| **17** | **S^2-^ conc. in water (in ppm)** | 8.4 | 3.8 | 0 | 0 | 5.7 | 3 | 0 | 0 |
| **18** | **S_2_O_3_^2-^ conc. in water (in ppm)** | 145.8 | 157 | 168.2 | 89.7 | 89.7 | 100.9 | 112.1 | 56.1 |
| **19** | **SO_3_^2-^ conc. in water (in ppm)** | 21.9 | 11.2 | 0 | 0 | 8.8 | 1.2 | 0 | 0 |
| **20** | **SO_4_^2-^ conc. in water (in ppm)** | 68.4 | 88.4 | 110.7 | 66.1 | 92.5 | 107.1 | 123.1 | 69.4 |

**Table S3.** Summary statistics of the metataxonomic analysis of the aquatic communities of HTP, MTP, LTP and RVW conducted at 6:30, 8:30, 14:30 and 20:30 h of 20 October 2014.

| **Sample Identity** | **SRA BioSample Accession number** | **SRA Run Accession number** | **Total reads** | **Reads after processing** | **Reads after de-replication** | **Total OTUs after removing singletons** | **Number of Singletons** | **ACE estimation** | **Shannon index** | **Simpson index** |
| --- | --- | --- | --- | --- | --- | --- | --- | --- | --- | --- |
|  | **Samples collected at 6:30 h** | | | | | | | | | |
| **HTP** | SAMN06833865 | SRR5482141 | 37489 | 18756 | 11323 | 96 | 2131 | 96 | 2.8610 | 0.1170 |
| **MTP** | SAMN06833866 | SRR5482154 | 100017 | 53241 | 28734 | 584 | 10929 | 584 | 3.7551 | 0.0855 |
| **LTP** | SAMN06833867 | SRR5482155 | 125456 | 62154 | 35433 | 735 | 9563 | 735 | 4.2091 | 0.0637 |
| **RVW** | SAMN06833868 | SRR5482156 | 119111 | 55950 | 29371 | 933 | 11323 | 933 | 4.6294 | 0.0450 |
|  | **Samples collected at 8:30 h** | | | | | | | | | |
| **HTP** | SAMN06833873 | SRR5482162 | 32864 | 16009 | 9363 | 109 | 910 | 109 | 2.6510 | 0.1665 |
| **MTP** | SAMN06833874 | SRR5482163 | 128000 | 90881 | 43950 | 635 | 4287 | 635 | 3.3404 | 0.1274 |
| **LTP** | SAMN06833875 | SRR5482164 | 217681 | 144153 | 68249 | 1013 | 6986 | 1013 | 3.7149 | 0.1113 |
| **RVW** | SAMN06833876 | SRR5482165 | 228024 | 157463 | 84109 | 1372 | 7396 | 1372 | 4.4025 | 0.0705 |
|  | **Samples collected at 14:30 h** | | | | | | | | | |
| **HTP** | SAMN06833877 | SRR5482166 | 156408 | 72176 | 42278 | 279 | 4619 | 279 | 3.6146 | 0.0742 |
| **MTP** | SAMN06833878 | SRR5482167 | 128062 | 92155 | 53011 | 1205 | 4651 | 1205 | 4.9466 | 0.0389 |
| **LTP** | SAMN06833879 | SRR5482168 | 215123 | 123390 | 60193 | 1691 | 5434 | 1691 | 4.2222 | 0.1007 |
| **RVW** | SAMN06833880 | SRR5482169 | 228477 | 168605 | 87944 | 2044 | 8413 | 2044 | 4.9473 | 0.0486 |
|  | **Samples collected at 20:30 h** | | | | | | | | | |
| **HTP** | SAMN06833869 | SRR5482158 | 93130 | 46578 | 25454 | 237 | 4497 | 237 | 3.7464 | 0.0642 |
| **MTP** | SAMN06833870 | SRR5482159 | 115632 | 55190 | 30872 | 1158 | 11760 | 1158 | 5.3514 | 0.0202 |
| **LTP** | SAMN06833871 | SRR5482160 | 187074 | 98723 | 51289 | 1407 | 15445 | 1407 | 4.7819 | 0.0512 |
| **RVW** | SAMN06833872 | SRR5482161 | 105051 | 50626 | 26938 | 1166 | 12283 | 1166 | 5.1711 | 0.0302 |

**Table S4.** Phylum-/class-level assortment of the total number of bacterial OTUs identified at the different sample-sites of the *Lotus Pond*-*Rulang* ecosystem at 6:30, 8:30, 14:30 and 20:30 h of 20 October 2014. Phyla for which at least 5 OTUs were detected in at least one of the 16 water-samples were regarded as major phyla/classes present in the system; others were referred to as minor phyla. Since this Table is longer and wider than one page it has been provided as a separate MS Excel file (Dataset 1).

**Table S5.** Complete list of the 267 genera identified in the *Lotus Pond*-*Rulang* ecosystem at 8:30 h of 20 October 2014, together with the information about their presence (green shade) or absence (no shade) in the four sample-sites, and their capabilities, if any, to oxidize or reduce sulfur species.

| **Genus** | **HTP** | **MTP** | **LTP** | **RVW** | **Sulfur species oxidized for chemolithotrophic nutrition (Reference)** | **Sulfur species reduced for anaerobic respiration**  **(Reference)** |
| --- | --- | --- | --- | --- | --- | --- |
| *Aquabacterium* |  |  |  |  |  |  |
| *Leifsonia* |  |  |  |  |  |  |
| *Ralstonia* |  |  |  |  |  |  |
| *Allochromatium* |  |  |  |  | S^0^, HS^-^, S_2_O_3_^2-^ (Dahl *et al*., 2005) |  |
| *Anaerolinea* |  |  |  |  |  |  |
| Armatimonadetes_gp7 |  |  |  |  |  |  |
| *Caldilinea* |  |  |  |  |  |  |
| *Conexibacter* |  |  |  |  |  |  |
| *Desulfobulbus* |  |  |  |  |  | SO_4_^2-^, S_2_O_3_^2-^ (Sorokin *et al*., 2012) |
| *Silanimonas* |  |  |  |  |  |  |
| *Thermosyntropha* |  |  |  |  |  |  |
| *Alistipes* |  |  |  |  |  |  |
| *Carnobacterium* |  |  |  |  |  |  |
| *Chrysiogenes* |  |  |  |  |  |  |
| Clostridium XI |  |  |  |  |  |  |
| Clostridium XlVa |  |  |  |  |  |  |
| *Deefgea* |  |  |  |  |  |  |
| *Dethiosulfatibacter* |  |  |  |  |  | S_2_O_3_^2-^, S^0^ (Takii *et al.,* 2007) |
| *Dickeya* |  |  |  |  |  |  |
| *Giesbergeria* |  |  |  |  |  |  |
| *Gillisia* |  |  |  |  |  |  |
| Gp7 |  |  |  |  |  |  |
| *Hyphomicrobium* |  |  |  |  |  |  |
| *Megamonas* |  |  |  |  |  |  |
| *Methylocystis* |  |  |  |  |  |  |
| *Methylomonas* |  |  |  |  |  |  |
| *Nitriliruptor* |  |  |  |  |  |  |
| *Peredibacter* |  |  |  |  |  |  |
| *Planomicrobium* |  |  |  |  |  |  |
| *Polaromonas* |  |  |  |  |  |  |
| *Polynucleobacter* |  |  |  |  |  |  |
| *Prevotella* |  |  |  |  |  |  |
| *Rheinheimera* |  |  |  |  |  |  |
| *Rhizobacter* |  |  |  |  |  |  |
| *Rhodoferax* |  |  |  |  |  |  |
| *Rickettsia* |  |  |  |  |  |  |
| *Rubrivivax* |  |  |  |  |  |  |
| *Saccharofermentans* |  |  |  |  |  |  |
| *Shewanella* |  |  |  |  |  |  |
| *Sulfurospirillum* |  |  |  |  |  | S^0^, S_2_O_3_^2-^, SO_3_^2-^ (Kodama *et al.,* 2007) |
| *Sunxiuqinia* |  |  |  |  |  |  |
| *Thiocystis* |  |  |  |  | S^0^, HS^-^ (Peduzzi *et al*., 2011) |  |
| *Acetobacterium* |  |  |  |  |  |  |
| Acetothermia_genera_incertae_sedis |  |  |  |  |  |  |
| *Acidovorax* |  |  |  |  |  |  |
| *Aeromonas* |  |  |  |  |  |  |
| *Algoriphagus* |  |  |  |  |  |  |
| *Alkalibacterium* |  |  |  |  |  |  |
| Aminicenantes_genera_incertae_sedis |  |  |  |  |  |  |
| *Aquicella* |  |  |  |  |  |  |
| *Arcobacter* |  |  |  |  | HS^-^ (Sievert *et al*., 2007) |  |
| Atribacteria_genera_incertae_sedis |  |  |  |  |  |  |
| *Azonexus* |  |  |  |  |  |  |
| *Bacteriovorax* |  |  |  |  |  |  |
| *Brachymonas* |  |  |  |  |  |  |
| *Bradyrhizobium* |  |  |  |  | S_2_O_3_^2-^ (Masuda *et al*., 2010) |  |
| *Calditerrivibrio* |  |  |  |  |  |  |
| *Caloramator* |  |  |  |  |  |  |
| *Cellvibrio* |  |  |  |  |  |  |
| *Chlorobaculum* |  |  |  |  | S_2_O_3_^2-^, SO_3_^2-^, HS^-^ (Gregersen *et al*., 2011) |  |
| *Chloroflexus* |  |  |  |  |  |  |
| *Chlorophyta* |  |  |  |  |  |  |
| *Cloacibacterium* |  |  |  |  |  |  |
| Clostridium III |  |  |  |  |  |  |
| *Cryobacterium* |  |  |  |  |  |  |
| *Cryptomonadaceae* |  |  |  |  |  |  |
| *Dechloromonas* |  |  |  |  |  |  |
| *Demequina* |  |  |  |  |  |  |
| *Desulfatirhabdium* |  |  |  |  |  | S_2_O_3_^2-^, SO_4_^2-^ (Balk *et al*., 2008) |
| *Desulfomicrobium* |  |  |  |  |  | SO_4_^2-^, SO_3_^2-^, S_2_O_3_^2-^ (Rozanova *et al*., 1988) |
| *Desulfovibrio* |  |  |  |  |  | SO_3_^2-^, S_2_O_3_^2-^, S^0^ (Mogensen *et al.,* 2005) |
| *Emticicia* |  |  |  |  |  |  |
| *Ferribacterium* |  |  |  |  |  |  |
| *Fibrobacter* |  |  |  |  |  |  |
| *Flavobacterium* |  |  |  |  |  |  |
| *Geobacter* |  |  |  |  |  |  |
| *Geothrix* |  |  |  |  |  |  |
| Gp16 |  |  |  |  |  |  |
| Gp3 |  |  |  |  |  |  |
| Gp6 |  |  |  |  |  |  |
| GpI |  |  |  |  |  |  |
| GpIIa |  |  |  |  |  |  |
| *Hydrogenophaga* |  |  |  |  |  |  |
| *Hydrogenophilus* |  |  |  |  |  |  |
| *Ignavibacterium* |  |  |  |  |  |  |
| *Legionella* |  |  |  |  |  |  |
| *Limnohabitans* |  |  |  |  |  |  |
| *Meiothermus* |  |  |  |  |  |  |
| *Methylobacter* |  |  |  |  |  |  |
| *Niabella* |  |  |  |  |  |  |
| *Nocardioides* |  |  |  |  |  |  |
| *Ohtaekwangia* |  |  |  |  |  |  |
| *Paludibacter* |  |  |  |  |  |  |
| *Proteiniclasticum* |  |  |  |  |  |  |
| *Psychroflexus* |  |  |  |  |  |  |
| *Rahnella* |  |  |  |  |  |  |
| *Rhodobacter* |  |  |  |  | HS^-^ (Hansen and Imhoff, 1985) |  |
| *Roseomonas* |  |  |  |  |  |  |
| *Rubrimonas* |  |  |  |  |  |  |
| *Runella* |  |  |  |  |  |  |
| Saccharibacteria_genera_incertae_sedis |  |  |  |  |  |  |
| *Smithella* |  |  |  |  |  |  |
| Sporolactobacillaceae_incertae_sedis |  |  |  |  |  |  |
| SR1_genera_incertae_sedis |  |  |  |  |  |  |
| *Sulfuricurvum* |  |  |  |  | HS^-^, S^0^, S_2_O_3_^2-^ (Kodama and Watanabe, 2004) |  |
| *Sulfurimonas* |  |  |  |  | S^0^, HS^-^, S_2_O_3_^2-^ (Inagaki *et al*., 2003) |  |
| *Syntrophorhabdus* |  |  |  |  |  |  |
| *Thermanaerovibrio* |  |  |  |  |  |  |
| *Thermodesulfobacterium* |  |  |  |  |  | SO_4_^2-^ (Jeanthon *et al*., 2002) |
| *Thermodesulfovibrio* |  |  |  |  |  | SO_4_^2-^, S_2_O_3_^2-^, SO_3_^2-^ (Henry *et al*., 1994) |
| *Thermoflexus* |  |  |  |  |  |  |
| *Thermotoga* |  |  |  |  |  |  |
| *Thiovirga* |  |  |  |  | S_2_O_3_^2-^, S^0^, HS^-^ (Ito *et al*., 2005) |  |
| *Treponema* |  |  |  |  |  |  |
| *Acinetobacter* |  |  |  |  | S_2_O_3_^2-^, S^0^ ( Aguilar *et al*., 2008) |  |
| *Advenella* |  |  |  |  | S_2_O_3_^2-^, S_4_O_6_^2-^ (Ghosh *et al*., 2005) |  |
| *Anoxybacillus* |  |  |  |  | S_4_O_6_^2-^ (Unpublished Mandal *et al*) |  |
| *Azospirillum* |  |  |  |  | S_2_O_3_^2-^, HS^-^ (Lavrinenko *et al*., 2010) |  |
| *Bacillariophyta* |  |  |  |  |  |  |
| *Bacillus* |  |  |  |  | S_2_O_3_^2-^ (Pérez-Ibarra *et al*., 2007) |  |
| *Beijerinckia* |  |  |  |  |  |  |
| *Brevibacillus* |  |  |  |  |  |  |
| *Brevundimonas* |  |  |  |  |  |  |
| *Burkholderia* |  |  |  |  | S_2_O_3_^2-^ , S_4_O_6_^2-^, S^0^, HS^-^ (Wittke *et al*., 1997) |  |
| *Caldicellulosiruptor* |  |  |  |  |  |  |
| *Clostridium* sensu stricto |  |  |  |  |  |  |
| *Dictyoglomus* |  |  |  |  |  |  |
| *Enhydrobacter* |  |  |  |  |  |  |
| *Escherichia/Shigella* |  |  |  |  |  |  |
| *Fervidobacterium* |  |  |  |  |  |  |
| *Geobacillus* |  |  |  |  |  |  |
| GpXIII |  |  |  |  |  |  |
| *Halomonas* |  |  |  |  | S_2_O_3_^2-^ (Sorokin, 2003) |  |
| *Hydrogenobacter* |  |  |  |  | S_2_O_3_^2-^ , S^0^ (Shima and Suzuki, 1993) |  |
| *Marinobacter* |  |  |  |  |  |  |
| *Methylobacterium* |  |  |  |  | S_2_O_3_^2-^ (Anandham *et al*., 2007) |  |
| *Propionibacterium* |  |  |  |  |  |  |
| *Providencia* |  |  |  |  |  |  |
| *Pseudomonas* |  |  |  |  | S_2_O_3_^2-^ (Sorokin *et al*., 1999) |  |
| *Psychrobacter* |  |  |  |  |  |  |
| *Sediminibacterium* |  |  |  |  |  |  |
| *Staphylococcus* |  |  |  |  |  |  |
| *Streptophyta* |  |  |  |  |  |  |
| *Sulfurihydrogenibium* |  |  |  |  | S^0^, S_2_O_3_^2-^ (Nakagawa *et al*., 2005) |  |
| *Tepidimonas* |  |  |  |  |  |  |
| *Thermomonas* |  |  |  |  |  |  |
| *Thermus* |  |  |  |  | S^0^, S_2_O_3_^2-^ (Skirnisdottir *et al*., 2001) |  |
| *Thiobacillus* |  |  |  |  | S_2_O_3_^2-^, S_4_O_6_^2-^, S^0^ (Wood and Kelly, 1991) |  |
| *Thiofaba* |  |  |  |  | S_2_O_3_^2-^, S^0^, HS^-^, S_4_O_6_^2-^ (Mori and Suzuki, 2008) |  |
| *Thiothrix* |  |  |  |  | S_2_O_3_^2-^, HS^-^(Chernousova *et al*., 2009) |  |
| GpIV |  |  |  |  |  |  |
| *Acholeplasma* |  |  |  |  |  |  |
| *Bacteroides* |  |  |  |  |  |  |
| *Curvibacter* |  |  |  |  |  |  |
| *Desulfosporosinus* |  |  |  |  |  | SO_4_^2-^, S_2_O_3_^2-^, S^0^ (Sánchez-Andrea *et al*., 2015) |
| *Exiguobacterium* |  |  |  |  |  |  |
| *Fervidicella* |  |  |  |  |  |  |
| *Pleomorphomonas* |  |  |  |  |  |  |
| *Serratia* |  |  |  |  |  |  |
| *Sphingobium* |  |  |  |  |  |  |
| *Streptococcus* |  |  |  |  |  |  |
| *Sulfuricella* |  |  |  |  | S^0^, S_2_O_3_^2-^ (Kojima and Fukui, 2010) |  |
| *Brachybacterium* |  |  |  |  |  |  |
| *Dehalogenimonas* |  |  |  |  |  |  |
| *Spirochaeta* |  |  |  |  | S_2_O_3_^2-^, S^0^, HS^-^ (Dubinina *et al*., 2011) |  |
| *Actinomycetospora* |  |  |  |  |  |  |
| *Buttiauxella* |  |  |  |  |  |  |
| *Peptoniphilus* |  |  |  |  |  |  |
| *Planobacterium* |  |  |  |  |  |  |
| *Sphingomonas* |  |  |  |  |  |  |
| *Actinomyces* |  |  |  |  |  |  |
| *Alcanivorax* |  |  |  |  |  |  |
| *Alkaliphilus* |  |  |  |  |  |  |
| *Arenimonas* |  |  |  |  |  |  |
| *Arthrobacter* |  |  |  |  |  |  |
| *Blastocatella* |  |  |  |  |  |  |
| *Comamonas* |  |  |  |  |  |  |
| *Cytophaga* |  |  |  |  |  |  |
| *Deinococcus* |  |  |  |  |  |  |
| *Desulforegula* |  |  |  |  |  | SO_4_^2-^ (Rees and Patel, 2001) |
| *Duganella* |  |  |  |  |  |  |
| *Fluviicola* |  |  |  |  |  |  |
| *Friedmanniella* |  |  |  |  |  |  |
| *Gallicola* |  |  |  |  |  |  |
| *Herminiimonas* |  |  |  |  |  |  |
| *Microvirga* |  |  |  |  |  |  |
| *Ornithinimicrobium* |  |  |  |  |  |  |
| *Panacagrimonas* |  |  |  |  |  |  |
| Parcubacteria_genera_incertae_sedis |  |  |  |  |  |  |
| *Pedomicrobium* |  |  |  |  |  |  |
| *Photobacterium* |  |  |  |  |  |  |
| *Piscinibacter* |  |  |  |  |  |  |
| *Porphyromonas* |  |  |  |  |  |  |
| *Sneathiella* |  |  |  |  |  |  |
| *Sphaerotilus* |  |  |  |  |  |  |
| *Thiomonas* |  |  |  |  | S_2_O_3_^2-^ (Vésteinsdóttir *et al.,* 2011) |  |
| *Tindallia* |  |  |  |  |  |  |
| *Actinoalloteichus* |  |  |  |  |  |  |
| *Aminiphilus* |  |  |  |  |  |  |
| *Balneola* |  |  |  |  |  |  |
| *Bosea* |  |  |  |  | S_2_O_3_^2-^, S_4_O_6_^2-^ (Das *et al*., 1996) |  |
| *Cecembia* |  |  |  |  |  |  |
| *Chryseobacterium* |  |  |  |  |  |  |
| *Desulfocapsa* |  |  |  |  |  |  |
| *Elioraea* |  |  |  |  |  |  |
| *Flexibacter* |  |  |  |  |  |  |
| *Formivibrio* |  |  |  |  |  |  |
| *Gemmatimonas* |  |  |  |  |  |  |
| *Ilumatobacter* |  |  |  |  |  |  |
| *Kinneretia* |  |  |  |  |  |  |
| *Kribbella* |  |  |  |  |  |  |
| *Marmoricola* |  |  |  |  |  |  |
| *Methylosoma* |  |  |  |  |  |  |
| *Mycobacterium* |  |  |  |  |  |  |
| *Nitrospira* |  |  |  |  |  |  |
| *Planococcus* |  |  |  |  |  |  |
| *Plesiomonas* |  |  |  |  |  |  |
| *Psychrosinus* |  |  |  |  |  |  |
| *Roseiflexus* |  |  |  |  |  |  |
| *Segetibacter* |  |  |  |  |  |  |
| *Succinispira* |  |  |  |  |  |  |
| *Thermomicrobium* |  |  |  |  |  |  |
| *Thioalkalimicrobium* |  |  |  |  | S_2_O_3_^2-^, HS^-^, S^0^, S_4_O_6_^2-^ (Sorokin *et al*., 2001) |  |
| *Thioalkalivibrio* |  |  |  |  | S_2_O_3_^2-^ (Banciu *et al*., 2004) |  |
| *Tistrella* |  |  |  |  |  |  |
| *Acidaminobacter* |  |  |  |  |  |  |
| *Aerococcus* |  |  |  |  |  |  |
| *Aliidiomarina* |  |  |  |  |  |  |
| *Anaerococcus* |  |  |  |  |  |  |
| *Anaeromyxobacter* |  |  |  |  |  |  |
| *Aquaspirillum* |  |  |  |  |  |  |
| *Arcicella* |  |  |  |  |  |  |
| *Bdellovibrio* |  |  |  |  |  |  |
| *Brevibacterium* |  |  |  |  |  |  |
| *Caldicoprobacter* |  |  |  |  |  |  |
| *Castellaniella* |  |  |  |  |  |  |
| *Desulfomonile* |  |  |  |  |  | SO_4_^2-^, SO_3_^2-^, S_2_O_3_^2-^ (DeWeerd *et al*., 1990) |
| *Desulfonatronum* |  |  |  |  |  | SO_4_^2-^, SO_3_^2-^, S_2_O_3_^2-^ (Zhilina *et al*., 2005) |
| *Dyadobacter* |  |  |  |  |  |  |
| *Erysipelothrix* |  |  |  |  |  |  |
| *Faecalibacterium* |  |  |  |  |  |  |
| *Ferruginibacter* |  |  |  |  |  |  |
| *Flectobacillus* |  |  |  |  |  |  |
| *Fusobacterium* |  |  |  |  |  |  |
| *Gaiella* |  |  |  |  |  |  |
| *Gemmobacter* |  |  |  |  |  |  |
| *Halochromatium* |  |  |  |  | HS^-^, S^0^ (Imhoff *et al*., 1998) |  |
| *Heliothrix* |  |  |  |  |  |  |
| *Hyphomonas* |  |  |  |  |  |  |
| *Luteolibacter* |  |  |  |  |  |  |
| *Lysobacter* |  |  |  |  |  |  |
| *Mongoliitalea* |  |  |  |  |  |  |
| *Neochlamydia* |  |  |  |  |  |  |
| *Nesterenkonia* |  |  |  |  |  |  |
| *Olsenella* |  |  |  |  |  |  |
| *Paenisporosarcina* |  |  |  |  |  |  |
| *Prosthecobacter* |  |  |  |  |  |  |
| *Sphaerochaeta* |  |  |  |  |  |  |
| *Stenotrophomonas* |  |  |  |  | S_2_O_3_^2-^ (Unpublished Mandal *et al*) |  |
| *Sulfuritalea* |  |  |  |  | S^0^, S_2_O_3_^2-^ (Kojima and Fukui, 2011) |  |
| *Thiocapsa* |  |  |  |  | HS^-^, S_2_O_3_^2-^, SO_3_^2-^, S^0^ (Puchkova *et al*., 2000) |  |
| *Vampirovibrio* |  |  |  |  |  |  |
| *Verrucomicrobium* |  |  |  |  |  |  |
| *Dietzia* |  |  |  |  |  |  |
| *Corynebacterium* |  |  |  |  |  |  |
| *Methylophilus* |  |  |  |  |  |  |
| *Paracoccus* |  |  |  |  | S_2_O_3_^2-^, S_4_O_6_^2-^, HS^-^, S^0^ (Ghosh *et al*., 2006) |  |

**Table S6.** Complete list of the 316 genera identified in the *Lotus Pond*-*Rulang* ecosystem at 14:30 h of 20 October 2014, together with the information about their presence (green shade) or absence (no shade) in the four sample-sites, and their capabilities, if any, to oxidize or reduce sulfur species.

| **Genus** | **HTP** | **MTP** | **LTP** | **RVW** | **Sulfur species oxidized for chemolithotrophic nutrition (Reference)** | **Sulfur species reduced for anaerobic respiration**  **(Reference)** |
| --- | --- | --- | --- | --- | --- | --- |
| Armatimonadetes_gp7 |  |  |  |  |  |  |
| Gp3 |  |  |  |  |  |  |
| *Pedomicrobium* |  |  |  |  |  |  |
| *Alkanindiges* |  |  |  |  |  |  |
| *Bosea* |  |  |  |  | S_2_O_3_^2-^, S_4_O_6_^2-^ (Das *et al*., 1996) |  |
| *Butyricicoccus* |  |  |  |  |  |  |
| *Chlorophyta* |  |  |  |  |  |  |
| *Desulfobulbus* |  |  |  |  |  | SO_4_^2-^, S_2_O_3_^2-^ (Sorokin *et al*., 2012) |
| *Meiothermus* |  |  |  |  |  |  |
| *Nesterenkonia* |  |  |  |  |  |  |
| *Nitrospira* |  |  |  |  |  |  |
| *Peredibacter* |  |  |  |  |  |  |
| *Thermodesulfobacterium* |  |  |  |  |  | SO_4_^2-^ (Jeanthon *et al*., 2002) |
| *Thioalkalimicrobium* |  |  |  |  | S_2_O_3_^2-^, HS^-^, S^0^, S_4_O_6_^2-^(Sorokin *et al*., 2001) |  |
| *Zoogloea* |  |  |  |  |  |  |
| *Alkaliflexus* |  |  |  |  |  |  |
| *Desulfosporosinus* |  |  |  |  |  | SO_4_^2-^, S^0^, S_2_O_3_^2-^ (Sánchez-Andrea *et al*., 2015) |
| *Fluviicola* |  |  |  |  |  |  |
| *Fusobacterium* |  |  |  |  |  |  |
| *Gemmobacter* |  |  |  |  |  |  |
| *Halanaerobium* |  |  |  |  |  |  |
| *Methylosoma* |  |  |  |  |  |  |
| *Ohtaekwangia* |  |  |  |  |  |  |
| *Pleomorphomonas* |  |  |  |  |  |  |
| *Spirochaeta* |  |  |  |  | S_2_O_3_^2-^, S^0^, HS^-^ (Dubinina *et al*., 2011) |  |
| *Thioalkalivibrio* |  |  |  |  | S_2_O_3_^2-^ (Banciu *et al*., 2004) |  |
| *Vibrio* |  |  |  |  |  |  |
| *Bacillariophyta* |  |  |  |  |  |  |
| *Caldicellulosiruptor* |  |  |  |  |  |  |
| *Enhydrobacter* |  |  |  |  |  |  |
| GpXIII |  |  |  |  |  |  |
| *Hydrogenobacter* |  |  |  |  | S^0^, S_2_O_3_^2-^ (Shima and Suzuki, 1993) |  |
| *Streptophyta* |  |  |  |  |  |  |
| *Tepidimonas* |  |  |  |  |  |  |
| *Thermodesulfovibrio* |  |  |  |  |  | SO_4_^2-^, S_2_O_3_^2-^, SO_3_^2-^ (Henry *et al*., 1994) |
| *Acetobacterium* |  |  |  |  |  |  |
| Acetothermia_genera_incertae_sedis |  |  |  |  |  |  |
| *Acholeplasma* |  |  |  |  |  |  |
| *Algoriphagus* |  |  |  |  |  |  |
| *Aliidiomarina* |  |  |  |  |  |  |
| *Alkalibacter* |  |  |  |  |  |  |
| *Alkalibacterium* |  |  |  |  |  |  |
| *Allochromatium* |  |  |  |  | S^0^, HS^-^, S_2_O_3_^2-^ (Dahl *et al*., 2005) |  |
| Aminicenantes_genera_incertae_sedis |  |  |  |  |  |  |
| *Aquicella* |  |  |  |  |  |  |
| *Arcicella* |  |  |  |  |  |  |
| *Arcobacter* |  |  |  |  | HS^-^ (Sievert *et al*., 2007) |  |
| Atribacteria_genera_incertae_sedis |  |  |  |  |  |  |
| *Azospirillum* |  |  |  |  | S_2_O_3_^2-^, HS^-^ (Lavrinenko *et al*., 2010) |  |
| *Bacteriovorax* |  |  |  |  |  |  |
| *Bacteroides* |  |  |  |  |  |  |
| *Bdellovibrio* |  |  |  |  |  |  |
| *Calditerrivibrio* |  |  |  |  |  |  |
| *Caloramator* |  |  |  |  |  |  |
| *Candidatus Cloacamonas* |  |  |  |  |  |  |
| *Carnobacterium* |  |  |  |  |  |  |
| *Cellvibrio* |  |  |  |  |  |  |
| *Chlorobaculum* |  |  |  |  | S_2_O_3_^2-^, SO_3_^2-^, HS^-^ (Gregersen *et al*., 2011) |  |
| *Chrysiogenes* |  |  |  |  |  |  |
| Clostridium III |  |  |  |  |  |  |
| *Clostridium* sensu stricto |  |  |  |  |  |  |
| *Cryobacterium* |  |  |  |  |  |  |
| *Cryptomonadaceae* |  |  |  |  |  |  |
| *Cytophaga* |  |  |  |  |  |  |
| *Dechloromonas* |  |  |  |  |  |  |
| *Deefgea* |  |  |  |  |  |  |
| *Demequina* |  |  |  |  |  |  |
| *Desulfatirhabdium* |  |  |  |  |  | S_2_O_3_^2-^, SO_4_^2-^ (Balk *et al*., 2008) |
| *Desulfomicrobium* |  |  |  |  |  | SO_4_^2-^, SO_3_^2-^, S_2_O_3_^2-^ (Rozanova *et al*., 1988) |
| *Desulfonatronovibrio* |  |  |  |  |  | SO_4_^2-^, SO_3_^2-^, S_2_O_3_^2-^ (Sorokin *et al*., 2012) |
| *Desulfonatronum* |  |  |  |  |  | SO_4_^2-^, SO_3_^2-^, S_2_O_3_^2-^ (Zhilina *et al*., 2005) |
| *Dethiosulfatibacter* |  |  |  |  |  | S_2_O_3_^2-^, S^0^ (Takii *et al*., 2007) |
| *Dickeya* |  |  |  |  |  |  |
| *Dyadobacter* |  |  |  |  |  |  |
| *Ectothiorhodospira* |  |  |  |  | HS^-^, S^0^ (Imhoff and Trüper, 1981) |  |
| *Emticicia* |  |  |  |  |  |  |
| *Erysipelothrix* |  |  |  |  |  |  |
| *Ferribacterium* |  |  |  |  |  |  |
| *Ferruginibacter* |  |  |  |  |  |  |
| *Fervidicella* |  |  |  |  |  |  |
| *Fibrobacter* |  |  |  |  |  |  |
| *Flavobacterium* |  |  |  |  |  |  |
| *Geobacter* |  |  |  |  |  |  |
| *Giesbergeria* |  |  |  |  |  |  |
| *Gillisia* |  |  |  |  |  |  |
| Gp16 |  |  |  |  |  |  |
| Gp6 |  |  |  |  |  |  |
| Gp7 |  |  |  |  |  |  |
| GpIIa |  |  |  |  |  |  |
| *Gracilimonas* |  |  |  |  |  |  |
| *Haliea* |  |  |  |  |  |  |
| *Halochromatium* |  |  |  |  | HS^-^, S^0^ (Imhoff *et al*., 1998) |  |
| *Hydrogenophaga* |  |  |  |  |  |  |
| *Hydrogenophilus* |  |  |  |  |  |  |
| *Hyphomonas* |  |  |  |  |  |  |
| *Ignavibacterium* |  |  |  |  |  |  |
| *Legionella* |  |  |  |  |  |  |
| *Limnohabitans* |  |  |  |  |  |  |
| *Lysobacter* |  |  |  |  |  |  |
| *Marinomonas* |  |  |  |  |  |  |
| *Megamonas* |  |  |  |  |  |  |
| *Methylobacter* |  |  |  |  |  |  |
| *Methylocystis* |  |  |  |  |  |  |
| *Methylomicrobium* |  |  |  |  |  |  |
| *Methylomonas* |  |  |  |  |  |  |
| *Mongoliitalea* |  |  |  |  |  |  |
| *Niabella* |  |  |  |  |  |  |
| *Nitriliruptor* |  |  |  |  |  |  |
| *Paenisporosarcina* |  |  |  |  |  |  |
| *Paludibacter* |  |  |  |  |  |  |
| *Planomicrobium* |  |  |  |  |  |  |
| *Polaromonas* |  |  |  |  |  |  |
| *Polynucleobacter* |  |  |  |  |  |  |
| *Prevotella* |  |  |  |  |  |  |
| *Proteiniclasticum* |  |  |  |  |  |  |
| *Providencia* |  |  |  |  |  |  |
| *Psychrobacter* |  |  |  |  |  |  |
| *Psychroflexus* |  |  |  |  |  |  |
| *Rahnella* |  |  |  |  |  |  |
| *Rhizobacter* |  |  |  |  |  |  |
| *Rhodobaca* |  |  |  |  |  |  |
| *Rhodobacter* |  |  |  |  | HS^-^ (Hansen and Imhoff, 1985) |  |
| *Rhodoferax* |  |  |  |  |  |  |
| *Roseomonas* |  |  |  |  |  |  |
| *Rubrimonas* |  |  |  |  |  |  |
| *Runella* |  |  |  |  |  |  |
| *Saccharofermentans* |  |  |  |  |  |  |
| *Sediminibacterium* |  |  |  |  |  |  |
| *Shewanella* |  |  |  |  |  |  |
| *Smithella* |  |  |  |  |  |  |
| SR1_genera_incertae_sedis |  |  |  |  |  |  |
| *Sulfuricella* |  |  |  |  | S^0^, S_2_O_3_^2-^ (Kojima and Fukui, 2010) |  |
| *Sulfuricurvum* |  |  |  |  | HS^-^, S^0^, S_2_O_3_^2-^ (Kodama and Watanabe, 2004) |  |
| *Sulfurospirillum* |  |  |  |  |  | S^0^, S_2_O_3_^2-^, SO_3_^2-^ (Kodama *et al.,* 2007) |
| *Sunxiuqinia* |  |  |  |  |  |  |
| *Syntrophorhabdus* |  |  |  |  |  |  |
| *Thermotoga* |  |  |  |  |  |  |
| *Thioalkalibacter* |  |  |  |  | S_2_O_3_^2-^, S^0^, HS^-^ (Banciu *et al*., 2008) |  |
| *Thiobacillus* |  |  |  |  | S_2_O_3_^2-^, S_4_O_6_^2-^, S^0^ (Wood and Kelly., 1991) |  |
| *Thiocapsa* |  |  |  |  | HS^-^, S_2_O_3_^2-^, SO_3_^2-^, S^0^ (Puchkova *et al*., 2000) |  |
| *Thiocystis* |  |  |  |  | HS^-^, S^0^ (Peduzzi *et al*., 2011) |  |
| *Thiomicrospira* |  |  |  |  | S_2_O_3_^2-^, S_4_O_6_^2-^, S^0^ (Sorokin *et al*., 2006) |  |
| *Thiovirga* |  |  |  |  | S_2_O_3_^2-^, S^0^, HS^-^ (Ito *et al*., 2005) |  |
| *Tissierella* |  |  |  |  |  |  |
| *Trichococcus* |  |  |  |  |  |  |
| *Vampirovibrio* |  |  |  |  |  |  |
| *Acidovorax* |  |  |  |  |  |  |
| *Acinetobacter* |  |  |  |  | S_2_O_3_^2-^, S^0^ ( Aguilar *et al*., 2008) |  |
| *Advenella* |  |  |  |  | S_2_O_3_^2-^, S_4_O_6_^2-^ (Ghosh *et al*., 2005) |  |
| *Aeromonas* |  |  |  |  |  |  |
| *Anoxybacillus* |  |  |  |  | S_4_O_6_^2-^ (Unpublished Mandal *et al*) |  |
| *Bacillus* |  |  |  |  | S_2_O_3_^2-^ (Pérez-Ibarra *et al*., 2007) |  |
| *Beijerinckia* |  |  |  |  |  |  |
| *Bradyrhizobium* |  |  |  |  | S_2_O_3_^2-^ (Masuda *et al*., 2010) |  |
| *Brevibacillus* |  |  |  |  |  |  |
| *Brevundimonas* |  |  |  |  |  |  |
| *Burkholderia* |  |  |  |  | S_2_O_3_^2-^, S_4_O_6_^2-^, S^0^, HS^-^ (Wittke *et al*., 1997) |  |
| *Chloroflexus* |  |  |  |  |  |  |
| *Cloacibacterium* |  |  |  |  |  |  |
| Clostridium XlVa |  |  |  |  |  |  |
| *Desulfovibrio* |  |  |  |  |  | SO_3_^2-^, S_2_O_3_^2-^, S^0^ (Mogensen *et al.,* 2005) |
| *Dictyoglomus* |  |  |  |  |  |  |
| *Dietzia* |  |  |  |  |  |  |
| *Escherichia/Shigella* |  |  |  |  |  |  |
| *Fervidobacterium* |  |  |  |  |  |  |
| *Geobacillus* |  |  |  |  |  |  |
| *Geothrix* |  |  |  |  |  |  |
| GpI |  |  |  |  |  |  |
| GpIV |  |  |  |  |  |  |
| *Halomonas* |  |  |  |  | S_2_O_3_^2-^ (Sorokin, 2003) |  |
| *Hyphomicrobium* |  |  |  |  |  |  |
| *Methylobacterium* |  |  |  |  | S_2_O_3_^2-^ (Anandham *et al*., 2007) |  |
| *Nocardioides* |  |  |  |  |  |  |
| *Paracoccus* |  |  |  |  | S_2_O_3_^2-^, S_4_O_6_^2-^, HS^-^, S^0^ (Ghosh *et al*., 2006) |  |
| *Pedobacter* |  |  |  |  |  |  |
| *Planococcus* |  |  |  |  |  |  |
| *Propionibacterium* |  |  |  |  |  |  |
| *Pseudomonas* |  |  |  |  | S_2_O_3_^2-^ (Sorokin *et al*., 1999) |  |
| *Rheinheimera* |  |  |  |  |  |  |
| Saccharibacteria_genera_incertae_sedis |  |  |  |  |  |  |
| Sporolactobacillaceae_incertae_sedis |  |  |  |  |  |  |
| *Sulfurihydrogenibium* |  |  |  |  | S^0^, S_2_O_3_^2-^ (Nakagawa *et al*., 2005) |  |
| *Sulfurimonas* |  |  |  |  | S^0^, HS^-^, S_2_O_3_^2-^ (Inagaki *et al*., 2003) |  |
| *Thermus* |  |  |  |  | S_2_O_3_^2-^, S^0^ (Skirnisdottir *et al*., 2001) |  |
| *Thiofaba* |  |  |  |  | S_2_O_3_^2-^, S^0^, HS^-^, S_4_O_6_^2-^ (Mori and Suzuki, 2008) |  |
| *Thiothrix* |  |  |  |  | S_2_O_3_^2-^, HS^-^ (Chernousova *et al*., 2009) |  |
| *Treponema* |  |  |  |  |  |  |
| *Brevibacterium* |  |  |  |  |  |  |
| *Chryseobacterium* |  |  |  |  |  |  |
| *Comamonas* |  |  |  |  |  |  |
| *Corynebacterium* |  |  |  |  |  |  |
| *Deinococcus* |  |  |  |  |  |  |
| *Methyloversatilis* |  |  |  |  |  |  |
| *Pelomonas* |  |  |  |  |  |  |
| *Serratia* |  |  |  |  |  |  |
| *Acidaminobacter* |  |  |  |  |  |  |
| *Anaeromyxobacter* |  |  |  |  |  |  |
| *Campylobacter* |  |  |  |  | S_2_O_3_^2-^, S_4_O_6_^2-^, HS^-^ (Voordouw *et al*., 1996) |  |
| Clostridium XI |  |  |  |  |  |  |
| *Curvibacter* |  |  |  |  |  |  |
| *Flectobacillus* |  |  |  |  |  |  |
| *Formivibrio* |  |  |  |  |  |  |
| *Gemmatimonas* |  |  |  |  |  |  |
| *Luteolibacter* |  |  |  |  |  |  |
| *Methylophilus* |  |  |  |  |  |  |
| *Parabacteroides* |  |  |  |  |  |  |
| *Pseudoxanthomonas* |  |  |  |  |  |  |
| *Rickettsia* |  |  |  |  |  |  |
| *Sphaerochaeta* |  |  |  |  |  |  |
| *Steroidobacter* |  |  |  |  |  |  |
| *Sulfuritalea* |  |  |  |  | S^0^, S_2_O_3_^2-^ (Kojima and Fukui, 2011) |  |
| *Thiobaca* |  |  |  |  | HS^-^ (Rees *et al*., 2002) |  |
| *Delftia* |  |  |  |  |  |  |
| *Litoreibacter* |  |  |  |  |  |  |
| *Paenibacillus* |  |  |  |  |  |  |
| *Rhodoplanes* |  |  |  |  |  |  |
| *Staphylococcus* |  |  |  |  |  |  |
| *Achromobacter* |  |  |  |  |  |  |
| *Actinomyces* |  |  |  |  |  |  |
| *Alcanivorax* |  |  |  |  |  |  |
| Armatimonadetes_gp5 |  |  |  |  |  |  |
| *Brachybacterium* |  |  |  |  |  |  |
| *Cruoricaptor* |  |  |  |  |  |  |
| *Finegoldia* |  |  |  |  |  |  |
| *Gemella* |  |  |  |  |  |  |
| Gp2 |  |  |  |  |  |  |
| *Klebsiella* |  |  |  |  | S_2_O_3_^2-^ (Mason and Kelly, 1988) |  |
| *Lysinibacillus* |  |  |  |  |  |  |
| *Marinococcus* |  |  |  |  |  |  |
| *Marmoricola* |  |  |  |  |  |  |
| *Microbacterium* |  |  |  |  | S_2_O_3_^2-^(Anandham *et al*., 2008) |  |
| *Nevskia* |  |  |  |  |  |  |
| *Patulibacter* |  |  |  |  |  |  |
| *Peptoniphilus* |  |  |  |  |  |  |
| *Photobacterium* |  |  |  |  |  |  |
| *Ralstonia* |  |  |  |  |  |  |
| *Rubrobacter* |  |  |  |  |  |  |
| *Sphingobium* |  |  |  |  |  |  |
| *Sphingomonas* |  |  |  |  |  |  |
| *Sporolituus* |  |  |  |  |  |  |
| *Stenotrophomonas* |  |  |  |  | S_2_O_3_^2-^ (Unpublished Mandal *et al*) |  |
| *Sulfitobacter* |  |  |  |  | S_2_O_3_^2-^, SO_3_^2-^ (Pukall *et al*., 1999) |  |
| *Thermoactinomyces* |  |  |  |  |  |  |
| *Truepera* |  |  |  |  |  |  |
| *Turicella* |  |  |  |  |  |  |
| *Variovorax* |  |  |  |  |  |  |
| *Aerococcus* |  |  |  |  |  |  |
| *Alkalilimnicola* |  |  |  |  | HS^-^, S_2_O_3_^2-^ (Hoeft *et al*., 2007) |  |
| *Aminiphilus* |  |  |  |  |  |  |
| *Aquimonas* |  |  |  |  |  |  |
| *Azospira* |  |  |  |  |  |  |
| *Caldilinea* |  |  |  |  |  |  |
| Clostridium XVIII |  |  |  |  |  |  |
| *Desulforegula* |  |  |  |  |  | SO_4_^2-^ (Rees and Patel, 2001) |
| *Euzebya* |  |  |  |  |  |  |
| Gp4 |  |  |  |  |  |  |
| *Holophaga* |  |  |  |  |  |  |
| *Lactobacillus* |  |  |  |  |  |  |
| *Leptospira* |  |  |  |  |  |  |
| *Marinospirillum* |  |  |  |  |  |  |
| *Mesorhizobium* |  |  |  |  | S_2_O_3_^2-^, S^0^ (Ghosh and Roy, 2006) |  |
| *Methylovulum* |  |  |  |  |  |  |
| *Rhodocyclus* |  |  |  |  | SO_3_^2-^ (Pfennig 1978) |  |
| *Sphingopyxis* |  |  |  |  |  |  |
| *Actinoalloteichus* |  |  |  |  |  |  |
| *Aquaspirillum* |  |  |  |  |  |  |
| *Arenimonas* |  |  |  |  |  |  |
| *Asticcacaulis* |  |  |  |  |  |  |
| *Azonexus* |  |  |  |  |  |  |
| *Desulfococcus* |  |  |  |  |  | SO_4_^2-^, SO_3_^2-^ (Platen *et al*., 1990) |
| Erysipelotrichaceae_incertae_sedis |  |  |  |  |  |  |
| *Faecalibacterium* |  |  |  |  |  |  |
| GpVIII |  |  |  |  |  |  |
| *Lactococcus* |  |  |  |  |  |  |
| *Oceanibaculum* |  |  |  |  |  |  |
| *Opitutus* |  |  |  |  |  |  |
| *Plesiomonas* |  |  |  |  |  |  |
| *Rhodospirillum* |  |  |  |  | S_2_O_3_^2-^, HS^-^ (Kumar *et al*., 2008) |  |
| *Roseococcus* |  |  |  |  | S_2_O_3_^2-^ (Yurkov *et al*., 1994) |  |
| *Rubrivivax* |  |  |  |  |  |  |
| *Salinicoccus* |  |  |  |  |  |  |
| Subdivision3_genera_incertae_sedis |  |  |  |  |  |  |
| *Thermoleophilum* |  |  |  |  |  |  |
| *Alishewanella* |  |  |  |  |  |  |
| *Alkalibaculum* |  |  |  |  |  |  |
| *Alkaliphilus* |  |  |  |  |  |  |
| *Anaerobiospirillum* |  |  |  |  |  |  |
| *Aquabacterium* |  |  |  |  |  |  |
| *Blastocatella* |  |  |  |  |  |  |
| *Brachymonas* |  |  |  |  |  |  |
| BRC1_genera_incertae_sedis |  |  |  |  |  |  |
| *Candidatus Endomicrobium* |  |  |  |  |  |  |
| *Dethiobacter* |  |  |  |  |  |  |
| *Friedmanniella* |  |  |  |  |  |  |
| *Haliscomenobacter* |  |  |  |  |  |  |
| *Ilumatobacter* |  |  |  |  |  |  |
| *Jeotgalicoccus* |  |  |  |  |  |  |
| Latescibacteria_genera_incertae_sedis |  |  |  |  |  |  |
| *Loktanella* |  |  |  |  |  |  |
| *Magnetococcus* |  |  |  |  | S_2_O_3_^2-^, SO_3_^2-^ (Bazylinski *et al*., 2013) |  |
| *Marinimicrobium* |  |  |  |  |  |  |
| *Ornithinimicrobium* |  |  |  |  |  |  |
| *Piscibacillus* |  |  |  |  |  |  |
| *Pseudoflavonifractor* |  |  |  |  |  |  |
| *Pseudorhodobacter* |  |  |  |  |  |  |
| *Rhizobium* |  |  |  |  |  |  |
| *Sphingobacterium* |  |  |  |  |  |  |
| *Terrimonas* |  |  |  |  |  |  |
| *Thermanaerovibrio* |  |  |  |  |  |  |
| ZB3_genera_incertae_sedis |  |  |  |  |  |  |
| *Dehalogenimonas* |  |  |  |  |  |  |
| *Marinobacter* |  |  |  |  |  |  |
| *Mycobacterium* |  |  |  |  |  |  |
| *Streptococcus* |  |  |  |  |  |  |
| *Thermomonas* |  |  |  |  |  |  |
| *Exiguobacterium* |  |  |  |  |  |  |

**References used in Tables S5 and S6**

Aguilar JRP, Cabriales JJP, Vega MM. Identification and characterization of sulfur-oxidizing bacteria in an artificial wetland that treats wastewater from a tannery. Int J Phytorem. 2008;10:359-370.

Anandham R, Indiragandhi P, Madhaiyan M, Kim K, Yim W, Saravanan V, et al. Thiosulfate oxidation and mixotrophic growth of *Methylobacterium oryzae*. Can J Microbiol. 2007;53:869-876.

Anandham R, Indiragandhi P, Madhaiyan M, Ryu KY, Jee HJ, Sa TM. Chemolithoautotrophic oxidation of thiosulfate and phylogenetic distribution of sulfur oxidation gene (soxB) in *rhizobacteria* isolated from crop plants. Res Microbiol. 2008;159:579-589.

Balk M, Altınbaş M, Rijpstra WIC, Damste JSS, Stams AJ. *Desulfatirhabdium butyrativorans* gen. nov., sp. nov., a butyrate-oxidizing, sulfate-reducing bacterium isolated from an anaerobic bioreactor. Int J Syst Evol Microbiol. 2008;58:110-115.

Banciu H, Sorokin DY, Galinski EA, Muyzer G, Kleerebezem R, Kuenen JG. *Thialkalivibrio halophilus* sp. nov., a novel obligately chemolithoautotrophic, facultatively alkaliphilic, and extremely salt-tolerant, sulfur-oxidizing bacterium from a hypersaline alkaline lake. Extremophiles. 2004;8:325-334.

Banciu HL, Sorokin DY, Tourova TP, Galinski EA, Muntyan MS, Kuenen JG, et al. Influence of salts and pH on growth and activity of a novel facultatively alkaliphilic, extremely salt-tolerant, obligately chemolithoautotrophic sufur-oxidizing Gammaproteobacterium *Thioalkalibacter halophilus* gen. nov., sp. nov. from South-Western Siberian soda lakes. Extremophiles. 2008;12:391-404.

Bazylinski DA, Williams TJ, Lefevre CT, Berg RJ, Zhang CL, Bowser SS, et al. *Magnetococcus marinus* gen. nov., sp. nov., a marine, magnetotactic bacterium that represents a novel lineage (*Magnetococcaceae fam*. nov., *Magnetococcales ord*. nov.) at the base of the Alphaproteobacteria. Int J Syst Evol Microbiol. 2013;63:801-808.

Chernousova E, Gridneva E, Grabovich M, Dubinina G, Akimov V, Rossetti S, et al. *Thiothrix caldifontis* sp. nov. and *Thiothrix lacustris* sp. nov., gammaproteobacteria isolated from sulfide springs. Int J Syst Evol Microbiol. 2009;59:3128-3135.

Dahl C, Engels S, Pott-Sperling AS, Schulte A, Sander J, Lübbe Y, et al. Novel genes of the dsr gene cluster and evidence for close interaction of Dsr proteins during sulfur oxidation in the phototrophic sulfur bacterium *Allochromatium vinosum*. J Bacteriol. 2005;187:1392-1404.

Das SK, Mishra AK, Tindall BJ, Rainey FA, Stackebrandt E. Oxidation of thiosulfate by a new bacterium, *Bosea thiooxidans*.(strain BI-42) gen. nov., sp. nov.: analysis of phylogeny based on chemotaxonomy and 16S ribosomal DNA sequencing. Int J Syst Evol Microbiol. 1996;46:981-987.

DeWeerd KA, Mandelco L, Tanner RS, Woese CR, Suflita JM. *Desulfomonile tiedjei* gen. nov. and sp. nov., a novel anaerobic, dehalogenating, sulfate-reducing bacterium. Arch Microbiol. 1990;154:23-30.

Dubinina G, Grabovich M, Leshcheva N, Rainey FA, Gavrish E. *Spirochaeta perfilievii* sp. nov., an oxygen-tolerant, sulfide-oxidizing, sulfur-and thiosulfate-reducing spirochaete isolated from a saline spring. Int J Syst Evol Microbiol. 2011;61:110-117.

Ghosh W, Roy P. *Mesorhizobium thiogangeticum* sp. nov., a novel sulfur-oxidizing chemolithoautotroph from rhizosphere soil of an Indian tropical leguminous plant. Int J Syst Evol Microbiol. 2006;56:91-97.

Ghosh W, Bagchi A, Mandal S, Dam B, Roy P. *Tetrathiobacter kashmirensis* gen. nov., sp. nov., a novel mesophilic, neutrophilic, tetrathionate-oxidizing, facultatively chemolithotrophic betaproteobacterium isolated from soil from a temperate orchard in Jammu and Kashmir, India. Int J Syst Evol Microbiol. 2005;55:1779-1787.

Ghosh W, Mandal S, Roy P. *Paracoccus bengalensis* sp. nov., a novel sulfur-oxidizing chemolithoautotroph from the rhizospheric soil of an Indian tropical leguminous plant. Syst Appl Microbiol. 2006;29:396-403.

Gregersen LH, Bryant DA, Frigaard NU. Mechanisms and evolution of oxidative sulfur metabolism in green sulfur bacteria. Front Microbiol. 2011;2:116.

Hansen T, Imhoff JF. *Rhodobacter veldkampii*, a new species of phototrophic purple nonsulfur bacteria. Int J Syst Evol Microbiol. 1985;35:115-116.

Henry E, Devereux R, Maki J, Gilmour C, Woese C, Mandelco L, et al. Characterization of a new thermophilic sulfate-reducing bacterium. Arch Microbiol. 1994;161:62-69.

Hoeft SE, Blum JS, Stolz JF, Tabita FR, Witte B, King GM, et al. *Alkalilimnicola ehrlichii* sp. nov., a novel, arsenite-oxidizing haloalkaliphilic gammaproteobacterium capable of chemoautotrophic or heterotrophic growth with nitrate or oxygen as the electron acceptor. Int J Syst Evol Microbiol. 2007;57:504-512.

Imhoff JF, Trüper, H.G. *Ectothiorhodospira abdelmalekii* sp. nov., a new halophilic and alkaliphilic phototrophic bacterium. Zentralblatt für Bakteriologie Mikrobiologie und Hygiene: I. Abt. Originale C: Allgemeine, angewandte und ökologische Mikrobiologie. 1981;2:228-234.

Imhoff JF, Süling J, PETRI R. Phylogenetic relationships among the Chromatiaceae, their taxonomic reclassification and description of the new genera *Allochromatium, Halochromatium, Isochromatium, Marichromatium, Thiococcus, Thiohalocapsa* and *Thermochromatium*. Int J Syst Evol Microbiol. 1998;48:1129-1143.

Inagaki F, Takai K, Kobayashi H, Nealson KH, Horikoshi K. *Sulfurimonas autotrophica* gen. nov., sp. nov., a novel sulfur-oxidizing ε-proteobacterium isolated from hydrothermal sediments in the Mid-Okinawa Trough. Int J Syst Evol Microbiol. 2003;53:1801-1805.

Ito T, Sugita K, Yumoto I, Nodasaka Y, Okabe S. *Thiovirga sulfuroxydans* gen. nov., sp. nov., a chemolithoautotrophic sulfur-oxidizing bacterium isolated from a microaerobic waste-water biofilm. Int J Syst Evol Microbiol. 2005;55:1059-1064.

Jeanthon C, L'Haridon S, Cueff V, Banta A, Reysenbach AL, Prieur D. *Thermodesulfobacterium hydrogeniphilum* sp. nov., a thermophilic, chemolithoautotrophic, sulfate-reducing bacterium isolated from a deep-sea hydrothermal vent at Guaymas Basin, and emendation of the genus *Thermodesulfobacterium*. Int J Syst Evol Microbiol. 2002;52:765-772.

Kodama Y, Watanabe K. *Sulfuricurvum kujiense* gen. nov., sp. nov., a facultatively anaerobic, chemolithoautotrophic, sulfur-oxidizing bacterium isolated from an underground crude-oil storage cavity. Int J Syst Evol Microbiol. 2004;54:2297-2300.

Kodama Y, Watanabe K. *Sulfurospirillum cavolei* sp. nov., a facultatively anaerobic sulfur-reducing bacterium isolated from an underground crude oil storage cavity. Int J Syst Evol Microbiol. 2007;57:827-831.

Kojima H, Fukui M. *Sulfuricella denitrificans* gen. nov., sp. nov., a sulfur-oxidizing autotroph isolated from a freshwater lake. Int J Syst Evol Microbiol. 2010;60:2862-2866.

Kojima H, Fukui M. *Sulfuritalea hydrogenivorans* gen. nov., sp. nov., a facultative autotroph isolated from a freshwater lake. Int J Syst Evol Microbiol. 2011;61:1651-1655.

Kumar PA, Aparna P, Srinivas T, Sasikala C, Ramana CV. *Rhodospirillum sulfurexigens* sp. nov., a phototrophic alphaproteobacterium requiring a reduced sulfur source for growth. Int J Syst Evol Microbiol. 2008;58:2917-2920.

Lavrinenko K, Chernousova E, Gridneva E, Dubinina G, Akimov V, Kuever J, et al. *Azospirillum thiophilum* sp. nov., a diazotrophic bacterium isolated from a sulfide spring. Int J Syst Evol Microbiol. 2010;60:2832-2837.

Mason J, Kelly DP. Thiosulfate oxidation by obligately heterotrophic bacteria. Microb Ecol. 1988;15:123-134.

Masuda S, Eda S, Sugawara C, Mitsui H, Minamisawa K. The cbbL gene is required for thiosulfate-dependent autotrophic growth of *Bradyrhizobium japonicum*. Microbes Environ. 2010;25:220-223.

Mogensen GL, Kjeldsen KU, Ingvorsen K. *Desulfovibrio aerotolerans* sp. nov., an oxygen tolerant sulphate-reducing bacterium isolated from activated sludge. Anaerobe. 2005;11:339-349.

Mori K, Suzuki KI. *Thiofaba tepidiphila* gen. nov., sp. nov., a novel obligately chemolithoautotrophic, sulfur-oxidizing bacterium of the Gammaproteobacteria isolated from a hot spring. Int J Syst Evol Microbiol. 2008;58:1885-1891.

Nakagawa S, Shtaih Z, Banta A, Beveridge T, Sako Y, Reysenbach AL. *Sulfurihydrogenibium yellowstonense* sp. nov., an extremely thermophilic, facultatively heterotrophic, sulfur-oxidizing bacterium from Yellowstone National Park, and emended descriptions of the genus *Sulfurihydrogenibium, Sulfurihydrogenibium subterraneum* and *Sulfurihydrogenibium azorense*. Int J Syst Evol Microbiol. 2005;55:2263-2268.

Peduzzi S, Welsh A, Demarta A, Decristophoris P, Peduzzi R, Hahn D, et al. *Thiocystis chemoclinalis* sp. nov. and *Thiocystis cadagnonensis* sp. nov., motile purple sulfur bacteria isolated from the chemocline of a meromictic lake. Int J Syst Evol Microbiol. 2011;61:1682-1687.

Pérez-Ibarra BM, Flores ME, García-Varela M. Isolation and characterization of *Bacillus thioparus* sp. nov., chemolithoautotrophic, thiosulfate-oxidizing bacterium. FEMS Microbiol Lett. 2007;271:289-296.

Pfennig N. *Rhodocyclus purpureus* gen. nov. and sp. nov., a ring-shaped, vitamin B12-requiring member of the family Rhodospirillaceae. Int J Syst Evol Microbiol. 1978;28:283-288.

Platen H, Temmes A, Schink B. Anaerobic degradation of acetone by *Desulfococcus biacutus* spec. nov. Arch Microbiol. 1990;154:355-361.

Puchkova NN, Imhoff JF, Gorlenko VM. *Thiocapsa litoralis* sp. nov., a new purple sulfur bacterium from microbial mats from the White Sea. Int J Syst Evol Microbiol. 2000;50:1441-1447.

Pukall R, Buntefuβ D, Frühling A, Rohde M, Kroppenstedt RM, Burghardt J, et al. *Sulfitobacter mediterraneus* sp. nov., a new sulfite-oxidizing member of the α-Proteobacteria. Int J Syst Evol Microbiol. 1999;49:513-519.

Rees GN, Patel B. *Desulforegula conservatrix* gen. nov., sp. nov., a long-chain fatty acid-oxidizing, sulfate-reducing bacterium isolated from sediments of a freshwater lake. Int J Syst Evol Microbiol. 2001;51:1911-1916.

Rees GN, Harfoot CG, Janssen PH, Schoenborn L, Kuever J, Lünsdorf H. *Thiobaca trueperi* gen. nov., sp. nov., a phototrophic purple sulfur bacterium isolated from freshwater lake sediment. Int J Syst Evol Microbiol. 2002;52:671-678.

Rozanova E, Nazina T, Galushko A. Isolation of a new genus of sulfate-reducing bacteria and description of a new species of this genus, *Desulfomicrobium apsheronum* gen. nov., sp. nov. Microbiology. 1988;57:514-520.

Sánchez-Andrea I, Stams AJ, Hedrich S, Ňancucheo I, Johnson DB. *Desulfosporosinus acididurans* sp. nov.: an acidophilic sulfate-reducing bacterium isolated from acidic sediments. Extremophiles. 2015;19:39-47.

Shima S, Suzuki KI. *Hydrogenobacter acidophilus* sp. nov., a thermoacidophilic, aerobic, hydrogen-oxidizing bacterium requiring elemental sulfur for growth. Int J Syst Evol Microbiol. 1993;43:703-708.

Sievert SM, Wieringa EB, Wirsen CO, Taylor CD. Growth and mechanism of filamentous‐sulfur formation by *Candidatus Arcobacter sulfidicus* in opposing oxygen‐sulfide gradients. Environ Microbiol. 2007;9:271-276.

Skirnisdottir S, Hreggvidsson GO, Holst O, Kristjansson JK. Isolation and characterization of a mixotrophic sulfur-oxidizing *Thermus scotoductus*. Extremophiles. 2001;5:45-51.

Sorokin D, Tourova T, Abbas B, Suhacheva M, Muyzer G. *Desulfonatronovibrio halophilus* sp. nov., a novel moderately halophilic sulfate-reducing bacterium from hypersaline chloride–sulfate lakes in Central Asia. Extremophiles. 2012;16:411-417.

Sorokin DY, Lysenko AM, Mityushina LL, Tourova TP, Jones BE, Rainey FA, et al. *Thioalkalimicrobium aerophilum* gen. nov., sp. nov. and *Thioalkalimicrobium sibericum* sp. nov., and *Thioalkalivibrio versutus* gen. nov., sp. nov., *Thioalkalivibrio nitratis* sp. nov., novel and *Thioalkalivibrio denitrificancs* sp. nov., novel obligately alkaliphilic and obligately chemolithoautotrophic sulfur-oxidizing bacteria from soda lakes. Int J Syst Evol Microbiol. 2001;51:565-580.

Sorokin DY, Teske A, Robertson LA, Kuenen JG. Anaerobic oxidation of thiosulfate to tetrathionate by obligately heterotrophic bacteria, belonging to the *Pseudomonas stutzeri* group. FEMS Microbiol Ecol. 1999;30:113-123.

Sorokin DY, Tourova TP, Kolganova TV, Spiridonova EM, Berg IA, Muyzer G. *Thiomicrospira halophila* sp. nov., a moderately halophilic, obligately chemolithoautotrophic, sulfur-oxidizing bacterium from hypersaline lakes. Int J Syst Evol Microbiol. 2006;56:2375-2380.

Sorokin DY, Tourova TP, Panteleeva AN, Muyzer G. *Desulfonatronobacter acidivorans* gen. nov., sp. nov. and *Desulfobulbus alkaliphilus* sp. nov., haloalkaliphilic heterotrophic sulfate-reducing bacteria from soda lakes. Int J Syst Evol Microbiol. 2012;62:2107-2113.

Sorokin DY. Oxidation of inorganic sulfur compounds by obligately organotrophic bacteria. Microbiology. 2003;72:641-653.

Takii S, Hanada S, Tamaki H, Ueno Y, Sekiguchi Y, Ibe A, et al. *Dethiosulfatibacter aminovorans* gen. nov., sp. nov., a novel thiosulfate-reducing bacterium isolated from coastal marine sediment via sulfate-reducing enrichment with Casamino acids. Int J Syst Evol Microbiol. 2007;57:2320-2326.

Vesteinsdottir H, Reynisdottir DB, Örlygsson J. *Thiomonas islandica* sp. nov., a moderately thermophilic, hydrogen-and sulfur-oxidizing betaproteobacterium isolated from a hot spring. Int J Syst Evol Microbiol. 2011;61:132-137.

Voordouw G, Armstrong SM, Reimer MF, Fouts B, Telang AJ, Shen Y, et al. Characterization of 16S rRNA genes from oil field microbial communities indicates the presence of a variety of sulfate-reducing, fermentative, and sulfide-oxidizing bacteria. Appl Environ Microbiol. 1996;62:1623-1629.

Wittke R, Ludwig W, Peiffer S, Kleiner D. Isolation and characterization of *Burkholderia norimbergensis* sp. nov., a mildly alkaliphilic sulfur oxidizer. Syst Appl Microbiol. 1997;20:549-553.

Wood AP, Kelly DP. Isolation and characterisation of *Thiobacillus halophilus* sp. nov., a sulphur-oxidising autotrophic eubacterium from a Western Australian hypersaline lake. Arch Microbiol. 1991;156:277-280.

Yurkov V, Stackebrandt E, Holmes A, Fuerst JA, Hugenholtz P, Golecki J, et al. Phylogenetic positions of novel aerobic, bacteriochlorophyll a-containing bacteria and description of *Roseococcus thiosulfatophilus* gen. nov., sp. nov., *Erythromicrobium ramosum* gen. nov., sp. nov., and *Erythrobacter litoralis* sp. nov. Int J Syst Evol Microbiol. 1994;44:427-434.

Zhilina TN, Zavarzina DG, Kuever J, Lysenko AM, Zavarzin GA. *Desulfonatronum cooperativum* sp. nov., a novel hydrogenotrophic, alkaliphilic, sulfate-reducing bacterium, from a syntrophic culture growing on acetate. Int J Syst Evol Microbiol. 2005;55:1001-1006.

**Table S7.** Concentrations of various redox species of sulfur, and number of sulfur-oxidizing and sulfur-reducing genera present, at the different sample-sites of the *Lotus Pond*-*Rulang* ecosystem at 8:30 and 14:30 h of 20 October 2014.

|  |  | **8:30 h** | | | | **14:30 h** | | | |
| --- | --- | --- | --- | --- | --- | --- | --- | --- | --- |
| **Serial**  **no.** |  | **HTP** | **MTP** | **LTP** | **RVW** | **HTP** | **MTP** | **LTP** | **RVW** |
| **1** | **pH** | 7.5 | 7.4 | 7.2 | 6.6 | 7.8 | 7.5 | 6.8 | 6.2 |
| **2** | **S^2-^ concentration* in the water (in mM)** | 262 | 120 | 0 | 0 | 178 | 93 | 0 | 0 |
| **3** | **δ^34^S of the dissolved sulfide (in ‰ VCDT)** | 4.6 | 7.0 | NA | NA | 4.3 | 5.2 | NA | NA |
| **4** | **S_2_O_3_^2-^ concentration in the water (in mM)** | 1.3 | 1.4 | 1.5 | 0.8 | 0.8 | 0.9 | 1 | 0.5 |
| **5** | **SO_3_^2-^ concentration in the water (in µM)** | 273 | 140 | 0 | 0 | 110 | 15 | 0 | 0 |
| **6** | **SO_4_^2-^ concentration in the water (in µM)** | 712 | 920 | 1152 | 688 | 963 | 1115 | 1281 | 722 |
| **7** | **δ^34^S of the dissolved sulfate (in ‰ VCDT)** | 16.3 | 16.1 | 16.2 | 12.4 | 16.5 | 16.5 | 16.6 | 13.1 |
| **8** | **Number of S^2-^, S^0^, S_2_O_3_^2-^ and/or SO_3_^2-^ oxidizing genera present** | 17 | 26 | 27 | 30 | 20 | 39 | 37 | 36 |
| **9** | **Number of S^2-^ oxidizing genera present** | 6 | 12 | 13 | 15 | 5 | 21 | 20 | 20 |
| **10** | **Number of S^0^ oxidizing genera present** | 9 | 13 | 13 | 17 | 8 | 22 | 21 | 20 |
| **11** | **Number of S_2_O_3_^2-^ oxidizing genera present** | 16 | 23 | 23 | 25 | 19 | 31 | 31 | 29 |
| **12** | **Number of SO_3_^2-^ oxidizing genera present** | 0 | 1 | 1 | 2 | 1 | 3 | 2 | 3 |
| **13** | **Number of S_2_O_3_^2-^, SO_3_^2-^ and/or SO_4_^2-^ reducing genera present** | 0 | 8 | 8 | 10 | 2 | 11 | 12 | 8 |
| **14** | **Number of S_2_O_3_^2-^ reducing genera present** | 0 | 6 | 7 | 9 | 2 | 9 | 10 | 8 |
| **15** | **Number of SO_3_^2-^ reducing genera present** | 0 | 3 | 4 | 6 | 2 | 6 | 7 | 5 |
| **16** | **Number of SO_4_^2-^ reducing genera present** | 0 | 7 | 5 | 7 | 1 | 8 | 9 | 5 |

* Dissolved sulfide measured collectively using the methylene blue colorimetric method included H_2_S, HS‾ and S_x_^2^‾.

NA = Not applicable.

**Table S8.** The 57 *Lotus Pond* genera that were detected exclusively at HTP and/or MTP, at 8:30 and/or 14:30 h of 20 October 2014; information^1^ about the upper temperature-limit for their laboratory growth is also given.

| **Phyla** | **Genera** | **Upper limit of temperature (in °C) for growth *in vitro*** | **Reference** |
| --- | --- | --- | --- |
| **Genera having no report of laboratory growth above 45°C** | | | |
| *Acidobacteria* | *Holophaga* | 35 | (Liesack *et al.,* 1994) |
| *Actinobacteria* | *Actinomycetospora* | 30 | (Yamamura *et al.,* 2011) |
|  | *Arthrobacter* | 45 | (Dastager *et al.,* 2014) |
|  | *Euzebya* | 35 | (Kurahashi *et al.,* 2010) |
|  | *Leifsonia* | 28 | (Evtushenko *et al.,* 2000) |
|  | *Microbacterium* | 42 | (Krishnamurthi *et al.,* 2012) |
|  | *Patulibacter* | 39 | (Almeida *et al.,* 2013) |
|  | *Turicella* | 37 | (Funke *et al.,* 1994) |
| *Bacteroidetes* | *Cruoricaptor* | 37 | (Yassin *et al.,* 2012) |
|  | *Porphyromonas* | 37 | (Willems and Collins, 1995) |
|  | *Planobacterium* | 37 | (Peng *et al.,* 2009) |
| *Firmicutes* | *Finegoldia* | 37 | (Murdoch and Shah, 1999) |
|  | *Gallicola* | 37 | (Ezaki, 2015) |
|  | *Gemella* | 37 | (Kilpper-Bälz and Schleifer, 1988) |
|  | *Lactobacillus* | 45 | (Kudo *et al.,* 2012) |
|  | *Lysinibacillus* | 45 | (Ahmed *et al.,* 2007) |
|  | *Peptoniphilus* | 43 | (Patel *et al.,* 2016) |
| *Proteobacteria* | *Achromobacter* | 40 | (Gomila *et al.,* 2011) |
|  | *Alcanivorax* | 40 | (Rahul *et al.,* 2014) |
|  | *Aquimonas* | 42 | (Saha *et al.,* 2005) |
|  | *Azospira* | 37 | (Lin *et al.,* 2013) |
|  | *Desulforegula* | 32 | (Rees and Patel, 2001) |
|  | *Duganella* | 30 | (Li *et al.,* 2004) |
|  | *Herminiimonas* | 37 | (Kämpfer *et al.,* 2013) |
|  | *Klebsiella* | 37 | (Bergey *et al.,* 1923) |
|  | *Mesorhizobium* | 37 | (Martínez-Hidalgo *et al.,* 2015) |
|  | *Methylovulum* | 34 | (Iguchi *et al.,* 2011) |
|  | *Microvirga* | 45 | (Kanso and Patel, 2003) |
|  | *Nevskia* | 25 | (Leandro *et al.,* 2012) |
|  | *Panacagrimonas* | 42 | (Im *et al.,* 2010) |
|  | *Pedomicrobium* | 40 | (Gebers, 1981) |
|  | *Piscinibacter* | 42 | (Stackebrandt *et al.,* 2009) |
|  | *Ralstonia* | 41 | (Yabuuchi *et al.,* 1995) |
|  | *Rhodocyclus* | 30 | (Pfennig, 1978) |
|  | *Sneathiella* | 37 | (Jordan *et al.,* 2007) |
|  | *Sphaerotilus* | 40 | (Gridneva *et al.,* *2011*) |
|  | *Sphingomonas* | 40 | (Yabuuchi *et al.,* 1990) |
|  | *Sphingopyxis* | 35 | (Baik *et al.,* 2013) |
|  | *Sulfitobacter* | 35 | (Hong *et al.,* 2015) |
|  | *Variovorax* | 30 | (Willems *et al.,* 1991) |
| *Spirochaetes* | *Leptospira* | 30 | (Faine and Stallmn, 1982) |
| **Genera having report of laboratory growth above 45°C** | | | |
| *Actinobacteria* | *Actinomyces* | 55 | (An *et al.,* 2006) |
|  | *Rubrobacter* | 60 | (Carreto *et al.,* 1996) |
| *Deinococcus-Thermus* | *Truepera* | 50 | (Albuquerque *et al.,* 2005) |
| *Firmicutes* | *Marinococcus* | 50 | (Balderrama-Subieta *et al.,* 2013) |
|  | *Sporolituus* | 60 | (Ogg and patel, 2009) |
|  | *Thermoactinomyces* | 60 | (Yao *et al.,* 2014) |
|  | *Tindallia* | 48 | (Pikuta *et al.,* 2003) |
| *Proteobacteria* | *Alkalilimnicola* | 55 | (Yakimov *et al.,* 2001) |
|  | *Marinospirillum* | 55 | (Namsaraev *et al.,* 2009) |
|  | *Photobacterium* | 50 | (Wang *et al.,* 2017) |
|  | *Thiomonas* | 50 | (Moreira *et al.,* 1997) |

^1^	Armatimonadetes_gp5, Acidobacteria Gp2, Acidobacteria Gp4, Parcubacteria_genera_incertae_sedis and Clostridium XVIII do not have any comparable information.

**References used in Table S8**

Ahmed I, Yokota A, Yamazoe A, Fujiwara T. Proposal of *Lysinibacillus boronitolerans* gen. nov. sp. nov., and transfer of *Bacillus fusiformis* to *Lysinibacillus fusiformis* comb. nov. and *Bacillus sphaericus* to *Lysinibacillus sphaericus* comb. nov. Int J Syst Evol Microbiol. 2007;57:1117-1125.

Albuquerque L, Simoes C, Nobre MF, Pino NM, Battista JR, Silva MT, et al. *Truepera radiovictrix gen*. nov., sp. nov., a new radiation resistant species and the proposal of Trueperaceae fam. nov. FEMS Microbiol Lett. 2005;247:161-169.

Almeida B, Vaz-Moreira I, Schumann P, Nunes OC, Carvalho G, Crespo MTB. *Patulibacter medicamentivorans* sp. nov., isolated from activated sludge of a wastewater treatment plant. Int J Syst Evol Microbiol. 2013;63:2588-2593.

An D, Cai S, Dong X. *Actinomyces ruminicola* sp. nov., isolated from cattle rumen. Int J Syst Evol Microbiol. 2006;9:2043-2048.

Baik KS, Choe HN, Park SC, Hwang YM, Kim EM, Park C, et al. *Sphingopyxis rigui* sp. nov. and *Sphingopyxis wooponensis* sp. nov., isolated from wetland freshwater, and emended description of the genus *Sphingopyxis*. Int J Syst Evol Microbiol. 2013;63:1297-1303.

Balderrama-Subieta A, Guzmán D, Minegishi H, Echigo A, Shimane Y, Hatada Y, et al. *Marinococcus tarijensis sp*. nov., a moderately halophilic bacterium isolated from a salt mine. Int J Syst Evol Microbiol. 2013;63:3319-3323.

Bergey DH, Harrison FC, Breed RS, Hammer BW, Huntoon FM. Bergey's Manual of Determinative Bacteriology, vol. The Williams & Wilkins Co, Baltimore; 1923.

Carreto L, Moore E, Nobre MF, Wait R, Riley PW, Sharp RJ, et al. *Rubrobacter xylanophilus* sp. nov., a new thermophilic species isolated from a thermally polluted effluent. Int J Syst Evol Microbiol. 1996;46:460-465.

Dastager SG, Qin L, Tang SK, Krishnamurthi S, Lee JC, Li WJ. *Arthrobacter enclensis* sp. nov., isolated from sediment sample. Arch Microbiol. 2014;196:775-782.

Evtushenko LI, Dorofeeva LV, Subbotin SA, Cole JR, Tiedje JM. *Leifsonia poae* gen. nov., sp. nov., isolated from nematode galls on Poa annua, and reclassification of *Corynebacterium aquaticum*’ Leifson 1962 as *Leifsonia aquatica* (ex Leifson 1962) gen. nov., nom. rev., comb. nov. and *Clavibacter xyli* Davis et al. 1984 with two subspecies as *Leifsonia xyli* (Davis et al. 1984) gen. nov., comb. nov. Int J Syst Evol Microbiol. 2000;50:371-380.

Ezaki T. Gallicola. Bergey's Manual of Systematics of Archaea and Bacteria. John Wiley & Sons, Ltd; 2015.

Faine S, Stallmn ND. Amended Descriptions of the Genus *Leptospira Noguchi* 1917 and the Species *L. interrogans* (Stimson 1907) Wenyon 1926 and *L. biflexa* (Wolbach and Binger 1914) Noguchi 1918. Int J Syst Evol Microbiol. 1982;32:461-463.

Funke G, Stubbs S, Altwegg M, Carlotti A, Collins MD. *Turicella otitidis* gen. nov., sp. nov., a coryneform bacterium isolated from patients with otitis media. Int J Syst Evol Microbiol. 1994;44:270-273.

Gebers R. Enrichment, Isolation, and Emended Description of *Pedomicrobium ferrugineum Aristovskaya* and *Pedomicrobium manganicum Aristovskaya*. Int J Syst Evol Microbiol.1981;31:302-316.

Gomila M, Tvrzova L, Teshim A, Sedláček I, Gonzalez-Escalona N, Zdráhal Z, et al. *Achromobacter marplatensis* sp. nov., isolated from a pentachlorophenol-contaminated soil. Int J Syst Evol Microbiol. 2011;61:2231-2237.

Gridneva E, Chernousova E, Dubinina G, Akimov V, Kuever J, Detkova E, et al. Taxonomic investigation of representatives of the genus *Sphaerotilus*: descriptions of *Sphaerotilus montanus* sp. nov., *Sphaerotilus hippei* sp. nov., *Sphaerotilus natans* subsp. natans subsp. nov. and *Sphaerotilus natans* subsp. *sulfidivorans* subsp. nov., and an emended description of the genus *Sphaerotilus*. Int J Syst Evol Microbiol. 2011;61:916-925.

Hong Z, Lai Q, Luo Q, Jiang S, Zhu R, Liang J, et al. *Sulfitobacter pseudonitzschiae* sp. nov., isolated from the toxic marine diatom Pseudo-nitzschia multiseries. Int J Syst Evol Microbiol. 2015;65:95-100.

Iguchi H, Yurimoto H, Sakai Y. *Methylovulum miyakonense* gen. nov., sp. nov., a type I methanotroph isolated from forest soil. Int J Syst Evol Microbiol. 2011;61:810-815.

Im WT, Liu QM, Yang JE, Kim MS, Kim SY, Lee ST, et al. *Panacagrimonas perspica* gen. nov., sp. nov., a novel member of Gammaproteobacteria isolated from soil of a ginseng field. J Microbiol. 2010;48:262-266.

Jordan EM, Thompson FL, Zhang X-H, Li Y, Vancanneyt M, Kroppenstedt RM et al. *Sneathiella chinensis* gen. nov., sp. nov., a novel marine alphaproteobacterium isolated from coastal sediment in Qingdao, China. Int J Syst Evol Microbiol. 2007;57:114-121.

Kämpfer P, Glaeser SP, Lodders N, Busse HJ, Falsen E. *Herminiimonas contaminans* sp. nov., isolated as a contaminant of biopharmaceuticals. Int J Syst Evol Microbiol. 2013;63:412-417.

Kanso S, Patel BKC. *Microvirga subterranea* gen. nov., sp. nov., a moderate thermophile from a deep subsurface Australian thermal aquifer. Int J Syst Evol Microbiol. 2003;53:401-406.

Kilpper-Bälz R, Schleifer K. Transfer of *Streptococcus morbillorum* to the genus *Gemella* as *Gemella morbillorum* comb. nov. Int J Syst Evol Microbiol. 1988;38:442-443.

Krishnamurthi S, Bhattacharya A, Schumann P, Dastager SG, Tang SK, Li WJ et al. *Microbacterium immunditiarum* sp. nov., an actinobacterium isolated from landfill surface soil, and emended description of the genus *Microbacterium*. Int J Syst Evol Microbiol. 2012;62:2187-2193.

Kudo Y, Oki K, Watanabe K. *Lactobacillus delbrueckii* subsp. sunkii subsp. nov., isolated from sunki, a traditional Japanese pickle. Int J Syst Evol Microbiol. 2012;62:2643-2649.

Kurahashi M, Fukunaga Y, Sakiyama Y, Harayama S, Yokota A. *Euzebya tangerina* gen. nov., sp. nov., a deeply branching marine actinobacterium isolated from the sea cucumber *Holothuria edulis*, and proposal of Euzebyaceae fam. nov., Euzebyales ord. nov. and Nitriliruptoridae subclassis nov. Int J Syst Evol Microbiol. 2010;60:2314-2319.

Leandro T, França L, Nobre MF, Schumann P, Rosselló-Móra R, da Costa MS. *Nevskia aquatilis* sp. nov. and *Nevskia persephonica* sp. nov., isolated from a mineral water aquifer and the emended description of the genus *Nevskia*. Syst Appl Microbiol. 2012;35: 297-301.

Li WJ, Zhang YQ, Park DJ, Li CT, Xu LH, Kim CJ, et al. *Duganella violaceinigra* sp. nov., a novel mesophilic bacterium isolated from forest soil. Int J Syst Evol Microbiol. 2004;54:1811-1814.

Liesack W, Bak F, Kreft JU, Stackebrandt E. *Holophaga foetida* gen. nov., sp. nov., a new, homoacetogenic bacterium degrading methoxylated aromatic compounds. Arch Microbiol. 1994;162:85-90.

Lin SY, Liu YC, Hameed A, Hsu YH, Lai WA, Shen FT, et al. *Azospirillum fermentarium* sp. nov., a nitrogen-fixing species isolated from a fermenter. Int J Syst Evol Microbiol. 2013;63:3762-3768.

Martínez-Hidalgo P, Ramírez-Bahena MH, Flores-Félix JD, Rivas R, Igual JM, Mateos PF, et al. Revision of the taxonomic status of type strains of *Mesorhizobium loti* and reclassification of strain USDA 3471T as the type strain of *Mesorhizobium erdmanii* sp. nov. and ATCC 33669T as the type strain of *Mesorhizobium jarvisii* sp. nov. Int J Syst Evol Microbiol. 2015;65:1703-1708.

Moreira D, Amils R. Phylogeny of *Thiobacillus cuprinus* and other mixotrophic *thiobacilli*: proposal for *Thiomonas* gen. nov. Int J Syst Evol Microbiol. 1997;47:522-528.

Murdoch DA, Shah HN. Reclassification of *Peptostreptococcus magnus* (Prevot 1933) Holdeman and Moore 1972 as *Finegoldia magna* comb. nov. and *Peptostreptococcus micros* (Prevot 1933) Smith 1957 as *Micromonas micros* comb. nov. Anaerobe. 1999;5:555-559.

Namsaraev Z, Akimov V, Tsapin A, Barinova E, Nealson K, Gorlenko V. *Marinospirillum celere* sp. nov., a novel haloalkaliphilic, helical bacterium isolated from Mono Lake. Int J Syst Evol Microbiol. 2009;59:2329-2332.

Ogg CD, Patel BK. *Sporolituus thermophilus* gen. nov., sp. nov., a citrate-fermenting thermophilic anaerobic bacterium from geothermal waters of the Great Artesian Basin of Australia. Int J Syst Evol Microbiol. 2009;59:2848-2853.

Patel NB, Tito RY, Obregon-Tito AJ, O'Neal L, Trujillo-Villaroel O, Marin-Reyes L, et al. *Peptoniphilus catoniae* sp. nov., isolated from a human faecal sample from a traditional Peruvian coastal community. Int J Syst Evol Microbiol. 2016;66:2019-2024.

Peng F, Liu M, Zhang L, Dai J, Luo X, An H, et al. *Planobacterium taklimakanense* gen. nov., sp. nov., a member of the family Flavobacteriaceae that exhibits swimming motility, isolated from desert soil. Int J Syst Evol Microbiol. 2009;59:1672-1678.

Pfennig N. *Rhodocyclus purpureus* gen. nov. and sp. nov., a Ring-Shaped, Vitamin B12-Requiring Member of the Family Rhodospirillaceae. Int J Syst Evol Microbiol. 1978;28:283-288.

Pikuta EV, Hoover RB, Bej AK, Marsic D, Detkova EN, Whitman WB, et al. *Tindallia californiensis* sp. nov., a new anaerobic, haloalkaliphilic, spore-forming acetogen isolated from Mono Lake in California. Extremophiles. 2003;7:327-334.

Rahul K, Sasikala C, Tushar L, Debadrita R, Ramana CV. *Alcanivorax xenomutans* sp. nov., a hydrocarbonoclastic bacterium isolated from a shrimp cultivation pond. Int J Syst Evol Microbiol. 2014;64:3553-3558.

Rees GN, Patel BK. *Desulforegula conservatrix* gen. nov., sp. nov., a long-chain fatty acid-oxidizing, sulfate-reducing bacterium isolated from sediments of a freshwater lake. Int J Syst Evol Microbiol. 2001;51:1911-1916.

Saha P, Krishnamurthi S, Mayilraj S, Prasad GS, Bora TC, Chakrabarti T. *Aquimonas voraii* gen. nov., sp. nov., a novel gammaproteobacterium isolated from a warm spring of Assam, India. Int J Syst Evol Microbiol. 2005;55:1491-1495.

Stackebrandt E, Verbarg S, Frühling A, Busse HJ, Tindall BJ. Dissection of the genus *Methylibium*: reclassification of *Methylibium fulvum* as *Rhizobacter fulvus* comb. nov., *Methylibium aquaticum* as *Piscinibacter aquaticus* gen. nov., comb. nov. and *Methylibium subsaxonicum* as *Rivibacter subsaxonicus* gen. nov., comb. nov. and emended descriptions of the genera *Rhizobacter* and *Methylibium*. Int J Syst Evol Microbiol. 2009;59:2552-2560.

Wang X, Wang Y, Yang X, Sun H, Li B, Zhang XH. *Photobacterium alginatilyticum* sp. nov., a marine bacterium isolated from bottom seawater. Int J Syst Evol Microbiol. 2017;67:1912-1917.

Willems A, Collins MD. Reclassification of *Oribaculum catoniae* (Moore and Moore 1994) as *Porphyromonas catoniae* comb. nov. and Emendation of the Genus *Porphyromonas*. Int J Syst Evol Microbiol. 1995;45:578-581.

Willems A, De Ley J, Gillis M, Kersters K. NOTES: Comamonadaceae, a New Family Encompassing the Acidovorans rRNA Complex, Including *Variovorax paradoxus* gen. nov., comb. nov., for *Alcaligenes paradoxus* (Davis 1969). Int J Syst Evol Microbiol. 1991;41:445-450.

Yabuuchi E, Kosako Y, Yano I, Hotta H, Nishiuchi Y. Transfer of Two *Burkholderia* and An *Alcaligenes* Species to *Ralstonia* Gen. Nov. Microbiol Immunol. 1995;39:897-904.

Yabuuchi E, Yano I, Oyaizu H, Hashimoto Y, Ezaki T, Yamamoto H. Proposals of *Sphingomonas paucimobilis* gen. nov. and comb. nov., *Sphingomonas parapaucimobilis* sp. nov., *Sphingomonas yanoikuyae* sp. nov., *Sphingomonas adhaesiva* sp. nov., *Sphingomonas capsulata* comb. nov., and two genospecies of the genus *Sphingomonas*. Microbiol Immunol. 1990;34:99-119.

Yakimov MM, Giuliano L, Chernikova TN, Gentile G, Abraham WR, Lünsdorf H, et al. *Alcalilimnicola halodurans* gen. nov., sp. nov., an alkaliphilic, moderately halophilic and extremely halotolerant bacterium, isolated from sediments of soda-depositing Lake Natron, East Africa Rift Valley. Int J Syst Evol Microbiol. 2001;51:2133-2143.

Yamamura H, Ashizawa H, Nakagawa Y, Hamada M, Ishida Y, Otoguro M, et al. *Actinomycetospora iriomotensis* sp. nov., a novel actinomycete isolated from a lichen sample. J Antibiot. 2011;64:289.

Yao S, Liu Y, Zhang M, Zhang X, Li H, Zhao T, et al. *Thermoactinomyces daqus* sp. nov., a thermophilic bacterium isolated from high-temperature Daqu. Int J Syst Evol Microbiol. 2014;64:206-210.

Yassin AF, Inglis TJJ, Hupfer H, Siering C, Schumann P, Busse HJ et al. *Cruoricaptor ignavus* gen. nov., sp. nov., a novel bacterium of the family Flavobacteriaceae isolated from blood culture of a man with bacteraemia. Syst Appl Microbiol. 2012;35:421-426.

**Table S9.** Highest temperature reported for the laboratory growth of the 47 validly published bacterial genera detected in HTP at 8:30 h of 20 October 2014.

| **Phyla** | **Genera** | **Upper limit of temperature (in °C) at which laboratory growth can occur** | **Reference** |
| --- | --- | --- | --- |
| *Actinobacteria* | *Actinomycetospora* | 30 | (Yamamura *et al.,* 2011) |
|  | *Brachybacterium* | 30 | (Collins *et al.,* 1988) |
|  | *Corynebacterium* | 42 | (Bernard *et al.,* 2016) |
|  | *Dietzia* | 30 | (Kim *et al.,* 2011) |
|  | *Leifsonia* | 28 | (Evtushenko *et al.,* 2000) |
|  | *Propionibacterium* | 45 | (Koussémon *et al.,* 2001) |
| *Aquificae* | *Hydrogenobacter* | 85 | (Takai *et al.,* 2001) |
| *Bacteroidetes* | *Planobacterium* | 37 | (Peng *et al.,* 2009) |
|  | *Sediminibacterium* | 42 | (Kang *et al.,* 2014) |
|  | *Sulfurihydrogenibium* | 80 | (O’Neill *et al.,* 2008) |
| *Chlorobi* | *Dehalogenimonas* | 42 | (Bowman *et al.,* 2013) |
| *Deinococcus-Thermus* | *Thermus* | 79 | (Brock and Freeze, 1969) |
| *Firmicutes* | *Anoxybacillus* | 70 | (Dulger *et al.,* 2004) |
|  | *Bacillus* | 70 | (Yang *et al.,* 2013) |
|  | *Brevibacillus* | 65 | (Inan *et al.,* 2012) |
|  | *Caldicellulosiruptor* | 90 | (Yang *et al.,* 2010) |
|  | *Dictyoglomus* | 80 | (Saiki *et al.,* 1985) |
|  | *Geobacillus* | 80 | (Coorevits *et al.,* 2012) |
|  | *Peptoniphilus* | 43 | (Patel *et al.,* 2016) |
|  | *Staphylococcus* | 40 | (Fuente *et al.,* 1985) |
| *Proteobacteria* | *Acinetobacter* | 44 | (Rooney *et al.,* 2016) |
|  | *Advenella* | 40 | (Shmareva *et al.,* 2016) |
|  | *Aquabacterium* | 40 | (Chen *et al.,* 2012) |
|  | *Azospirillum* | 37 | (Lin *et al.,* 2013) |
|  | *Beijerinckia* | 30 | (Oggerin *et al.,* 2009) |
|  | *Brevundimonas* | 42 | (Choi *et al.,* 2010) |
|  | *Burkholderia* | 37 | (Aizawa *et al.,* 2011) |
|  | *Buttiauxella* | 42 | (Muller *et al.,* 1996) |
|  | *Enhydrobacter* | 41 | (Staley *et al.,* 1987) |
|  | *Escherichia/Shigella* | 37 | (Liu *et al.,* 2015) |
|  | *Halomonas* | 50 | (Guan *et al.,* 2010) |
|  | *Marinobacter* | 50 | (Wang *et al.,* 2009) |
|  | *Methylobacterium* | 37 | (Wood *et al*., 1998) |
|  | *Methylophilus* | 37 | (Jenkins *et al.,* 1987) |
|  | *Paracoccus* | 45 | (Sun *et al.,* 2015) |
|  | *Providencia* | 45 | (Khunthongpan *et al.,* 2013) |
|  | *Pseudomonas* | 41 | (Wang and Sun, 2016) |
|  | *Psychrobacter* | 38 | (Maruyama *et al.,* 2000) |
|  | *Ralstonia* | 41 | (Yabuuchi *et al.,* 1995) |
|  | *Sphingomonas* | 40 | (Yabuuchi *et al.,* 1990) |
|  | *Tepidimonas* | 65 | (Moreira *et al.,* 2000) |
|  | *Thermomonas* | 50 | (Busse *et al.,* 2002) |
|  | *Thiobacillus* | 42 | (Brinkhoff *et al*., 1999) |
|  | *Thiofaba* | 51 | (Mori and Suzuki, 2008) |
|  | *Thiothrix* | 37 | (Howarth *et al.,* 1999) |
| *Spirochaetes* | *Spirochaeta* | 73 | (Aksenova *et al.,* 1992) |
| *Thermotogae* | *Fervidobacterium* | 90 | (Cai *et al.,* 2007) |

**Table S10.** Highest temperature reported for laboratory growth of the 88 bacterial genera that were detected in HTP at 14:30 h of 20 October 2014, and for which information is available regarding upper temperature-limit of laboratory growth.

| **Phyla** | **Genera** | **Upper limit of temperature (in °C) at which laboratory growth can occur** | **Reference** |
| --- | --- | --- | --- |
| *Acidobacteria* | *Geothrix* | 35 | (Coates *et al.,* 1999) |
| *Actinobacteria* | *Actinomyces* | 55 | (An *et al.,* 2006) |
|  | *Brachybacterium* | 30 | (Collins *et al.,* 1988) |
|  | *Brevibacterium* | 42 | (Kim *et al.,* 2013) |
|  | *Corynebacterium* | 42 | (Bernard *et al.,* 2016) |
|  | *Dietzia* | 30 | (Kim *et al.,* 2011) |
|  | *Marmoricola* | 40 | (Lee *et al.,* 2016) |
|  | *Microbacterium* | 42 | (Krishnamurthi *et al.,* 2012) |
|  | *Mycobacterium* | 43 | (Kusunoki and Ezaki, 1992) |
|  | *Nocardioides* | 37 | (Prauser, 1976) |
|  | *Patulibacter* | 39 | (Almeida *et al.,* 2013) |
|  | *Pedomicrobium* | 40 | (Gebers, 1981) |
|  | *Propionibacterium* | 45 | (Koussémon *et al.,* 2001) |
|  | *Rubrobacter* | 60 | (Carreto *et al.,* 1996) |
|  | *Turicella* | 37 | (Funke *et al.,* 1994) |
| *Aquificae* | *Hydrogenobacter* | 85 | (Takai *et al.,* 2001) |
| *Bacteroidetes* | *Chryseobacterium* | 37 | (Yang *et al.,* 2015) |
|  | *Cloacibacterium* | 40 | (Chun *et al.,* 2017) |
|  | *Cruoricaptor* | 37 | (Yassin *et al.,* 2012) |
|  | *Pedobacter* | 35 | (Zhang *et al.,* 2015) |
|  | *Sulfurihydrogenibium* | 80 | (O’Neill *et al.,* 2008) |
| *Chlorobi* | *Dehalogenimonas* | 42 | (Bowman *et al.,* 2013) |
| *Chloroflexi* | *Chloroflexus* | 59 | (Gaisin *et al.,* 2017) |
| *Deinococcus-Thermus* | *Deinococcus* | 45 | (Hussain *et al.,* 2016) |
|  | *Thermus* | 79 | (Brock and Freeze, 1969) |
|  | *Truepera* | 50 | (Albuquerque *et al.,* 2005) |
| *Firmicutes* | *Anoxybacillus* | 70 | (Dulger *et al.,* 2004) |
|  | *Bacillus* | 70 | (Yang *et al.,* 2013) |
|  | *Brevibacillus* | 65 | (Inan *et al.,* 2012) |
|  | *Caldicellulosiruptor* | 90 | ( Yang *et al.,* 2010) |
|  | *Dictyoglomus* | 80 | (Saiki *et al.,* 1985) |
|  | *Exiguobacterium* | 49 | (Crapart *et al.,* 2007) |
|  | *Finegoldia* | 37 | (Murdoch and Shah, 1999) |
|  | *Gemella* | 37 | (Kilpper-Bälz and Schleifer, 1988) |
|  | *Geobacillus* | 80 | (Coorevits *et al.,* 2012) |
|  | *Lysinibacillus* | 45 | (Ahmed *et al.,* 2007) |
|  | *Marinococcus* | 50 | (Balderrama-Subieta *et al.,* 2013) |
|  | *Paenibacillus* | 37 | (Shida *et al*., 1997) |
|  | *Peptoniphilus* | 43 | (Patel *et al.,* 2016) |
|  | *Planococcus* | 45 | (Gan *et al.,* 2018 ) |
|  | *Sporolituus* | 60 | (Ogg and Patel, 2009) |
|  | *Staphylococcus* | 40 | (Fuente *et al.,* 1985) |
|  | *Streptococcus* | 45 | (Sherman and Stark, 1931) |
|  | *Thermoactinomyces* | 60 | (Yao *et al.,* 2014) |
| *Nitrospirae* | *Thermodesulfovibrio* | 70 | (Sekiguchi *et al.,* 2008) |
| *Proteobacteria* | *Achromobacter* | 40 | (Gomila *et al.,* 2011) |
|  | *Acidovorax* | 37 | (Heylen *et al.,* 2008) |
|  | *Acinetobacter* | 44 | (Rooney *et al.,* 2016) |
|  | *Advenella* | 40 | (Shmareva *et al.,* 2016) |
|  | *Aeromonas* | 37 | (Galbis *et al.,* 2007) |
|  | *Alcanivorax* | 40 | (Rahul *et al.,* 2014) |
|  | *Beijerinckia* | 30 | (Oggerin *et al.,* 2009) |
|  | *Bradyrhizobium* | 37 | (Araújo *et al.,* 2017) |
|  | *Brevundimonas* | 42 | (Choi *et al.,* 2010) |
|  | *Burkholderia* | 37 | (Aizawa *et al.,* 2011) |
|  | *Comamonas* | 44 | (Chang *et al*., 2002) |
|  | *Delftia* | 45 | (Li *et al.,* 2015) |
|  | *Desulfovibrio* | 37 | (Maarel *et al.,* 1996) |
|  | *Enhydrobacter* | 41 | (Staley *et al.,* 1987) |
|  | *Escherichia/Shigella* | 37 | (Liu *et al.,* 2015) |
|  | *Halomonas* | 50 | (Guan *et al.,* 2010) |
|  | *Hyphomicrobium* | 30 | (McDonald *et al.,* 2001) |
|  | *Klebsiella* | 37 | (Bergey *et al.,* 1923) |
|  | *Litoreibacter* | 37 | (Romanenko *et al.,* 2011) |
|  | *Marinobacter* | 50 | (Wang *et al.,* 2009) |
|  | *Methylobacterium* | 37 | (Wood *et al*., 1998) |
|  | *Methyloversatilis* | 37 | (Kalyuzhnaya *et al.,* 2006) |
|  | *Nevskia* | 25 | (Leandro *et al.,* 2012) |
|  | *Paracoccus* | 45 | (Sun *et al.,* 2015) |
|  | *Pelomonas* | 40 | (Xie and Yokota, 2005) |
|  | *Photobacterium* | 50 | (Wang *et al.,* 2017) |
|  | *Pseudomonas* | 41 | (Wang and Sun, 2016) |
|  | *Ralstonia* | 41 | (Yabuuchi *et al.,* 1995) |
|  | *Rheinheimera* | 35 | (Ryu *et al.,* 2008) |
|  | *Rhodoplanes* | 43 | (Okamura *et al.,* 2009) |
|  | *Serratia* | 35 | (Geiger *et al.,* 2010) |
|  | *Sphingobium* | 37 | (Kumari *et al.,* 2009) |
|  | *Sphingomonas* | 40 | (Yabuuchi *et al.,* 1990) |
|  | *Stenotrophomonas* | 42 | (Lee *et al.,* 2011) |
|  | *Sulfitobacter* | 35 | (Hong *et al.,* 2015) |
|  | *Sulfurimonas* | 40 | (Inagaki *et al.,* 2003) |
|  | *Tepidimonas* | 60 | (Chen *et al.,* 2013) |
|  | *Thermomonas* | 50 | (Busse *et al.,* 2002) |
|  | *Thiofaba* | 51 | (Mori and Suzuki, 2008) |
|  | *Thiothrix* | 37 | (Howarth *et al.,* 1999) |
|  | *Variovorax* | 30 | (Willems *et al.,* 1991) |
| *Spirochaetes* | *Treponema* | 30 | (Evans *et al*., 2009) |
| *Thermotogae* | *Fervidobacterium* | 90 | (Cai *et al.,* 2007) |

**References used in Tables S9 and S10**

Ahmed I, Yokota A, Yamazoe A, Fujiwara T. Proposal of *Lysinibacillus boronitolerans* gen. nov. sp. nov., and transfer of *Bacillus fusiformis* to *Lysinibacillus fusiformis* comb. nov. and *Bacillus sphaericus* to *Lysinibacillus sphaericus* comb. nov. Int J Syst Evol Microbiol. 2007;57:1117-1125.

Aizawa T, Vijarnsorn P, Nakajima M, Sunairi M. *Burkholderia bannensis* sp. nov., an acid-neutralizing bacterium isolated from torpedo grass (*Panicum repens*) growing in highly acidic swamps. Int J Syst Evol Microbiol. 2011;61:1645-1650.

Aksenova HY, Rainey FA, Janssen PH, Zavarzin GA, Morgan HW. *Spirochaeta thermophila* sp. nov., an obligately anaerobic, polysaccharolytic, extremely thermophilic bacterium. Int J Syst Evol Microbiol. 1992;42:175-177.

Albuquerque L, Simoes C, Nobre MF, Pino NM, Battista JR, Silva MT, et al. *Truepera radiovictrix* gen. nov., sp. nov., a new radiation resistant species and the proposal of Trueperaceae fam. nov. FEMS Microbiol Lett. 2005;247:161-169.

Almeida B, Vaz-Moreira I, Schumann P, Nunes OC, Carvalho G, Crespo MTB. *Patulibacter medicamentivorans* sp. nov., isolated from activated sludge of a wastewater treatment plant. Int J Syst Evol Microbiol. 2013;63:2588-2593.

An D, Cai S, Dong X. *Actinomyces ruminicola* sp. nov., isolated from cattle rumen. Int J Syst Evol Microbiol. 2006;9:2043-2048.

Araújo J, Flores-Félix JD, Igual JM, Peix A, González-Andrés F, Díaz-Alcántara CA, et al. *Bradyrhizobium cajani* sp. nov. isolated from nodules of *Cajanus cajan*. Int J Syst Evol Microbiol. 2017;67:2236-2241.

Balderrama-Subieta A, Guzmán D, Minegishi H, Echigo A, Shimane Y, Hatada Y, et al. *Marinococcus tarijensis sp*. nov., a moderately halophilic bacterium isolated from a salt mine. Int J Syst Evol Microbiol. 2013;63:3319-3323.

Bergey DH, Harrison FC, Breed RS, Hammer BW, Huntoon FM. Bergey's Manual of Determinative Bacteriology, vol. The Williams & Wilkins Co, Baltimore; 1923.

Bernard KA, Pacheco AL, Loomer C, Burdz T, Wiebe D, Huynh C, et al. *Corynebacterium lowii* sp. nov. and *Corynebacterium oculi* sp. nov., derived from human clinical disease and an emended description of *Corynebacterium mastitidis*. Int J Syst Evol Microbiol. 2016;66:2803-2812.

Bowman KS, Nobre MF, da Costa MS, Rainey FA, Moe WM. *Dehalogenimonas alkenigignens* sp. nov., a chlorinated-alkane-dehalogenating bacterium isolated from groundwater. Int J Syst Evol Microbiol. 2013;63:1492-1498.

Brinkhoff T, Muyzer G, Wirsen CO, Kuever J. *Thiomicrospira chilensis* sp. nov., a mesophilic obligately chemolithoautotrophic sulfuroxidizing bacterium isolated from a Thioploca mat. Int J Syst Bacteriol. 1999;49:875-879.

Brock TD, Freeze H. *Thermus aquaticus* gen. n. and sp. n., a nonsporulating extreme thermophile. J. Bacteriol. 1969;98:289-297.

Busse H, Kämpfer P, Moore E, Nuutinen J, Tsitko I, Denner E, et al. *Thermomonas haemolytica* gen. nov., sp. nov., a gamma-proteobacterium from kaolin slurry. Int J Syst Evol Microbiol. 2002;52:473-483.

Cai J, Wang Y, Liu D, Zeng Y, Xue Y, Ma Y, et al. *Fervidobacterium changbaicum* sp. nov., a novel thermophilic anaerobic bacterium isolated from a hot spring of the Changbai Mountains, China. Int J Syst Evol Microbiol. 2007;57:2333-2336.

Carreto L, Moore E, Nobre MF, Wait R, Riley PW, Sharp RJ, et al. *Rubrobacter xylanophilus* sp. nov., a new thermophilic species isolated from a thermally polluted effluent. Int J Syst Evol Microbiol.1996;46:460-465.

Chang YH, Han JI, Chun J, Lee KC, Rhee MS, Kim YB, et al. *Comamonas koreensis* sp. nov., a non-motile species from wetland in Woopo, Korea. Int J Syst Evol Microbiol. 2002;52:377-381.

Chen WM, Cho NT, Yang SH, Arun A, Young CC, Sheu SY. *Aquabacterium limnoticum* sp. nov., isolated from a freshwater spring. Int J Syst Evol Microbiol. 2012;62:698-704.

Chen WM, Huang HW, Chang JS, Han YL, Guo TR, Sheu SY. *Tepidimonas fonticaldi* sp. nov., a slightly thermophilic betaproteobacterium isolated from a hot spring. Int J Syst Evol Microbiol. 2013;63:1810-1816.

Choi JH, Kim MS, Roh SW, Bae JW. *Brevundimonas basaltis* sp. nov., isolated from black sand. Int J Syst Evol Microbiol. 2010;60:1488-1492.

Chun BH, Lee Y, Jin HM, Jeon CO. *Cloacibacterium caeni* sp. nov., isolated from activated sludgeInt J Syst Evol Microbiol. 2017;67:1688-1692.

Coates JD, Ellis DJ, Gaw CV, Lovley DR. *Geothrix fermentans* gen. nov., sp. nov., a novel Fe (III)-reducing bacterium from a hydrocarbon-contaminated aquifer. Int J Syst Evol Microbiol. 1999;49:1615-1622.

Collin MD, Brown J, Jones D. *Brachybacterium faecium* gen. nov., sp. nov., a coryneform bacterium from poultry deep litter. Int J Syst Evol Microbiol. 1988;38:45-48.

Coorevits A, Dinsdale AE, Halket G, Lebbe L, De Vos P, Van Landschoot A, et al. Taxonomic revision of the genus *Geobacillus*: emendation of *Geobacillus*, *G. stearothermophilus*, *G. jurassicus*, *G. toebii*, *G. thermodenitrificans* and *G. thermoglucosidans* (nom. corrig., formerly ‘thermoglucosidasius’); transfer of *Bacillus thermantarcticus* to the genus as *G. thermantarcticus* comb. nov.; proposal of *Caldibacillus debilis* gen. nov., comb. nov.; transfer of *G. tepidamans* to *Anoxybacillus* as *A. tepidamans* comb. nov.; and proposal of *Anoxybacillus caldiproteolyticus* sp. nov. Int J Syst Evol Microbiol. 2012;62:1470-1485.

Crapart S, Fardeau ML, Cayol JL, Thomas P, Sery C, Ollivier B, et al. *Exiguobacterium profundum* sp. nov., a moderately thermophilic, lactic acid-producing bacterium isolated from a deep-sea hydrothermal vent. Int J Syst Evol Microbiol. 2007;57:287-292.

De la Fuente R, Suarez G, Schleifer K. *Staphylococcus aureus* subsp. anaerobius subsp. nov., the causal agent of abscess disease of sheep. Int J Syst Evol Microbiol. 1985;35:99-102.

Dulger S, Demirbag Z, Belduz AO. *Anoxybacillus ayderensis* sp. nov. and *Anoxybacillus kestanbolensis* sp. Nov. Int J Syst Evol Microbiol. 2004;54:1499-1503.

Evans NJ, Brown JM, Demirkan I, Murray RD, Birtles RJ, Hart CA, et al. *Treponema pedis* sp. nov., a spirochaete isolated from bovine digital dermatitis lesions. Int J Syst Evol Microbiol. 2009;59:987-991.

Evtushenko LI, Dorofeeva LV, Subbotin SA, Cole JR, Tiedje JM. *Leifsonia poae* gen. nov., sp. nov., isolated from nematode galls on Poa annua, and reclassification of *Corynebacterium aquaticum*’ Leifson 1962 as *Leifsonia aquatica* (ex Leifson 1962) gen. nov., nom. rev., comb. nov. and *Clavibacter xyli* Davis et al. 1984 with two subspecies as *Leifsonia xyli* (Davis et al. 1984) gen. nov., comb. nov. Int J Syst Evol Microbiol. 2000;50:371-380.

Funke G, Stubbs S, Altwegg M, Carlotti A, Collins MD. *Turicella otitidis* gen. nov., sp. nov., a coryneform bacterium isolated from patients with otitis media. Int J Syst Evol Microbiol. 1994;44:270-273.

Gaisin VA, Kalashnikov AM, Grouzdev DS, Sukhacheva MV, Kuznetsov BB, Gorlenko VM. *Chloroflexus islandicus* sp. nov., a thermophilic filamentous anoxygenic phototrophic bacterium from a geyser. Int J Syst Evol Microbiol. 2017;67:1381-1386.

Gan L, Zhang H, Tian J, Li X, Long X, Zhang Y, et al. *Planococcus salinus* sp. nov., a moderately halophilic bacterium isolated from a saline-alkali soil. Int J Syst Evol Microbiol. 2018;68:589-595.

Gebers R. Enrichment, Isolation, and Emended Description of *Pedomicrobium ferrugineum Aristovskaya* and *Pedomicrobium manganicum Aristovskaya*. Int J Syst Evol Microbiol.1981;31:302-316.

Geiger A, Fardeau ML, Falsen E, Ollivier B, Cuny G. *Serratia glossinae* sp. nov., isolated from the midgut of the tsetse fly *Glossina palpalis gambiensis*. Int J Syst Evol Microbiol. 2010;60:1261-1265.

Gomila M, Tvrzova L, Teshim A, Sedláček I, Gonzalez-Escalona N, Zdráhal Z, et al. *Achromobacter marplatensis* sp. nov., isolated from a pentachlorophenol-contaminated soil. Int J Syst Evol Microbiol. 2011;61:2231-2237.

Guan TW, Xiao J, Zhao K, Luo XX, Zhang XP, Zhang LL. *Halomonas xinjiangensis* sp. nov., a halotolerant bacterium isolated from a salt lake. Int J Syst Evol Microbiol. 2010;60:349-352.

Heylen K, Lebbe L, De Vos P*. Acidovorax caeni* sp. nov., a denitrifying species with genetically diverse isolates from activated sludge. Int J Syst Evol Microbiol. 2008;58:73-77.

Hong Z, Lai Q, Luo Q, Jiang S, Zhu R, Liang J, et al . *Sulfitobacter pseudonitzschiae* sp. nov., isolated from the toxic marine diatom Pseudo-nitzschia multiseries. Int J Syst Evol Microbiol. 2015;65:95-100.

Howarth R, Unz RF, Seviour EM, Seviour RJ, Blackall LL, Pickup RW, et al. Phylogenetic relationships of filamentous sulfur bacteria (*Thiothrix spp*. and *Eikelboom* type 021N bacteria) isolated from waste water treatment plants and description of *Thiothrix eikelboomii* sp. nov., *Thiothrix unzii* sp. nov., *Thiothrix fructosivorans* sp. nov. and *Thiothrix defluvii* sp. nov. Int J Syst Evol Microbiol. 1999;49:1817-1827.

Hussain F, Khan IU, Habib N, Xian WD, Hozzein WN, Zhang ZD, et al. *Deinococcus saudiensis* sp. nov., isolated from desert. Int J Syst Evol Microbiol. 2016;66:5106-5111.

Inagaki F, Takai K, Kobayashi H, Nealson KH, Horikoshi K. *Sulfurimonas autotrophica* gen. nov., sp. nov., a novel sulfur-oxidizing ε-proteobacterium isolated from hydrothermal sediments in the Mid-Okinawa Trough. Int J Syst Evol Microbiol. 2003;53:1801-1805.

Inan K, Canakci S, Belduz AO, Sahin F. *Brevibacillus aydinogluensis* sp. nov., a moderately thermophilic bacterium isolated from Karakoc hot spring. Int J Syst Evol Microbiol. 2012;62:849-855.

Jenkins O, Byrom D, JonesD. *Methylophilus*: a new genus of methanol-utilizing bacteria. Int J Syst Evol Microbiol. 1987;37:446-448.

Kalyuzhnaya MG, De Marco P, Bowerman S, Pacheco CC, Lara JC, Lidstrom ME, et al. *Methyloversatilis universalis* gen. nov., sp. nov., a novel taxon within the Betaproteobacteria represented by three methylotrophic isolates. Int J Syst Evol Microbiol. 2006;56:2517-2522.

Kang H, Kim H, Lee BI, Joung Y, Joh K. *Sediminibacterium goheungense* sp. nov., isolated from a freshwater reservoir. Int J Syst Evol Microbiol. 2014;64:1328-1333.

Khunthongpan S, Sumpavapol P, Tanasupawat S. *Providencia thailandensis* sp. nov., isolated from seafood processing wastewater. J Gen Appl Microbiol. 2013;59:185-190.

Kilpper-Bälz R, Schleifer K. Transfer of *Streptococcus morbillorum* to the genus *Gemella* as *Gemella morbillorum* comb. nov. Int J Syst Evol Microbiol. 1988;38:442-443.

Kim J, Roh SW, Choi JH, Jung MJ, Nam YD, Kim MS, et al. *Dietzia alimentaria* sp. nov., isolated from a traditional Korean food. Int J Syst Evol Microbiol. 2011;61:2254-2258.

Kim J, Srinivasan S, You T, Bang JJ, Park S, Lee SS. *Brevibacterium ammoniilyticum* sp. nov., an ammonia-degrading bacterium isolated from sludge of a wastewater treatment plant. Int J Syst Evol Microbiol. 2013;63:1111-1118.

Koussémon M, Combet-Blanc Y, Patel B, Cayol JL, Thomas P, Garcia JL, et al. *Propionibacterium microaerophilum* sp. nov., a microaerophilic bacterium isolated from olive mill wastewater. Int J Syst Evol Microbiol. 2001;51:1373-1382.

Krishnamurthi S, Bhattacharya A, Schumann P, Dastager SG, Tang SK, Li WJ . *Microbacterium immunditiarum* sp. nov., an actinobacterium isolated from landfill surface soil, and emended description of the genus *Microbacterium*. Int J Syst Evol Microbiol. 2012;62:2187-2193.

Kumari H, Gupta SK, Jindal S, Katoch P, Lal R. *Sphingobium lactosutens* sp. nov., isolated from a hexachlorocyclohexane dump site and *Sphingobium abikonense* sp. nov., isolated from oil-contaminated soil. Int J Syst Evol Microbiol. 2009;59:2291-2296.

Kusunoki S, Ezaki T. Proposal of *Mycobacterium peregrinum* sp. nov., nom. rev., and elevation of *Mycobacterium chelonae* subsp. abscessus (Kubica et al.) to species status: *Mycobacterium abscessus* comb. nov. Int J Syst Evol Microbiol. 1992;42:240-245.

Leandro T, França L, Nobre MF, Schumann P, Rosselló-Móra R, da Costa MS . *Nevskia aquatilis* sp. nov. and *Nevskia persephonica* sp. nov., isolated from a mineral water aquifer and the emended description of the genus *Nevskia*. Syst Appl Microbiol. 2012;35:297-301.

Lee HY, Liu Q, Kang MS, Kim SK, Lee SY, Im WT. *Marmoricola ginsengisoli* sp. nov. and *Marmoricola pocheonensis* sp. nov. isolated from a ginseng-cultivating field. Int J Syst Evol Microbiol. 2016;66:1996-2001.

Lee M, Woo SG, Chae M, Shin MC, Jung HM, Ten LN. *Stenotrophomonas daejeonensis* sp. nov., isolated from sewage. Int J Syst Evol Microbiol. 2011;61:598-604.

Li CT, Yan ZF, Chu X, Hussain F, Xian WD, Yunus Z, et al. *Delftia deserti* sp. nov., isolated from a desert soil sample. Antonie van Leeuwenhoek. 2015;107:1445-1450.

Lin SY, Liu YC, Hameed A, Hsu YH, Lai WA, Shen FT, et al. Azospirillum fermentarium sp. nov., a nitrogen-fixing species isolated from a fermenter. Int J Syst Evol Microbiol. 2013;63:3762-3768.

Liu S, Jin D, Lan R, Wang Y, Meng Q, Dai H, et al. *Escherichia marmotae* sp. nov., isolated from faeces of Marmota himalayana. Int J Syst Evol Microbiol. 2015;65:2130-2134.

Maruyama A, Honda D, Yamamoto H, Kitamura K, Higashihara T. Phylogenetic analysis of psychrophilic bacteria isolated from the Japan Trench, including a description of the deep-sea species *Psychrobacter pacificensis* sp. nov. Int J Syst Evol Microbiol. 2000;50:835-846.

McDonald IR, Doronina NV, Trotsenko YA, McAnulla C, Murrell JC. *Hyphomicrobium chloromethanicum* sp. nov. and *Methylobacterium chloromethanicum* sp. nov., chloromethane-utilizing bacteria isolated from a polluted environment. Int J Syst Evol Microbiol. 2001;51:119-122.

Minana-Galbis D, Farfan M, Fusté MC, Lorén JG. *Aeromonas bivalvium* sp. nov., isolated from bivalve molluscs. Int J Syst Evol Microbiol. 2007;57:582-587.

Moreira C, Rainey FA, Nobre MF, da Silva MT, da Costa MS. *Tepidimonas ignava* gen. nov., sp. nov., a new chemolithoheterotrophic and slightly thermophilic member of the beta-Proteobacteria. Int J Syst Evol Microbiol. 2000;50:735-742.

Mori K, Suzuki KI. *Thiofaba tepidiphila* gen. nov., sp. nov., a novel obligately chemolithoautotrophic, sulfur-oxidizing bacterium of the Gammaproteobacteria isolated from a hot spring. Int J Syst Evol Microbiol. 2008;58:1885-1891.

Müller HE, Brenner DJ, Fanning GR, Grimont PA, Kämpfer P. Emended description of *Buttiauxella agrestis* with recognition of six new species of *Buttiauxella* and two new species of *Kluyvera*: *Buttiauxella ferragutiae* sp. nov., *Buttiauxella gaviniae* sp. nov., *Buttiauxella brennerae* sp. nov., *Buttiauxella izardii* sp. nov., *Buttiauxella noackiae* sp. nov., *Buttiauxella warmboldiae* sp. nov., *Kluyvera cochleae* sp. nov., and *Kluyvera georgiana* sp. nov. Int J Syst Evol Microbiol. 1996;46:50-63.

Murdoch DA, Shah HN. Reclassification of *Peptostreptococcus magnus* (Prevot 1933) Holdeman and Moore 1972 as *Finegoldia magna* comb. nov. and *Peptostreptococcus micros* (Prevot 1933) Smith 1957 as *Micromonas micros* comb. nov. Anaerobe. 1999;5:555-559.

Ogg CD, Patel BK. *Sporolituus thermophilus* gen. nov., sp. nov., a citrate-fermenting thermophilic anaerobic bacterium from geothermal waters of the Great Artesian Basin of Australia. Int J Syst Evol Microbiol. 2009;59:2848-2853.

Oggerin M, Arahal DR, Rubio V, Marín I. Identification of *Beijerinckia fluminensis* strains CIP 106281T and UQM 1685T as *Rhizobium radiobacter* strains, and proposal of *Beijerinckia doebereinerae* sp. nov. to accommodate *Beijerinckia fluminensis* LMG 2819. Int J Syst Evol Microbiol. 2009;59:2323-2328.

Okamura K, Kanbe T, Hiraishi A. *Rhodoplanes serenus* sp. nov., a purple non-sulfur bacterium isolated from pond water. Int J Syst Evol Microbiol. 2009;59:531-535.

O'Neill AH, Liu Y, Ferrera I, Beveridge TJ, Reysenbach AL. *Sulfurihydrogenibium rodmanii* sp. nov., a sulfur-oxidizing chemolithoautotroph from the Uzon Caldera, Kamchatka Peninsula, Russia, and emended description of the genus *Sulfurihydrogenibium*. Int J Syst Evol Microbiol. 2008;58:1147-1152.

Patel NB, Tito RY, Obregon-Tito AJ, O'Neal L, Trujillo-Villaroel O, Marin-Reyes L, et al. *Peptoniphilus catoniae* sp. nov., isolated from a human faecal sample from a traditional Peruvian coastal community. Int J Syst Evol Microbiol. 2016;66:2019-2024.

Peng F, Liu M, Zhang L, Dai J, Luo X, An H, et al . *Planobacterium taklimakanense* gen. nov., sp. nov., a member of the family Flavobacteriaceae that exhibits swimming motility, isolated from desert soil. Int J Syst Evol Microbiol. 2009;59:1672-1678.

Prauser H. *Nocardioides*, a new genus of the order Actinomycetales. Int J Syst Evol Microbiol. 1976;26:58-65.

Rahul K, Sasikala C, Tushar L, Debadrita R, Ramana CV. *Alcanivorax xenomutans* sp. nov., a hydrocarbonoclastic bacterium isolated from a shrimp cultivation pond. Int J Syst Evol Microbiol. 2014;64:3553-3558.

Romanenko LA, Tanaka N, Frolova GM, Svetashev VI, Mikhailov VV. *Litoreibacter albidus* gen. nov., sp. nov. and *Litoreibacter janthinus* sp. nov., members of the class Alphaproteobacteria isolated from the seashore. Int J Syst Evol Microbiol. 2011;61:148-154.

Rooney AP, Dunlap CA, Flor-Weiler LB. *Acinetobacter lactucae* sp. nov., isolated from iceberg lettuce (Asteraceae: *Lactuca sativa*). Int J Syst Evol Microbiol. 2016;66:3566-3572.

Ryu SH, Chung BS, Park M, Lee SS, Lee SS, Jeon CO. *Rheinheimera soli* sp. nov., a gammaproteobacterium isolated from soil in Korea. Int J Syst Evol Microbiol. 2008;58:2271-2274.

Saiki T, Kobayashi Y, Kawagoe K, Beppu T. *Dictyoglomus thermophilum* gen. nov., sp. nov., a chemoorganotrophic, anaerobic, thermophilic bacterium. Int J Syst Evol Microbiol. 1985;35:253-259.

Sekiguch Y, Muramatsu M, Imachi H, Narihiro T, Ohashi A, Harada H, et al. *Thermodesulfovibrio aggregans* sp. nov. and *Thermodesulfovibrio thiophilus* sp. nov., anaerobic, thermophilic, sulfate-reducing bacteria isolated from thermophilic methanogenic sludge, and emended description of the genus *Thermodesulfovibrio*. Int J Syst Evol Microbiol. 2008;58:2541-2548.

Sherman JM, Stark P. *Streptococci* which grow at high temperatures. J Bacteriol. 1931;*22*:275-285.

Shida O, Takagi H, Kadowaki K, Nakamura LK, Komagata K. Transfer of *Bacillus alginolyticus*, *Bacillus chondroitinus*, *Bacillus curdlanolyticus*, *Bacillus glucanolyticus*, *Bacillus kobensis*, and *Bacillus thiaminolyticus* to the genus *Paenibacillus* and emended description of the genus *Paenibacillus*. Int J Syst Bacteriol. 1997;47:289-98.

Shmareva M, Agafonova N, Kaparullina E, Doronina N, Trotsenko YA. Emended Descriptions of *Advenella kashmirensis* subsp. kashmirensis subsp. nov., *Advenella kashmirensis* subsp. methylica subsp. nov., *and Methylopila turkiensis* sp. nov. Microbiology. 2016;85:646-648.

Staley JT, Irgens RL, Brenner DJ. *Enhydrobacter aerosaccus* gen. nov., sp. nov., a gas-vacuolated, facultatively anaerobic, heterotrophic rod. Int J Syst Evol Microbiol. 1987;37:289-291.

Sun X, Luo P, Li M. *Paracoccus angustae* sp. nov., isolated from soil. Int J Syst Evol Microbiol. 2015;65:3469-3475.

Takai K, Komatsu T, Horikoshi K. *Hydrogenobacter subterraneus* sp. nov., an extremely thermophilic, heterotrophic bacterium unable to grow on hydrogen gas, from deep subsurface geothermal water. Int J Syst Evol Microbiol. 2001;51:1425-1435.

Van der Maarel MJ, van Bergeijk S, van Werkhoven AF, Laverman AM, Meijer WG, Stam WT, et al. Cleavage of dimethylsulfoniopropionate and reduction of acrylate by *Desulfovibrio acrylicus* sp. nov. Arch Microbiol. 1996;166:109-115.

Wang CY, Ng CC, Tzeng WS, Shyu YT. *Marinobacter szutsaonensis* sp. nov., isolated from a solar saltern. Int J Syst Evol Microbiol. 2009;59:2605-2609.

Wang M, Sun L. *Pseudomonas oceani* sp. nov., isolated from deep seawater. Int J Syst Evol Microbiol. 2016;66:4250-4255.

Wang X, Wang Y, Yang X, Sun H, Li B, Zhang XH. *Photobacterium alginatilyticum* sp. nov., a marine bacterium isolated from bottom seawater. Int J Syst Evol Microbiol. 2017;67:1912-1917.

Willems A, De Ley J, Gillis M, Kersters K. NOTES: Comamonadaceae, a New Family Encompassing the *Acidovorans* rRNA Complex, Including *Variovorax paradoxus* gen. nov., comb. nov., for *Alcaligenes paradoxus* (Davis 1969). Int J Syst Evol Microbiol. 1991;41:445-450.

Wood AP, Kelly DP, McDonald IR, Jordan SL, Morgan TD, Khan S, et al. A novel pink-pigmented facultative methylotroph, *Methylobacterium thiocyanatum* sp. nov., capable of growth on thiocyanate or cyanate as sole nitrogen sources. Arch Microbiol.1998;169:148-158.

Xie CH, Yokota A. Reclassification of *Alcaligenes latus* strains IAM 12599T and IAM 12664 and *Pseudomonas saccharophila* as *Azohydromonas lata* gen. nov., comb. nov., *Azohydromonas australica* sp. nov. and *Pelomonas saccharophila* gen. nov., comb. nov., respectively. Int J Syst Evol Microbiol. 2005;55:2419-2425.

Yabuuchi E, Kosako Y, Yano I, Hotta H, Nishiuchi Y. Transfer of Two *Burkholderia* and An *Alcaligenes* Species to *Ralstonia* Gen. Nov. Microbiol Immunol. 1995;39:897-904.

Yabuuchi E, Yano I, Oyaizu H, Hashimoto Y, Ezaki T, Yamamoto H. Proposals of *Sphingomonas paucimobilis* gen. nov. and comb. nov., *Sphingomonas parapaucimobilis* sp. nov., *Sphingomonas yanoikuyae* sp. nov., *Sphingomonas adhaesiva* sp. nov., *Sphingomonas capsulata* comb. nov., and two genospecies of the genus *Sphingomonas*. Microbiol Immunol. 1990;34:99-119.

Yamamura H, Ashizawa H, Nakagawa Y, Hamada M, Ishida Y, Otoguro M, et al. *Actinomycetospora iriomotensis* sp. nov., a novel actinomycete isolated from a lichen sample. J Antibiot. 2011;64:289.

Yang F, Liu Hm, Zhang R, Chen Db, Wang X, Li Sp, et al. *Chryseobacterium shandongense* sp. nov., isolated from soil. Int J Syst Evol Microbiol. 2015;65:1860-1865.

Yang G, Chen M, Yu Z, Lu Q, Zhou S. *Bacillus composti sp*. nov. and *Bacillus thermophilus sp*. nov., two thermophilic, Fe (III)-reducing bacteria isolated from compost. Int J Syst Evol Microbiol. 2013;63:3030-3036.

Yang SJ, Kataeva I, Wiegel J, Yin Y, Dam P, Xu Y, et al. Classification of ‘*Anaerocellum thermophilum*’ strain DSM 6725 as *Caldicellulosiruptor bescii* sp. nov. Int J Syst Evol Microbiol. 2010;60,2011-2015.

Yao S, Liu Y, Zhang M, Zhang X, Li H, Zhao T, et al. *Thermoactinomyces daqus* sp. nov., a thermophilic bacterium isolated from high-temperature Daqu. Int J Syst Evol Microbiol. 2014;64:206-210.

Yassin AF, Inglis TJJ, Hupfer H, Siering C, Schumann P, Busse HJ, et al. *Cruoricaptor ignavus* gen. nov., sp. nov., a novel bacterium of the family Flavobacteriaceae isolated from blood culture of a man with bacteraemia. Syst Appl Microbiol. 2012;35:421-426.

Zhang H, Zhang J, Song M, Cheng Mg, Wu Yd, Guo Sh, et al. *Pedobacter nanyangensis* sp. nov., isolated from herbicide-contaminated soil. Int J Syst Evol Microbiol. 2015;65:3517-3521.

**Table S11.** Summary of the PCR-amplified 16S rRNA gene cloning and sequencing-based analysis of the aquatic community present at the HTP site in August 2009.

| **Phylum affiliation of the clones** | **Serial no. Name of the representative 16S rRNA gene clones constructed from total environmental DNA**  **(GenBank/EMBL accession number of the sequence)**  **# Number of sibling clones present in the library for that representative** | **Type strain(s) with which maximum sequence similarity was recorded** | **Identity (Query coverage)** | **Upper limit of temperature for growth *in vitro*** | **Reference** |
| --- | --- | --- | --- | --- | --- |
| *Bacteroidetes* | 1. 2010V2W clone B (FR848401) # 3 | *Lentimicrobium saccharophilum* TBC1^T^ | 89 % (99 %) | 40 | (Sun *et al*., 2016) |
|  | 2. 2010V2W clone C (FR848402) # 4 |  | 89 % (97 %) |  |  |
| *Actinobacteria* | 3. clone 2b21 (FN594534) # 2 | **Microbacterium schleiferi* DSM 20489^T^ | 93 % (98 %) | 42 | (Krishnamurthi *et al.,* 2012) |
| *Alphaproteobacteria* | 4. 2010V2W clone A (FR848400) # 4 | *Sandaracinobacter sibiricus* RB16-17^T^ | 92 % (98 %) | 30 | (Yurkov *et al*., 1997) |
|  |  | **Pedomicrobium* *manganicum* ATCC 33121^T^ | 93 % (96 %) | 40 | (Gebers, 1981) |
|  | 5. clone TWC 222 (HE774688) # 1 | *Rhodobacter capsulatus* ATCC 11166^T^  *Rhodobacter sediminis* N1^T^ | 99 % (100 %) | 45 | (Subhash *et al*., 2016) |
| *Betaproteobacteria* | 6. clone Puga_TWC_XXX383 (HE798186) # 3 | *Limnobacter thiooxidans* CS-K2^T^ | 99 % (100 %) | 50 | (Nguyen *et al.*, 2017) |
|  | 7. clone TWC 8 (HE774686) # 1 | *Hydrogenophaga laconesensis* HWB-10^T^ | 96 % (39 %) | 37 | (Yoon *et al*., 2008) |
|  | 8. clone TWC 14 (HE774687) # 1 | *Hydrogenophaga laconesensis* HWB-10^T^ | 96 % (40 %) |  |  |
|  | 9. Puga_TWC_XXX281 (HE798187) # 2 | ***Herbaspirillum canariense* SUEMI03^T^ | 97 % (92 %) | 35 | (Baldani *et al*., 1986) |
|  |  | *Oxalicibacterium solurbis* NBRC 102665^T^ |  | 37 | (Sahin *et al*., 2010) |
| *Gammaproteobacteria* | 10. clone 2b27 (FN594535) # 4 | ***Lysobacter enzymogenes* DSM 2043^T^ | 97 % (97 %) | 40 | (Christensen *et al*., 1978) |
|  | 11. clone 2b28 (FN594536) # 6 | ***Lysobacter* enzymogenes DSM 2043^T^ | 99 % (98 %) |  |  |
|  | 12. clone 1b3 (FN594537) # 5 | ***Lysobacter enzymogenes* DSM 2043^T^ | 99 % (99 %) |  |  |
|  | 13. clone 1a1 (FN594538) # 4 | ***Lysobacter enzymogenes* DSM 2043^T^ | 99 % (97 %) |  |  |
|  | 14. clone 1b16 (FN594539) # 3 | ***Lysobacter enzymogenes* DSM 2043^T^ | 96 % (98 %) |  |  |
|  | 15. Puga_TWC_XXX711 (HE798188) # 5 | **Acinetobacter junii* ATCC 17908^T^ | 100 % (100 %) | 44 | (Rooney *et al.,* 2016) |
|  | 16. clone TWC 7 (HE774684) # 1 | **Pseudomonas putida* NBRC 14164^T^ | 100 % (33 %) | 41 | (Wang and Sun, 2016) |
|  | 17. clone TWC 15 (HE774685) # 1 | *Samsonia erythrinae* CFBP 5236^T^ | 100 % (26 %) | 39 | (Sutra *et al*., 2001) |

*	These genera were also detected in 2014.

**	These genera were also detected in 2013.

**References used in Table S11**

Baldani JI, Baldani VLD, Seldin L, Döbereiner J. Characterization of *Herbaspirillum seropedicae* gen. nov., sp. nov., a Root-Associated Nitrogen-Fixing Bacterium. Int J Syst Evol Microbiol. 1986;36:86-93.

Christensen P, Cook FD. *Lysobacter*, a New Genus of Nonfruiting, Gliding Bacteria with a High Base Ratio. Int J Syst Evol Microbiol. 1978;28:367-93.

Gebers R. Enrichment, Isolation, and Emended Description of *Pedomicrobium ferrugineum Aristovskaya* and *Pedomicrobium manganicum Aristovskaya*. Int J Syst Evol Microbiol.1981;31:302-316.

Krishnamurthi S, Bhattacharya A, Schumann P, Dastager SG, Tang SK, Li WJ . *Microbacterium immunditiarum* sp. nov., an actinobacterium isolated from land fill surface soil, and emended description of the genus *Microbacterium*. Int J Syst Evol Microbiol. 2012;62:2187-2193.

Nguyen TM, Kim J. *Limnobacter humi* sp. nov., a thiosulfate-oxidizing, heterotrophic bacterium isolated from humus soil, and emended description of the genus *Limnobacter* Spring et al. 2001. J Microbiol. 2017;55:508-13.

Rooney AP, Dunlap CA, Flor-Weiler LB. *Acinetobacter lactucae* sp. nov., isolated from iceberg lettuce (Asteraceae: Lactuca sativa). Int J Syst Evol Microbiol. 2016;66:3566-3572.

Sahin N, Gonzalez JM, Iizuka T, Hill JE. Characterization of two aerobic ultramicrobacteria isolated from urban soil and a description of *Oxalicibacterium solurbis* sp. nov. FEMS Microbiol Lett. 2010;307:25-9.

Subhash Y, Lee SS. *Rhodobacter sediminis* sp. nov., isolated from lagoon sediments. Int J Syst Evol Microbiol. 2016;66:2965-70.

Sun L, Toyonaga M, Ohashi A, Tourlousse DM, Matsuura N, Meng XY, et al. *Lentimicrobium saccharophilum* gen. nov., sp. nov., a strictly anaerobic bacterium representing a new family in the phylum Bacteroidetes, and proposal of Lentimicrobiaceae fam. nov. Int J Syst Evol Microbiol. 2016;66:2635-42.

Sutra L, Christen R, Bollet C, Simoneau P, Gardan *L. Samsonia erythrinae* gen. nov., sp. nov., isolated from bark necrotic lesions of *Erythrina sp*., and discrimination of plant-pathogenic Enterobacteriaceae by phenotypic features. Int J Syst Evol Microbiol. 2001;51:1291-304.

Wang M, Sun L. *Pseudomonas oceani* sp. nov., isolated from deep seawater. Int J Syst Evol Microbiol. 2016;66:4250-4255.

Yoon JH, Kang SJ, Ryu SH, Jeon CO, Oh TK. *Hydrogenophaga bisanensis* sp. nov., isolated from wastewater of a textile dye works. Int J Syst Evol Microbiol. 2008;58:393-7.

Yurkov V, Stackebrandt E, Buss O, Vermeglio A, Gorlenko V, Beatty JT. Reorganization of the genus *Erythromicrobium*: description of “*Erythromicrobium sibiricum*” as *Sandaracinobacter sibiricus* gen. nov., sp. nov., and of “*Erythromicrobium ursincola*” as *Erythromonas ursincola* gen. nov., sp. nov. Int J Syst Evol Microbiol. 1997;47:1172-8.

**Supplementary Note 1**

Additional features of spatiotemporal distribution of genera across the *Lotus Pond*-*Rulang* ecosystem (refer to Fig. 5, and Table S5 and S6) which suggest that influx of new bacteria from the vent as well as the river, into the hydrothermal territory, increase with the progress of the day.

- At 8:30 h, only 5 genera were found to be unique to the HTP, while 38 were unique to the RVW. At 14:30 h, 29 and 27 genera were unique to the HTP and RVW respectively.
- Of the 5 genera unique to the HTP at 8:30 h, 2 were still present as unique to the HTP at 14:30 h while the other 3 were not found anywhere between HTP and RVW, and seemed to have been carried away by the hot-water flow.
- On the other hand, out of the 29 genera found as unique to the HTP at 14:30 h, 19 were not present in any of the sample-sites at 8:30 h; so they were likely to have been transported along with the sub-surface geothermal waters over the day (the other 10 however, were already there in one or more sample-sites at 8:30 h).

**These distribution patterns indicated that the vent-water’s contribution to the influx of new bacteria into the *Lotus Pond* ecosystem increases with the day.**

- Of the 38 genera unique to the RVW at 8:30 h, 27 were found in MTP and/or LTP at 14:30 h, irrespective of whether they were also found in RVW; 1 was found as unique to the HTP while 10 were not detected in any of the sample-sites, not even in RVW.

**This suggested that in the course of the day, some of the river bacteria potentially migrate into the hydrothermal territory even as some get carried away by the river current.**

- Out of the 27 genera unique to RVW at 14:30 h, 19 were nowhere there between the vent and the river at 8:30 h; 6, however, were present at HTP and/or MTP, but not at LTP or RVW; the remaining 2 were present at MTP, LTP and RVW.

**This suggested that very few river bacteria possibly migrate into the hydrothermal territory over the night.**

- Out of 31 genera found as common and restricted to LTP and RVW at 8:30 h, 23 were detected at MTP through RVW at 14:30 h, while 3 were found at HTP through RVW, 2 at MTP and RVW, and 1 each at MTP and LTP, and LTP alone (only 1 was not detected at any of the sample-sites).

**This distribution pattern buttressed the possibility of migration of copious river bacteria into the high temperature zones of *Lotus Pond*’s hydrothermal gradient over the day-time.**

- Out of 12 genera found as common and restricted to LTP and RVW at 14:30 h, 5 were present at HTP or MTP, irrespective of whether they were also found in LTP or RVW; 2 each were found as unique to LTP and RVW while 1 was not detected in any of the sample-sites.
- At 8:30 h, no genus was found as common and restricted to HTP, MTP and LTP, even though all other combinations plausible for the distribution of unique and shared genera across the sample-sites were detected. At 14:30 h, however, all plausible combinations for the distribution of unique and shared genera across the sample-sites were detected, including 8 genera common and restricted to HTP, MTP and LTP.

**This pattern was indicative of microbial transport in the vent-to-river trajectory during the day-time.**

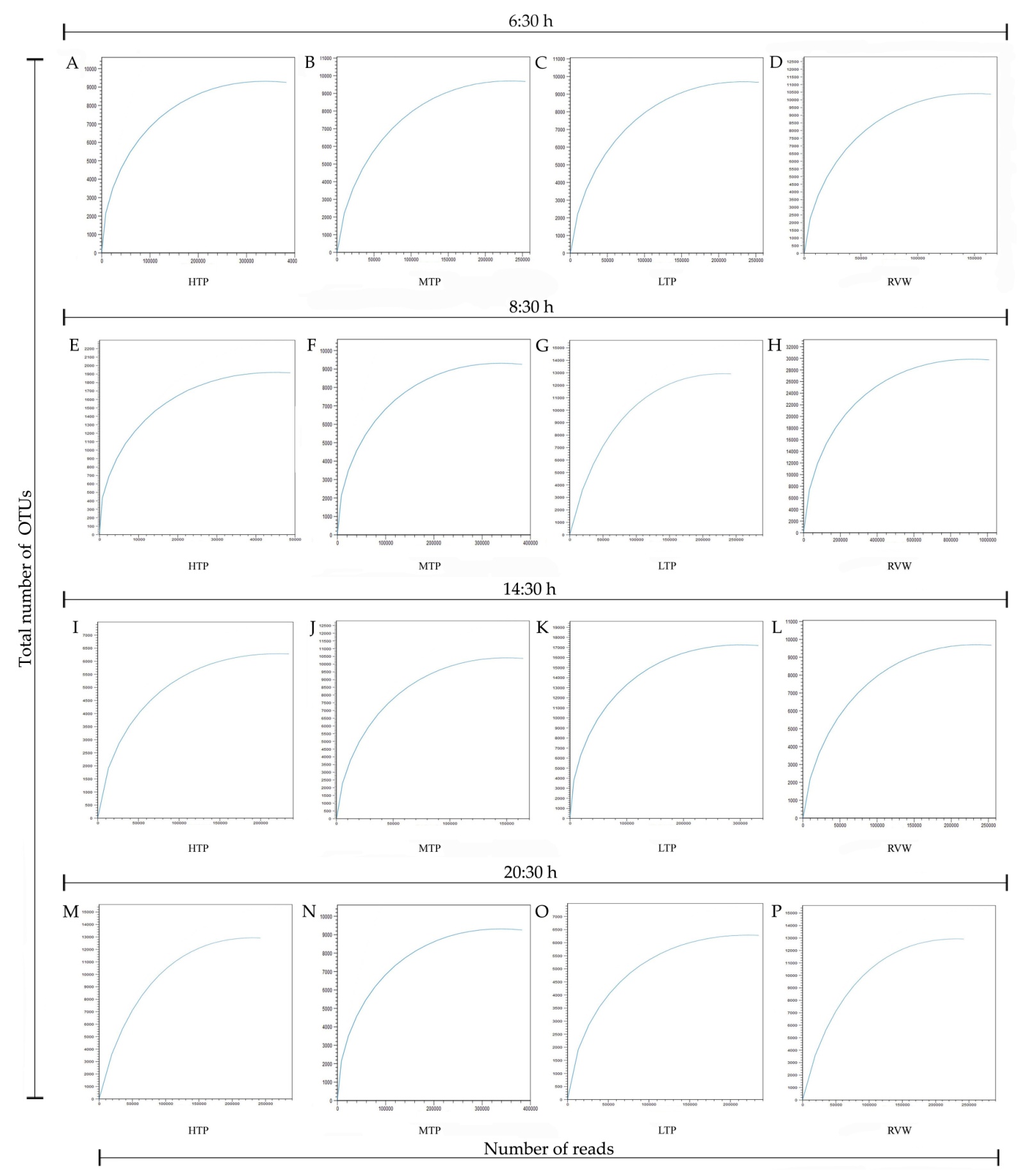

**Fig. S1.** Rarefaction curves showing the proportionality between OTU-level metataxonomic diversity revealed and the number of amplified 16S rRNA gene sequence (V3 region) reads analyzed for the individual water-samples collected from HTP, MTP, LTP and RVW separately at (**A** through **D**) 6:30 h, (**E** through **H**) 8:30 h, (**I** through **L**) 14:30 h and (**M** through **P**) 20:30 h of 20 October 2014. For each of the 16 samples, number of reads used in OTU-building are plotted along the X-axis, while number of OTUs (including singletons) created at the 97 % sequence identity level are plotted along the Y-axis.

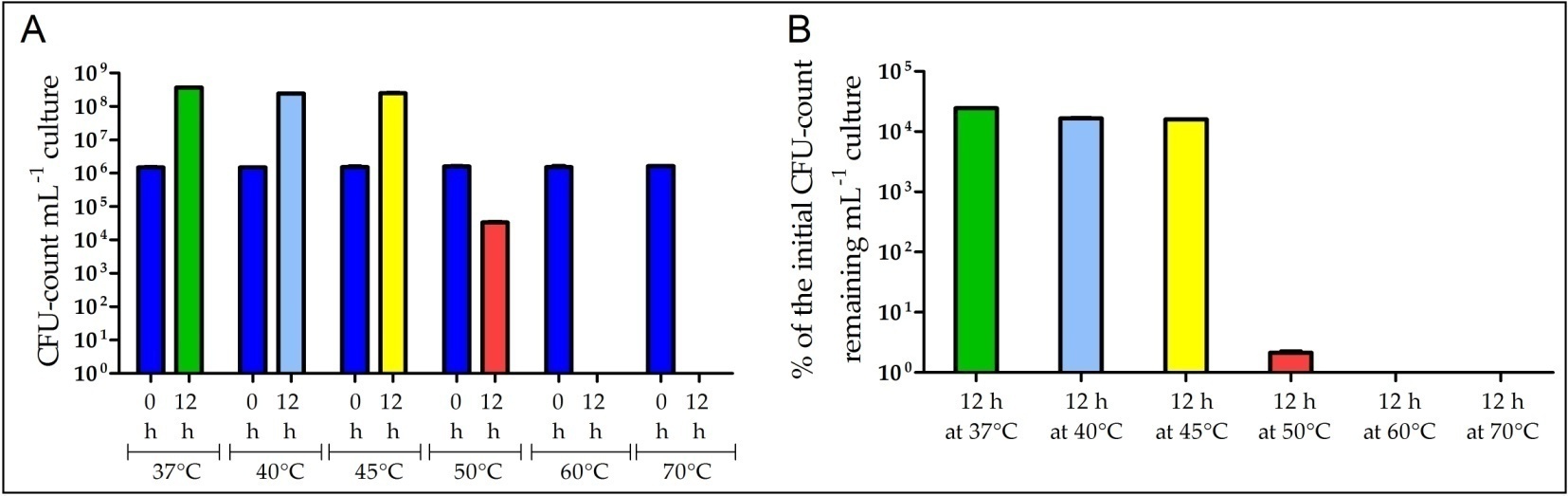

**Fig. S2.** Cellular growth yields of SMMA_5 in chemolithoautotrophic MST broth medium after 12 h incubations at 37°C, 40°C, 45°C, 50°C, 60°C and 70°C: (**A**) total number of colony-forming units (CFUs) present mL^-1^ of the broth culture at 0 h (blue bars for all the temperatures tested) and 12 h of incubation at 37°C (green bar), 40°C (light blue bar), 45°C (yellow bar), 50°C (red bar), 60°C (pink bar) or 70°C (orange bar); (**B**) percentages of the initial CFU-counts that remained in the cultures after 12 h incubations at 37°C, 40°C, 45°C, 50°C, 60°C or 70°C; temperature-wise color code for the bars in (**B**) is same as that for (**A**). All the data shown in this figure are means of the data obtained from three different experiments; error bars indicate the standard deviations.

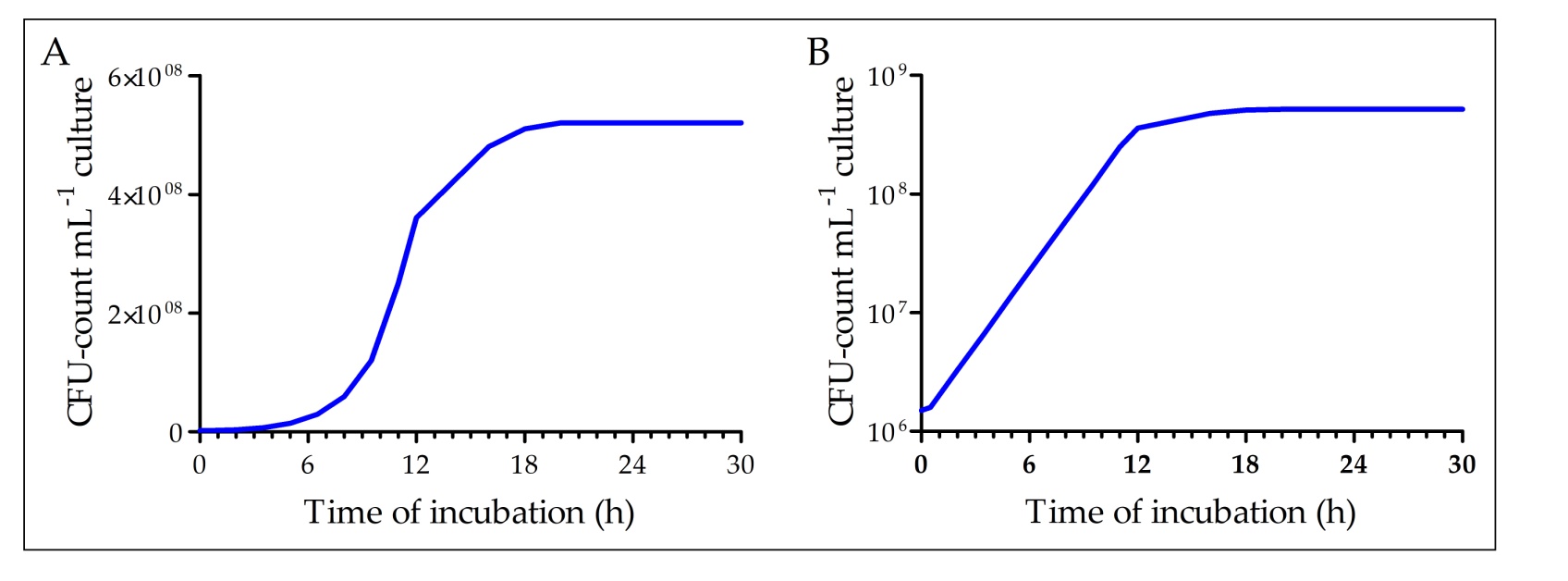

**Fig. S3.** Growth curve of *Paracoccus* sp. SMMA_5 in MST culture at 37°C: values for CFU-count mL^-1^ culture are plotted in (**A**) arithmatic and (**B**) logarithmic scales in the two different graphs.

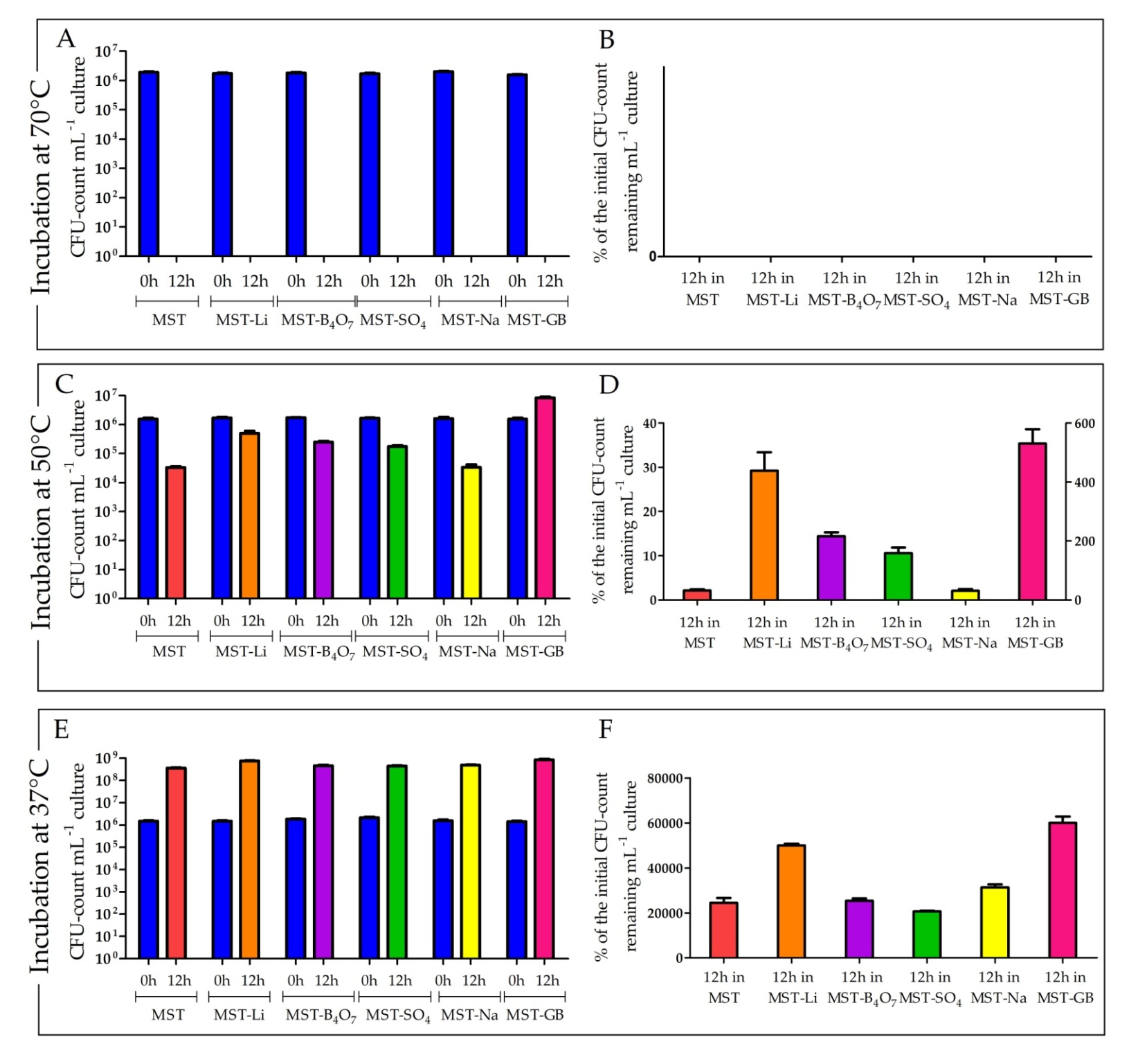

**Fig. S4.** CFU-counts recorded mL^-1^ culture at 0 h and 12 h, and percentages of the initial CFU-counts that remained ready-to-divide after incubating SMMA_5 for 12 h, in different variants of MST at 70°C, 50°C and 37°C: (**A**, **C** and **E**) CFU-counts recorded mL^-1^ culture in MST, or MST supplemented with salts of Li^+^ (MST-Li), B_4_O_7_^2-^ (MST- B_4_O_7_), SO_4_^2-^ (MST-SO_4_), Na^+^ (MST-Na) or glycine-betaine (MST-GB), at 0 and 12 hours of incubation at 70°C, 50°C and 37°C respectively; (**B**, **D** and **F**) percentages of the initial CFU-counts that remained ready-to-divide in MST, MST-Li, B_4_O_7_^2-^, MST-SO_4_, MST-Na or MST-GB, after 12 h incubation at 70°C, 50°C and 37°C respectively. In (**A**, **C** and **E**), all 0 h CFU-counts, irrespective of the test medium and incubation temperature, are represented by blue bars; CFU-count recorded after 12 h incubations in MST, MST-Li, MST- B_4_O_7_, MST-SO_4_, MST-Na or MST-GB are represented by red, orange, violate, green, yellow or pink bars respectively. In (**B**, **D** and **F**), percentages of the initial CFU-counts that remained ready-to-divide in MST, MST-Li, MST-B_4_O_7_^2-^, MST-SO_4_, MST-Na or MST-GB, after 12 h incubation, irrespective of the incubation temperature, are represented by red, orange, violate, green, yellow or pink bars respectively. In (**D**), the percentage of the initial CFU-count that remained ready-to-divide in MST-GB, after 12 h incubation at 50°C is plotted along the secondary Y-axis. All the data shown in this figure are means for three different experiments; error bars indicate the standard deviations.
